# Rapid breakdown of *Cf-6*-mediated immunity in tomato through loss or mutation of the *Avr6* effector gene in *Fulvia fulva*

**DOI:** 10.1101/2025.06.11.659170

**Authors:** Christiaan R. Schol, Like Fokkens, Anan Hu, Ruifang Jia, Silvia de la Rosa, Bilal Ökmen, Anne M. Hilgers, Samuel L. van Zwoll, Luuk D.H. Veenendaal, Marie Turner, Claudie Monot, Kazuya Maeda, Yuichiro Iida, Alex Z. Zaccaron, Ioannis Stergiopoulos, Pierre J.G.M de Wit, Danny Esselink, Anne-Marie A. Wolters, Yuling Bai, Matthieu H.A.J. Joosten, Carl H. Mesarich

## Abstract

Genetic resistance mediated by Cf cell-surface receptors is a cornerstone of tomato breeding against leaf mold disease caused by the fungal pathogen *Fulvia fulva*. After widespread deployment of the *Cf-9* resistance locus, breeders increasingly relied on *Cf-6*, yet *Cf-6*-mediated resistance has already been overcome in multiple regions. The molecular basis of this breakdown, and the identity of the matching avirulence (Avr) effector, have remained unresolved. Here, we use comparative genomics and functional genetics to identify the *F. fulva* effector Avr6 as the previously described apoplastic protein Ecp5. Sequencing of *Cf-6*-breaking strains revealed either deletion of the *Ecp5* locus or non-synonymous mutations in its coding sequence. Using transient expression, targeted gene knockout, and complementation assays, we show that *Ecp5* is both necessary and sufficient for Cf-6-mediated recognition and defence activation. We then use Avr6-triggered cell death as a phenotypic marker in a bulked segregant analysis combined with Comparative Subsequence Sets analysis (CoSSa) to map *Cf-6* to a ∼2 Mb interval on the short arm of chromosome 12 that overlaps the previously described *Cf-Ecp5.12* locus. Sequencing of *Avr6* in a broad strain collection reveals a predominantly conserved allele under apparent purifying selection, with recent independent loss-of-function variants in Europe and South America that confer virulence on *Cf-6* plants. Together, these findings establish the *Avr6–Cf-6* gene-for-gene pair, explain the erosion of *Cf-6*-mediated resistance, and illustrate how effector loss or mutation provides a rapid route for *F. fulva* adaptation, with important implications for designing more durable resistance strategies against tomato leaf mold.

## Introduction

*Fulvia fulva*, previously known as *Cladosporium fulvum*, is a Dothideomycete fungal pathogen that causes leaf mold disease of tomato (*Solanum lycopersicum*) under humid greenhouse conditions (Cooke, 1883; Thomma et al., 2005; Videira et al., 2017). It has no known sexual stage, although there is evidence of cryptic sexual reproduction (de la Rosa & Schol et al., 2024; Gül et al., 2022; Stergiopoulos, Groenewald, et al., 2007). Introgression of genetic host resistance from wild tomato relatives, mediated by cell-surface anchored, receptor-like proteins (RLPs) that recognize specific apoplastic Avirulence (Avr) effector proteins of *F. fulva* (Dixon et al., 1998; Dixon et al., 1996; Jones et al., 1994; Takken et al., 1999; Thomas et al., 1997), is an important breeding strategy against this disease. These RLPs, encoded by *Cf* (resistance to *C. fulvum*) genes, typically reside in clusters throughout the tomato genome and mediate immune responses like the hypersensitive response (HR), which is associated with a burst of reactive oxygen species (ROS) (Gust & Felix, 2014; Huang & Joosten, 2024; Kang & Yeom, 2018), the accumulation of pathogenesis-related (PR) proteins (Joosten & de Wit, 1989; Wubben et al., 1993), and the release of antimicrobial phytoalexins (de Wit & Flach, 1979). Our understanding of these RLPs and their corresponding Avr effectors, together with extensive genetic and molecular characterization of both partners, has positioned the *F. fulva*–tomato interaction as a long-standing model for cell-surface receptor-mediated immunity and effector adaptation (de Wit, 2016; Mesarich et al., 2023; Thomma et al., 2005).

Upon widespread deployment of *Cf* resistance genes in tomato cultivars, strong selection pressure is exerted on *F. fulva* to circumvent Avr recognition to regain virulence. This has so far been shown to be achieved through mutations in, or complete loss of the corresponding *Avr* gene (de la Rosa & Schol et al., 2024; Iida et al., 2015; Joosten et al., 1994; Mesarich et al., 2023; Stergiopoulos, de Kock, et al., 2007). To date, all known *F. fulva Avr* genes are highly expressed during infection and encode small proteins that are secreted into the tomato leaf apoplast (Mesarich et al., 2023). These Avr effectors are cysteine-rich and, in some cases, can evade recognition by their corresponding Cf receptor through mutation of a single cysteine residue, which disrupts an associated disulfide bond (Joosten et al., 1997; Joosten et al., 1994). Most Avr effectors of *F. fulva* show little homology to other proteins and lack known functional domains (Mesarich et al., 2018). The only exceptions are Avr4 (chitin-binding domain) and Avr2 (protease inhibitor), which protect against PR chitinases and proteases, respectively (Rooney et al., 2005; Van Esse et al., 2007). Recently, a combined transcriptomics and proteomics approach was used to show that *F. fulva* secretes at least 75 small proteins, named extracellular proteins (Ecps), into the apoplast during infection, of which many are expected to be yet-identified Avr effectors (Mesarich et al., 2018). In support of this, 10 Ecps were found to elicit an HR upon expression in various wild tomato accessions (Mesarich et al., 2018).

We recently demonstrated that *F. fulva* overcame the widely deployed *Cf-9* resistance locus by sequential loss of *Avr9* and loss or mutation of *Avr9B* (de la Rosa & Schol et al., 2024). Both of these *Avr* genes encode small apoplastic proteins and are recognized by the products of two RLP-encoding genes at the *Milky Way* (MW) locus on the short arm of chromosome 1, *Cf-9C* and *Cf-9B*, respectively (Jones et al., 1994; Parniske et al., 1997). This prompted breeders to deploy another resistance gene, *Cf-6*, which was tentatively mapped to the short arm of chromosome 12 in the breeding line Ontario 7818 and reported to originate from *S. pimpinellifolium* PI211839 (Kanwar et al., 1980). More recently, the International Seed Federation (ISF) updated the *F. fulva*–tomato differential sets used to define virulence and resistance spectra, now including five additional *F. fulva* races (9, 2.9, 4.9, 4.5.9 and 2.6.9) and designating Ontario 7818 as the differential line for *Cf-6* (Sangster, 2022). The inclusion of a race 2.6.9 strain (breaking *Cf-2*-, *Cf-6*- and *Cf-9*-mediated resistance) demonstrated that *Cf-6* provides race-specific resistance. Since then, *Cf-6*-breaking strains have been detected in multiple European countries, including Germany (Meyer & Gärber, 2021) and the Bretagne region in France, where several race 2.6.9 strains were collected between 2017 and 2021. These observations reveal that the more recently deployed *Cf-6* locus has already been overcome, with direct consequences for resistance breeding strategies.

To determine the genetic and molecular basis of *Cf-6* resistance breakdown, we set out to identify Avr6, the effector recognized by *Cf-6*-bearing tomato plants, using a comparative genomics approach. Previous efforts to identify *F. fulva Avr* genes such as *Avr5* and *Avr9B* by comparative transcriptomics and genomics compared a single resistance-breaking strain with a single non-breaking strain (de la Rosa & Schol et al., 2024; Mesarich et al., 2014). Here, we perform whole-genome sequencing and assembly of 15 *F. fulva* race 2, 2.9 and 2.6.9 strains, allowing us to systematically scan polymorphisms in small secreted protein-encoding genes, most of which had previously been characterized as Ecps (Laugé et al., 2000; Mesarich et al., 2018; Van den Ackerveken et al., 1993). We then validate the Avr6–Cf-6 interaction through PVX-mediated *in planta* expression, gene knockout, and complementation assays. Using Avr6-triggered hypersensitive cell death as a phenotypic marker, we apply bulked segregant analysis (BSA) coupled to a Comparative Subsequence Sets analysis (CoSSa) pipeline (Prodhomme et al., 2019) to map *Cf-6* in Ontario 7818. Finally, we link Avr6 recognition to resistance to *F. fulva* in segregating populations. In doing so, we identify *Avr6*, map *Cf-6* to the short arm of chromosome 12, and provide a mechanistic explanation for the breakdown of *Cf-6*-mediated resistance, while highlighting broader challenges for achieving durable resistance to tomato leaf mold.

## Materials and Methods

### Fungal and plant material

Information on the *F. fulva* strains used in this study can be found in **Table S1**. Novel French strains (sample IDs starting with FR_) were collected by VEGENOV and their virulence spectrum was determined by inoculation onto a tomato differential set carrying different *Cf* genes (including *Cf-6*), according to the ISF (Sangster, 2022). Argentinean *F. fulva* strains were previously collected and characterized by Medina et al. (2015) and Lucentini et al. (2021). Their virulence spectrum was determined by the authors using a tomato differential set (excluding *Cf-6*) and, in this study we assessed their virulence on Ontario 7818 (CGN15839; carrying *Cf-6*) and Moneymaker (MM) plants (CGN14330). Transgenic *F. fulva Avr6* knockout strains in the 0WU background were previously generated by Ökmen (2013). Strain 0WU (race 0) (de Wit et al., 2012) was used as a control strain in the inoculation studies and strain IPO 2679 (de la Rosa & Schol et al., 2024) was included as it produces Avr6^Cys30Tyr^.

For BSA-sequencing, an F2 population of 133 individuals derived from a cross between Ontario 7818 (*Cf-6*) and MM-*Cf-0* was utilized. *S. lycopersicum* Ontario 7818 (*Cf-6*), breeding line G1.1161 (*Cf-Ecp5*) (Haanstra et al., 2000) and MM-*Cf-9* (CGN15338) were utilized for PVX-mediated expression of Avr6, Avr9 and Avr9B. Detailed information regarding CGN accessions as provided by the Center of Genetic Resources Netherlands (CGN) in Wageningen, The Netherlands, can be found in **Table S2**.

### DNA isolation, whole-genome sequencing and assembly

*F. fulva* strains were grown on potato dextrose agar (PDA) at 25°C for 10–14 days, after which spores and mycelium were harvested for genomic DNA (gDNA) extraction by scraping with a sterile scalpel blade. gDNA was extracted from *F. fulva* strains FR_2_1, FR_2_2, FR_269_1, FR_269_2, FR_269_3, FR_269_4, FR_269_5, FR_29_1, FR_29_2, FR_29_3, FR_29_4, FR_29_5, FR_29_6, FR_29_7 and NZ_269_1 using a cetyltrimethylammonium bromide (CTAB) DNA protocol (Murray & Thompson, 1980) and purified using a Genomic DNA Clean & Concentrator (Zymo Research). gDNA quality and concentration were measured using the Nanodrop™ One (Thermo Scientific) and Qubit 2.0 Fluorometer (Thermo-Scientific). One hundred and fifty bp paired-end (PE150)-sequencing was performed by Novogene (https://www.novogene.com) on an Illumina NovaSeq6000 sequencing platform. Sequencing reads were trimmed with Trimmomatic v0.39 (Bolger et al., 2014) with parameters set to SLIDINGWINDOW:4:20, MINLEN:100 using Truseq2 adapter sequences, and quality of trimmed reads was checked with FASTQC (v0.12.1) (Andrews, 2019). Trimmed reads were assembled with Spades v4.0.0 (Bankevich et al., 2012) and contigs < 1 kb were removed post assembly. Assembly quality was assessed with Quast (v5.3.0) (Gurevich et al., 2013) and BUSCO (v5.7.1, genome-mode, with dothideomycetes_odb10 as reference BUSCO set and metaEUK as gene predictor) (Manni et al., 2021) (**Table S3**).

For *F. fulva* strains CF301, CF-Nangol, Can_38, 2_5, IPO_2459_60787, IPO_248911, IPO_249, Turk_1a, IMI_Argent_358077, 2_5_9 and 18-A6, gDNA was extracted using an SDS-based method (Penouilh-Suzette et al., 2020) and quality and concentration measured as above. gDNA sequencing libraries were prepared with the Invitrogen Collibri ES DNA Library Prep Kit for Illumina Systems (Thermo Fisher Scientific) according to the manufacturer’s instructions (protocol MAN001845) and sequenced (PE150) on an Illumina NovaSeq 6000 at the UC Davis Genome Center. Sequencing read quality was assessed using FastQC (v0.11.9) (Andrews, 2010) and fastp (v0.23.2) (Chen et al., 2018). Assemblies were generated using the shovill pipeline (v1.1) (https://github.com/tseemann/shovill), which merged overlapping PE reads with FLASH (v1.2.11) (Magoč & Salzberg 2011) and assembled the reads with SPAdes (v3.15.5) (Bankevich et al., 2012).

gDNA was extracted from leaves of 13-day old MM-*Cf-0*, Ontario 7818 (*Cf-6*) and Ontario 7818 x MM-*Cf-*0 F2 plants (grown under greenhouse conditions) using a cetyltrimethylammonium bromide (CTAB) DNA protocol (Murray & Thompson, 1980) and purified using a Genomic DNA Clean & Concentrator (Zymo Research). For BSA-sequencing, 200 ng of gDNA per F2 plant was pooled into the according R-or NR-pool and then purified, with gDNA quality and concentration measured as above. All 150 bp PE-sequencing of (pooled) tomato gDNA was performed by BGI (https://www.bgi.com) on a DNBseq sequencing platform.

### Compiling a dataset of public genomes and inference of phylogeny

Publicly available genome sequences were downloaded from GenBank using the query ‘*Fulvia fulva*’ (**Table S4**). The genome of *Dothistroma septosporum* NZE10 (GCA_000340195.1) was downloaded separately. We used BUSCO as described above to identify single-copy BUSCO genes in these assemblies and those generated in this study. For each BUSCO query gene, we created a fasta file with corresponding DNA sequences from each strain (33 *F. fulva* + 1 *D. septosporum*) if it was present as single-copy in all assemblies using a custom Python script. Next, we used Muscle (v 5.3.linux64) (Edgar, 2022) with default settings to infer a multiple sequence alignment for each fasta and trimmed alignments with TrimAl (v1.5.rev0) (Capella-Gutiérrez et al., 2009) with settings optimized for maximum likelihood inference (-automated1). Trimmed alignments were then concatenated with a second custom Python script into a single alignment with 6917 parsimony informative sites. We used IQtree (Minh et al., 2020) with ModelFinder (Kalyaanamoorthy et al., 2017) and UFBoot (-m MFP -B 1000 - bnni -alrt 1000) (Hoang et al., 2018) to identify an optimal substitution model (GTR+F+I+R10) and infer a consensus maximum likelihood phylogeny from this concatenated alignment with 1000 bootstrap replicates. For the figures, the outgroup (*D. septosporum* NZE10) and isolate CBS131901 (short-read assembly of 0WU) were removed. All custom scripts can be downloaded from https://git.wur.nl/like.fokkens/clado_avr6.

### Identification of *MAT* loci and sequence comparison of known and predicted effector genes

To infer mating types of *F. fulva* strains, we used megablast (v 2.13.0+) (Morgulis et al., 2008) with default settings to align *Mat1-1* and *Mat1-2* sequences (accessions DQ659350.2 and DQ659351.2 respectively). For identification of effector genes in the assemblies, we used a custom python script implementing bedtools getfasta (v2.30.0) (Quinlan & Hall, 2010) and a list of predicted effector genes from Zaccaron et al. (2022) to obtain a fasta file with sequences from the assembly of the Race 5 reference strain (GCA_020509005.2) including introns. We then used megaBLAST (v 2.13.0+) with – default settings to search for homologs of each of these genes. To obtain a similarity score for each strain/assembly and each effector gene, relative to the Race 5 reference, we divided the maximum Smith-Waterman score of the effector gene against the assembly with its maximum score against the assembly of Race 5. We used a custom Python script to parse the BLAST output table and calculate these scores. For known *Avr* genes we generated MSAs to identify distinct genotypes. To confirm presence/absence of predicted effectors and to determine the size of the deletion resulting in loss of *Avr6*, we used Bowtie2 (v2.5.4, with -I 100 -X 1000) (Langmead & Salzberg, 2012) to align reads to the assembly of the Race 5 strain, and checked the regions where effectors were predicted to be absent in IGV (Robinson et al., 2011). For some strains, certain effectors were not present in the assembly, while we did find reads mapped to the gene in the Race 5 genome, which is indicated in **Figure S1**. We used custom R scripts to plot the phylogeny with different *Avr* genotypes (**Figure 1**), and with a heatmap with effector gene sequence similarity (**Figure S1**).

**Figure 1:**
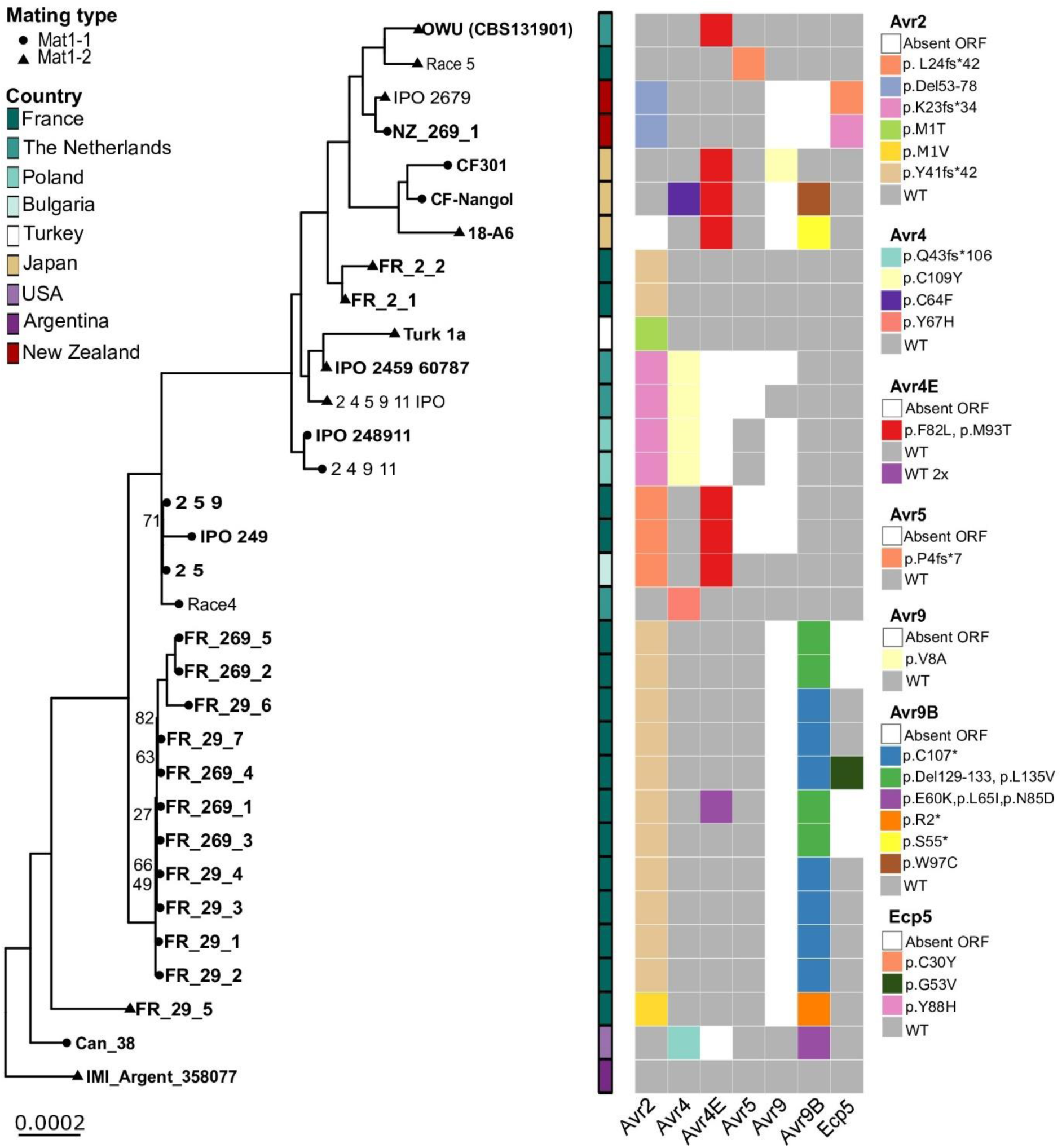
Phylogeny and predicted amino acid sequence comparisons of known Avrs, and Ecp5, across a panel of 32 *F. fulva* genomes. Maximum-likelihood phylogeny inferred from a concatenated alignment of 3484 BUSCO genes, rooted with *Dothistroma septosporum* as outgroup (not shown in the tree). Only branches with bootstrap values < 95 are shown in the tree. Sequences generated for this study are shown in bold. Mating types are indicated with symbols at the tips of the tree (circles and triangles), and the country of origin is indicated with colored rectangles. Distinct predicted amino acid sequence mutations in known Avr proteins Avr2, Avr4, Avr4E, Avr5, Avr9 and Avr9B, and in Ecp5 are indicated with squares in different colors in the right panel, with wild-type (WT) sequences indicated in grey, and absence of an open reading frame (ORF) indicated in white. p., protein sequence numbering; del, deletion; fs, frame shift; *, stop codon.

### Bioinformatics pipeline for BSA-sequencing by means of Comparative Subsequence Sets analysis (CoSSA)

A Comparative Subsequence Sets Analysis (CoSSA) bioinformatics pipeline was applied to find the resistance-specific haplotype (Prodhomme et al., 2019). Sequence reads were trimmed and sequencing adaptors were removed using fastp (version 0.23.4) (Chen, 2023) (**Table S5**). *K*-mer Counter (KMC) and KMC tools (version 3.2.4) were used to produce *k-*mer (*k*=31) tables from the trimmed reads, and *k*-mer table subtraction, intersection and coverage-based filtering (Kokot et al., 2017). GenomeScope 2.0 (Ranallo-Benavidez et al., 2020) was used to visualize the *k-*mer coverage profiles. The NR-bulk specific *k-*mer table was subtracted from the R-bulk specific table, and a *k*-mer table of the susceptible parent (MM-*Cf-0*) was subtracted from the resulting R-bulk specific *k*-mers, and intersected by a *k*-mer table of the resistant parent (Ontario 7818). *K*-mer sequences with coverage outside a range of 26 to 43 were removed to reduce error-to-noise ratio. Cutoff values are calculated as 25% margins of the expected coverage of the *R* haplotype. Filtered *k-*mers were mapped to the SL4.00 tomato reference genome (Hosmani et al., 2019) using BWA aln (version 0.7.18) (Li & Durbin, 2009) and plotted based on genomic location and coverage. Used scripts reflect the workflow of (Prodhomme et al., 2019) and are available on Github (https://github.com/PBR/CoSSA). Structural predictions of Cf-6 candidate proteins were performed using predicted protein sequences from the ITAG4.0 annotation of the Heinz 1706 SL4.0 genome using InterPro (Paysan-Lafosse et al., 2023).

### Homolog searches and AlphaFold2 structural predictions

The Basic Local Alignment Search Tool for nucleotide (BLASTn), translated nucleotide (tBLASTn), and protein (BLASTp) were used to identify *Avr6-/*Avr6-like sequences in the National Centre for Biotechnology Information (NCBI) and Joint Genome Institute (JGI) Mycocosm sequence databases. (Laugé et al., 2000) Signal peptides were predicted using SignalP v4.1 (Nielsen, 2017). The ColabFold (Mirdita et al., 2022) implementation of AlphaFold2 (Jumper et al., 2021) was used to predict the tertiary structure of Avr6 (without its endogenous signal peptide) and was assisted through use of a custom MSA generated with Avr6-like proteins, using Geneious v9.1.8 (Kearse et al., 2012). Foldseek (Van Kempen et al., 2024) was used to identify structural homologs of Avr6 in filamentous fungi in the RCSB Protein Data Bank. The predicted tertiary structure of Avr6, along with those of proteins identified using Foldseek, were visualized, rendered, and aligned (CEalign plugin) using PyMOL (DeLano, 2002).

### *F. fulva* disease assays

*F. fulva* strains were inoculated from glycerol stocks on PDA plates supplemented with 1.5% extra technical agar and 33 mg/L chloramphenicol and grown for approximately 2–3 weeks at room temperature in the dark. Then, conidia were harvested and suspended in tap water to a concentration of approximately 1×10^6^ conidia/mL. This suspension was then spray-inoculated on the abaxial side of 3- to 4-week-old tomato plants which were subsequently kept under high (90–100%) humidity in a plastic tent for 3 days, after which the tent was opened. Plants were maintained in a greenhouse compartment set to 21–23°C. Disease symptoms were scored at 14 days post-inoculation (dpi).

### PVX-mediated expression assays

Systemic PVX-based expression of Avr effectors was performed by means of agroinfiltration of the cotyledons of 10-day (for the validation of Avr6) or 16-day (for BSA-sequencing of *Cf-6*) old tomato seedlings, as previously described by Mesarich et al. (2018) and de la Rosa & Schol et al. (2024). The *Agrobacterium* strain GV3101 carrying *pSfinx::Ecp5* was previously generated by Laugé et al. (2000) and the strain carrying *pSfinx::Avr9* by Hammond-Kosack et al. (1995). Symptoms were scored at 10 dpi.

### *F. fulva* gene complementation

Complementation of strains with the wild-type *Avr6* gene was performed by means of *Agrobacterium*-mediated transformation using the same method and construct as described in (de la Rosa & Schol et al., 2024).

### *Avr6* allelic variation analysis

*Avr6* was amplified from the genomic DNA of 90 *F. fulva* strains using primers Avr6 Fw2 (5’-CTGACATAGCATTACACGAGCAG-3’) and Avr6 Rev (5’-CGCGTCGCCTGATAGATTTG-3) by PCR, as previously described for the *Avr9B* gene (de la Rosa & Schol et al., 2024). Amplicons were purified using ExoSAP-IT™ PCR Product Cleanup Reagent (Thermo Scientific) and sequenced by Macrogen™ Europe (https://www.macrogeneurope.com) by means of Sanger Sequencing using an ABI 3730xl DNA analyzer (Applied Biosystems). Consensus sequences were generated and imported in Benchling (Biology Software, https://benchling.com) for end-trimming and sequence alignment to the *Ecp5* coding sequence (CLAFUR5_03423) from the *F. fulva* strain Race 5 reference genome (Zaccaron et al., 2022).

## Results

### Deletion or mutation of *Extracellular protein 5* (*Ecp5*) is associated with *Cf-6*-breaking *F. fulva* strains

As a starting point for identifying *Avr6*, we obtained 14 French *F. fulva* strains that were sampled from diseased tomato plants in the French region of Brittany between 2017 and 2021, of which five were reported to break *Cf-6,* and a strain from New Zealand that was reported to break *Cf-6* in 2012. We confirmed that six of these strains indeed break *Cf-6*-mediated resistance with a disease assay on MM-*Cf-0* and Ontario 7818 (*Cf-6*) tomato plants (**Figure S2**). To further expand the allelic diversity of candidate effectors, we also obtained 11 additional *F. fulva* strains from diverse locations around the world (Stergiopoulos et al., 2007). Subsequently, all 26 *F. fulva* strains were subjected to Illumina sequencing to generate whole-genome assemblies (**Table S3**) and we compiled these with publicly available *F. fulva* assemblies (**Table S4**) into a dataset of 32 genomes. Then, we inferred a phylogeny based on a concatenated alignment of 3484 conserved, single-copy orthologs and found that the strains fall into four distinct clades, with most (11) clustering into a single, closely related clade (**Figure 1**). Then, we inferred mating types for all strains in the dataset and found that 11 had *Mat1-1* (36.4%) and 21 had *Mat1-2* (63.6%) at their mating type (MAT) locus. Further inspection of the distribution of genotypes for known *Avr* genes *Avr2*, *Avr4*, *Avr4E*, *Avr5*, *Avr9* and *Avr9B* revealed that predicted mutations in Avr2 (p.del54-78, p.Tyr41fs*34, p.Lys23fs*42), Avr4 (p.Cys109*) and Avr4E (p.Phe82Leu and p.Met93Thr) occur in both mating types, which suggests occurrence of recombination and thus sexual reproduction within *F. fulva* populations.

Next, we queried these genomes to identify candidate genes for *Avr6*, i.e. putative effector genes that have non-synonymous mutations or are (partially) absent in all race 2.6.9 strains but not in others. Hypothesizing that *Avr6* should be an small apoplastic protein, we used BLAST to compare the sequences of a set of known and predicted *Ecp* genes from the *F. fulva* race 5 reference strain (Mesarich et al., 2018; Zaccaron et al., 2022) to our dataset, and searched for those that were absent or exclusively had one or more mutations in the race 2.6.9 strains (**Figure S1**). As previously reported/predicted, the *Ecp* genes, although displaying some minor sequence variation, are present in most or all strains (Mesarich et al., 2018; Stergiopoulos, de Kock, et al., 2007), with *Ecp24-2* and *Ecp5* being notable exceptions. Strikingly, we found that *Ecp5* (Laugé et al., 2000) was deleted in four of the six *Cf-6*-breaking strains (**Figure 1**, **Figure S1**). In these strains, we observed a deletion of approximately 18 kb on chromosome 2, resulting in the loss of *Ecp5* (*CLAFUR5_03423*) and three adjacent genes, i.e. *CLAFUR5_20157*, *CLAFUR5_03424*, and *CLAFUR5_03425* (**Figure S3**). However, these three genes are predicted to code for uncharacterized, non-secreted proteins and are therefore unlikely candidates for *Avr6*. The deletion of the 18 kb locus was likely mediated by transposable elements (TEs) that, as previously shown, surround *Ecp5* (de Wit et al., 2012). In the other two *Cf-6*-breaking strains, non-synonymous mutations in *Ecp5*, resulting in amino acid substitutions p.Tyr88His and p.Gly53Val, were observed. Strain IPO 2679, which was used for the identification of *Avr9B* (de la Rosa & Schol et al., 2024), was the only other strain showing sequence variation for *Ecp5*, resulting in a p.Cys30Tyr substitution. In all other strains, a wild-type *Ecp5* allele was present. For *Ecp24-2*, we did not observe a clear correlation with the ability to overcome *Cf-6* (**Figure S1**) and it was therefore omitted as a candidate for *Avr6*. Together, these results suggest that *Ecp5* is *Avr6*.

### Molecular validation confirms that Ecp5 is Avr6

To confirm that Ecp5 is Avr6, we systemically expressed it in *Cf-6* tomato (Ontario 7818) and *S. lycopersicum* G1.1161, the line carrying the *Cf-Ecp5* locus that was mapped to the short arm of chromosome 1 (Haanstra et al., 2000). Here, potato virus X (PVX)-based expression was employed, whereby plants are transiently transformed with a recombinant version of PVX containing *Ecp5* (PVX::*Ecp5*) using *Agrobacterium tumefaciens* (Haanstra et al., 2000; Laugé et al., 2000). As controls, *Ecp5* was also expressed in MM-*Cf-0* plants that are universally susceptible to *F. fulva* and in MM-*Cf-9* plants carrying the *Cf-9* locus. Concomitant with the systemic movement of the recombinant virus in the plant, Ecp5 is produced systemically. Should Ecp5 be recognized by Cf-6, a systemic, visible HR, resulting in chlorosis, necrosis, stunting, or death of the plant is expected to occur (Haanstra et al., 2000; Laugé et al., 2000).

An HR developed in Ontario 7818 and G1.1161 plants, but not in MM-*Cf-0* and MM-*Cf-9* plants, supporting that Ecp5 is Avr6 (**Figure 2**). To further validate that *Cf-6*-mediated resistance is solely based on Ecp5 recognition, we inoculated MM-*Cf-0* and Ontario 7818 plants with the wild-type race 0 *F. fulva* strain 0WU that produces Ecp5 (de Wit et al., 2012), and two independent *Ecp5* knockout strains in this background, generated previously by Ökmen (2013). As expected, the wild-type 0WU strain was virulent on MM-*Cf-0* plants, but avirulent on Ontario 7818 plants, whereas the two *Ecp5* knockout strains both had gained virulence on Ontario 7818 plants, with symptoms visually indistinguishable from those caused by the wild-type 0WU strain on *Cf-0* plants (**Figure 3**). This confirms that only Ecp5 is recognized by *Cf-6* tomato plants to mediate resistance to *F. fulva.* From here on, we will refer to Ecp5 as Avr6.

**Figure 2:**
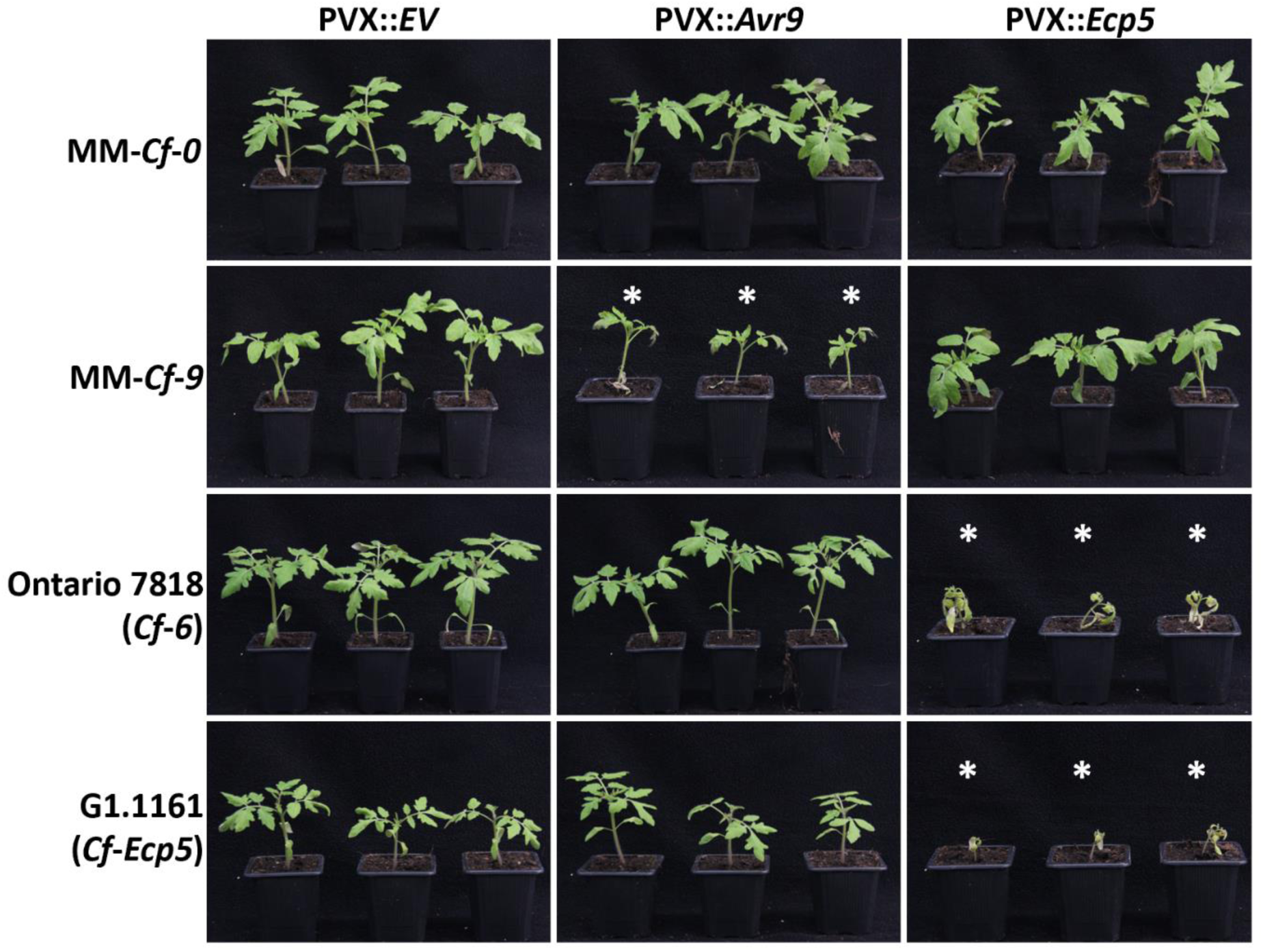
Ecp5 elicits a systemic HR in young tomato plants carrying *Cf-6* and *Cf-Ecp5*. Ecp5 was systemically expressed in seedlings of MM-*Cf-0* (no *Cf* genes), MM-*Cf-9* (*Cf-9B* and *Cf-9C*), Ontario 7818 (*Cf-6*), and G1.1161 (*Cf-Ecp5*), using the PVX-based expression system, with three replicates per tomato line. The recombinant virus PVX::*EV* (empty vector) was used as a negative control, while the recombinant virus PVX::*Avr9* was used as positive control. All recombinant viruses were delivered by *A*. *tumefaciens*-mediated transient transformation assays, through infiltration of the cotyledons of 10-day-old seedlings. Photographs were taken 10 days post agro-infiltration. White asterisks indicate plants undergoing systemic HR.

**Figure 3:**
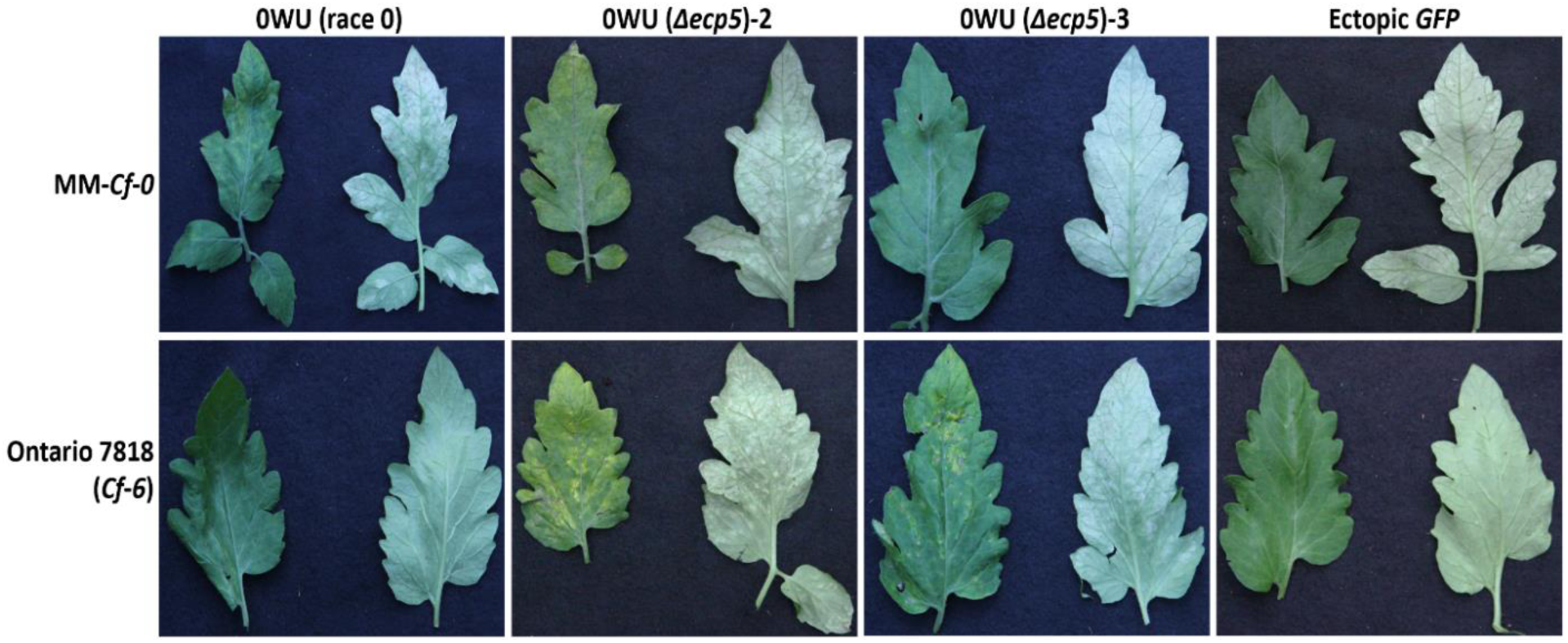
Deletion of *Ecp5* in the *F. fulva* race 0 strain 0WU confers virulence on Ontario 7818 (*Cf-6*) tomato plants. The 0WU wild-type strain, two independent *F. fulva Ecp5* knockout strains (*Δecp5*-2 and *Δecp5*-3) and an ectopic *green fluorescent protein* (*GFP*) transformant control were inoculated on 4-week-old Ontario 7818 (*Cf-6*) and MM-*Cf-0* (no *Cf* genes) tomato plants. The upper and lower surfaces of leaflets were photographed at 14 days post-inoculation.

### Polymorphisms in *Avr6* allows *F. fulva* to overcome *Cf-6*-mediated resistance

Next, we validated that polymorphisms in Avr6 observed in race 2.6.9 strains allow circumvention of *Cf-6*-mediated resistance. *F. fulva* strain IPO 2679 carries a p.Cys30Tyr mutation in Avr6, which we hypothesized would allow it to break *Cf-6*-mediated resistance. To test this, we inoculated transgenic IPO 2679 strains complemented with the wild-type *Avr6* gene from strain 0WU, generated by de la Rosa & Schol et al. (2024), onto MM-*Cf-0* and Ontario 7818 (*Cf-6*) plants. We observed that the wild-type IPO 2679 strain is virulent on both lines, whereas the two complemented transformants have lost the ability to infect Ontario 7818 (*Cf-6*), but not MM-*Cf-0* plants (**Figure 4A**). Similarly, when strains FR_269_1 (del*Avr6*) and NZ_269_1 (p.Tyr88His) were complemented with wild-type *Avr6*, virulence was abolished on *Cf-6* plants but not on MM-*Cf-0* (**Figure 4B**). We conclude that these three mutations in Avr6 each allow the fungus to break *Cf-6*-mediated resistance. This contrasts with previous results, in which no allelic variation had been described in *Avr6* across a large core collection of *F. fulva* strains (Iida et al., 2015; Stergiopoulos, de Kock, et al., 2007). To determine whether there is evidence for further selection pressure on *Avr6* in response to *Cf-6* deployment, we sequenced *Avr6* in 155 additional *F. fulva* strains that were predominantly collected over the past two decades and mostly originate from Europe and Japan. A wild-type *Avr6* gene was present in almost all strains (**Table S1**), suggesting that these are unable to cause disease in *Cf-6* plants. Only in one recently collected German strain (P3), we identified a predicted splice site mutation at the end of the second intron of *Avr6* which resulted in (partial) restoration of virulence on *Cf-6* plants (**Figure S4**). Although the observed symptoms were not as pronounced as on MM-*Cf-0*, they were more severe compared to the symptoms caused by strain P30, which has a wild-type *Avr6* gene, indicating that this mutation impairs the recognition of Avr6 by Cf-6. This partial restoration of virulence is anticipated to be due to a reduction in the amount of correctly spliced *Avr6* transcript and, thus, Avr6 protein available for recognition by Cf-6. In all 154 other strains, we found a wild-type *Avr6* gene, suggesting that *Cf-6*-breaking strains recently emerged.

**Figure 4:**
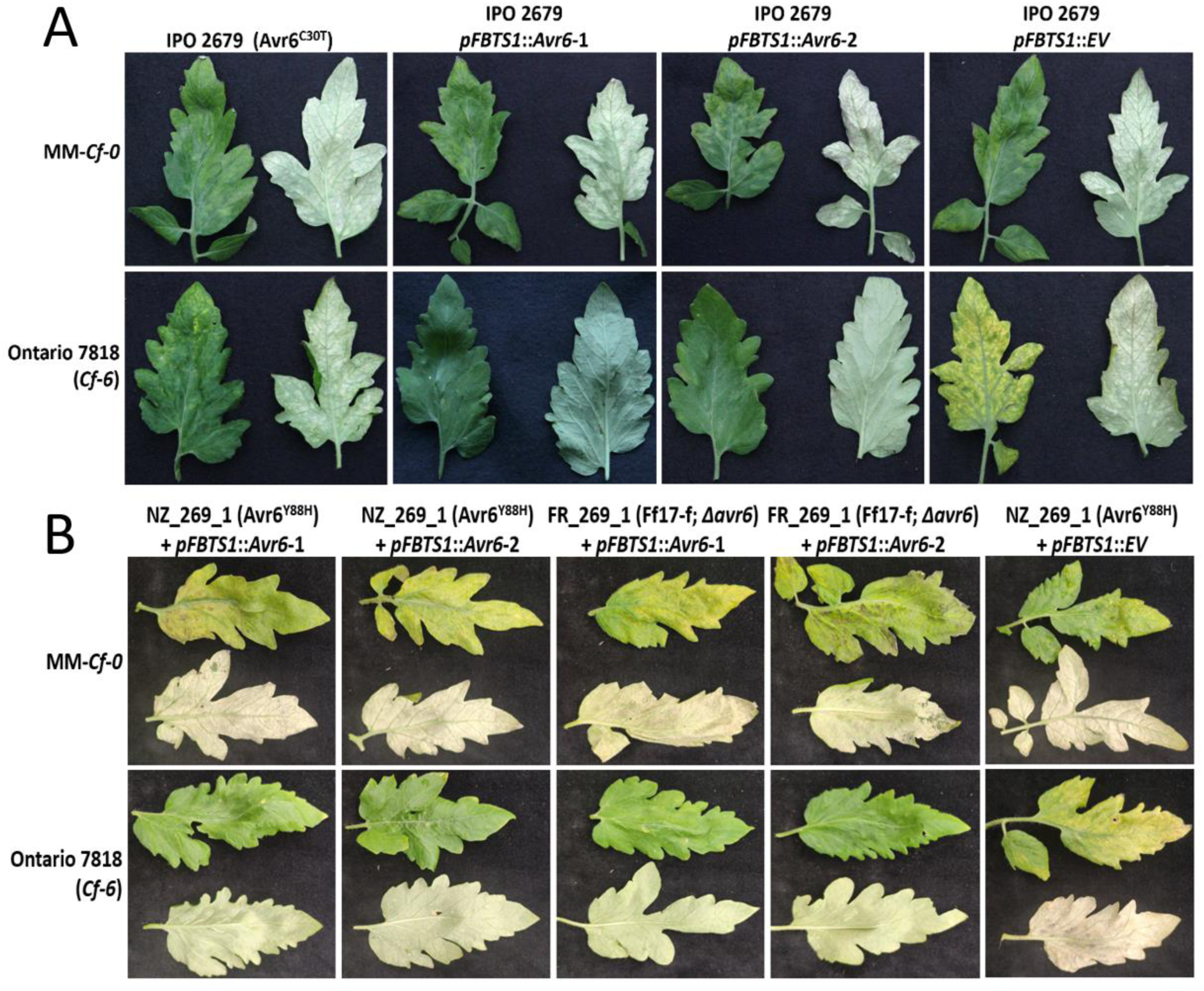
Complementation of *Cf-6*-breaking *F. fulva* strains with wild-type *Avr6* from strain 0WU results in loss of virulence on Ontario 7818 (*Cf-6*) tomato plants. The IPO 2679 (Avr6, p.Cys30Tyr) wild-type strain, two independently generated IPO 2679 *Avr6*-complemented *F. fulva* (*Avr6*-1 and *Avr6*-2) transformants, and an empty vector (EV) control transformant were inoculated on 4-week-old Ontario 7818 (*Cf-6*) and MM-*Cf-0* (no *Cf* genes) tomato plants (Panel A). Two independently generated NZ_269_1 and FR_269_1 *Avr6*-complemented *F. fulva* (*Avr6*-1 and *Avr6*-2) transformants, and an EV control, were inoculated on 4-week-old Ontario 7818 (*Cf-6*) and MM-*Cf-0* (no *Cf* genes) tomato plants (Panel B). Leaflets were photographed at 14 days post-inoculation.

Recently, however, non-synonymous and frame-shift mutations were reported in *Avr6* of several *F. fulva* strains collected in Argentina (Lucentini et al., 2021; Medina et al., 2015). To investigate whether these mutations also result in virulence on *Cf-6* tomato, we inoculated strains expressing Avr6 carrying the mutations p.Lys22*, p.Cys30fs*74, p.Val40fs*101 and p.Val98fs*104 (**Table S1**) onto MM-*Cf-0* and Ontario 7818 plants. This resulted in visible disease symptoms on both genotypes for strains harboring these mutant alleles, but not for the control strains 0WU or CIDEFI 305, which both contain wild-type *Avr6* and are therefore unable to overcome *Cf-6* (**Figure 5**). As all mutations are predicted to result in either frameshifts or cysteine mutations, it is not surprising that recognition by Cf-6 is abolished. Unfortunately, we could not test strain CIDEFI 318 (p.Cys48Tyr) due to its poor growth and sporulation *in vitro*, but it is likely that the p.Cys48Tyr mutation also abolishes recognition. In all, apart from deletion of the *Avr6* ORF and the splice site mutation in P31, we identified eight different non-synonymous and frame-shift mutations in *Avr6* across strains (**Figure 6**), of which seven were confirmed to result in breakdown of *Cf-6* by *F. fulva*.

**Figure 5:**
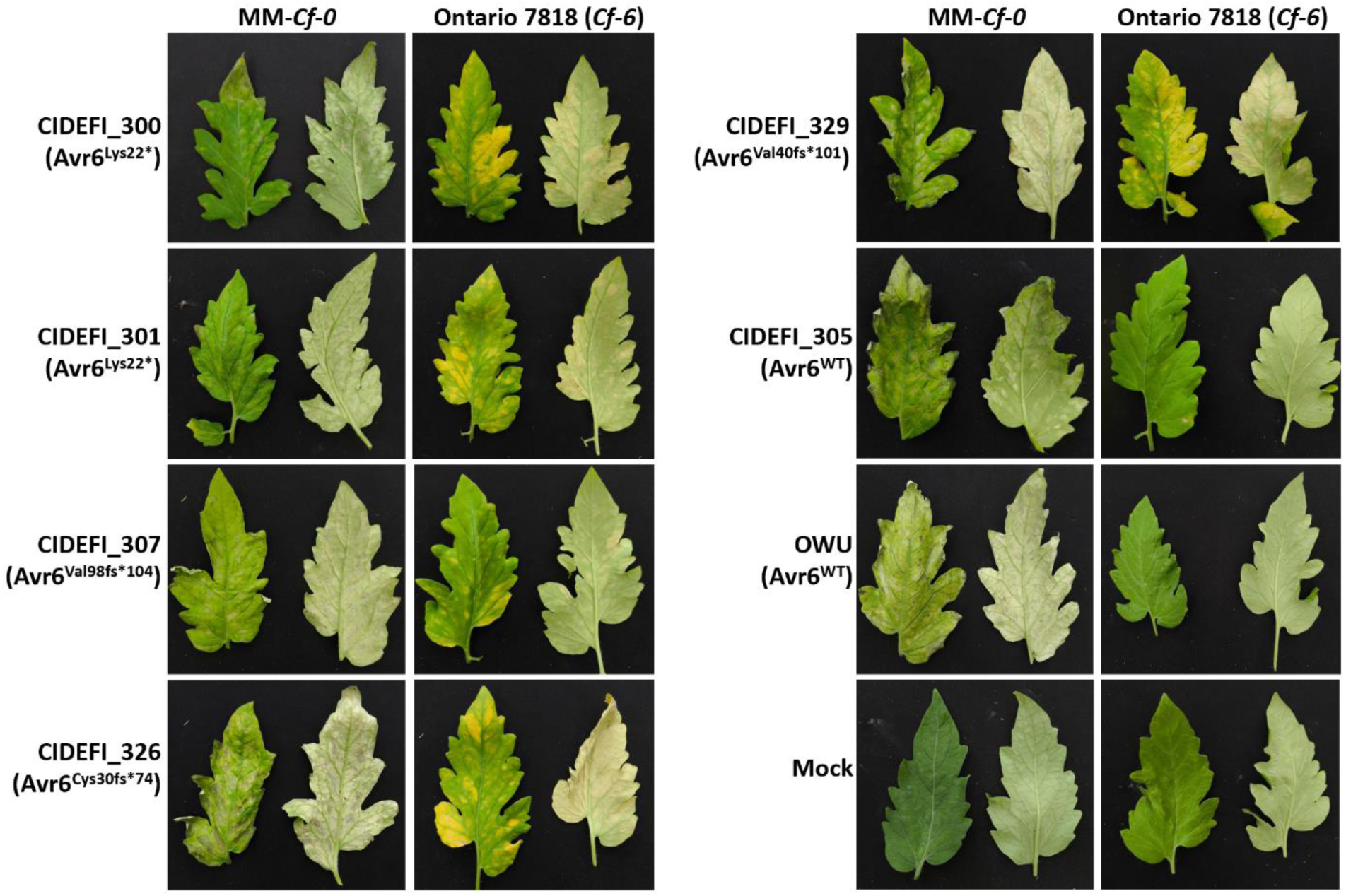
Non-synonymous and frame-shift mutations in *Avr6* of Argentinian *F. fulva* strains confer virulence on *Cf-6* tomato plants. Four-week-old MM-*Cf-0* (no *Cf* genes) and Ontario 7818 (*Cf-6*) tomato plants were inoculated with several of the Argentinian *F. fulva* strains reported by Medina et al. (2015) and Lucentini et al. (2021) to harbor mutations in *Avr6* (race 2.6 (initially reported as race 2): CIDEFI 300, 301, 307 and 329; race 6 (initially reported as race 0): CIDEFI 326). As controls, strain 0WU (race 0) and CIDEFI 305 (race 2), which can infect MM-*Cf-0* but not *Cf-6* plants, were also inoculated. Symptoms were photographed at 14 days post-inoculation.

**Figure 6:**
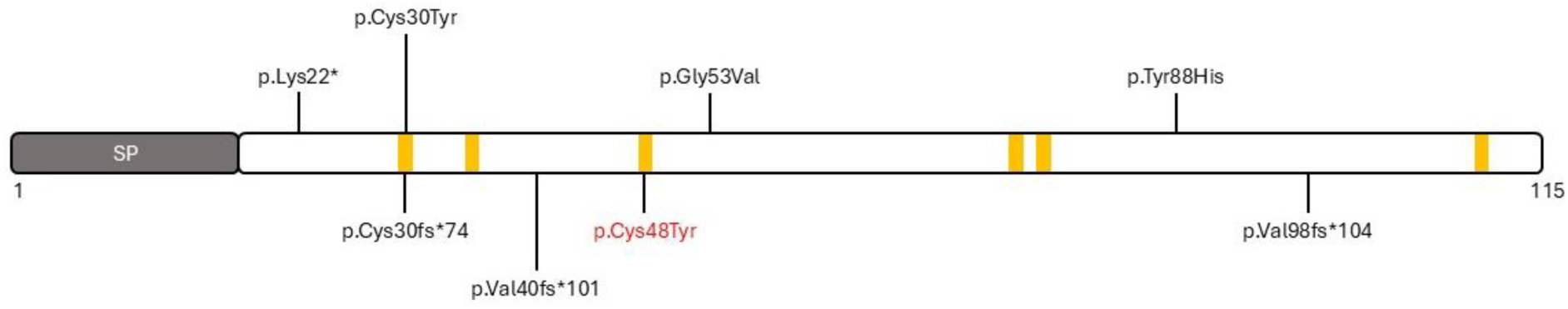
Schematic representation of the Avr6 protein showing the location of mutations that allow *F. fulva* to overcome *Cf-6*-mediated resistance. The signal peptide (SP) is indicated in grey and yellow bars indicate the positions of cysteine residues. The strain carrying the predicted p.Cys48Tyr mutation (in red font) could not be tested.

### Many plant-pathogenic Dothideomycete and Sordariomycete fungi are predicted to produce an Avr6-like protein

To investigate whether other fungi produce Avr6-like proteins, BLAST searches were performed against fungal genomes (tBLASTn) and their associated protein catalogues (BLASTp) in publicly available sequence databases using *F. fulva* Avr6 as a query. This resulted in the identification of several Avr6-like proteins, all from plant-pathogenic Dothideomycetes and Sordariomycetes (**Figure S5**). One such protein was identified in *Pseudocercospora fuligena* (**Figure S5**), a Dothideomycete responsible for black leaf mold disease of tomato (Phengsintham et al., 1938; Zaccaron and Stergiopoulos, 2020). All identified Avr6-like proteins were predicted to possess an N-terminal signal peptide for extracellular targeting (**Notes S1**). Furthermore, based on a sequence alignment, the six cysteine residues present in Avr6 were conserved in all Avr6-like proteins (**Figure S5**). Aside from these conserved cysteine residues, several other amino acid residues were highly conserved, including Tyr88 and Gly53, which were mutated in *F. fulva* strains NZ_269_1 (p.Tyr88His) and FR_269_4 (p.Gly53Val), indicating that they may play a crucial role in the overall structure of Avr6 and/or its role in virulence (**Figure S5**).

### The predicted tertiary structure of Avr6 shows similarity to Leptosphaeria AviRulence and Suppressing (LARS) effectors

To gain insight into possible functions and constraints on Avr6, we predicted its tertiary structure (without the signal peptide) using AlphaFold2 and compared it to experimentally determined fungal structures in the RCSB Protein Data Bank using Foldseek. The predicted structure consists of six β-strands and one α-helix, stabilized by three putative disulfide bonds (Cys13-Cys18, Cys31-Cys59 and Cys61-Cys94), consistent with a compact, cysteine-rich fold (**Figure 7**). One predicted disulfide bond (Cys61–Cys94) was not explicitly called by AlphaFold2 but is strongly suggested by the close spatial proximity of the two residues. Mapping the positions of the Gly53 and Tyr88 substitutions found in *Cf-6*-breaking strains indicates that Gly53 lies in a surface-exposed loop linking the sole α-helix to β-strand 2, whereas Tyr88 is predicted to be buried, suggesting that these substitutions are likely to affect protein stability and/or interaction surfaces in distinct ways.

**Figure 7:**
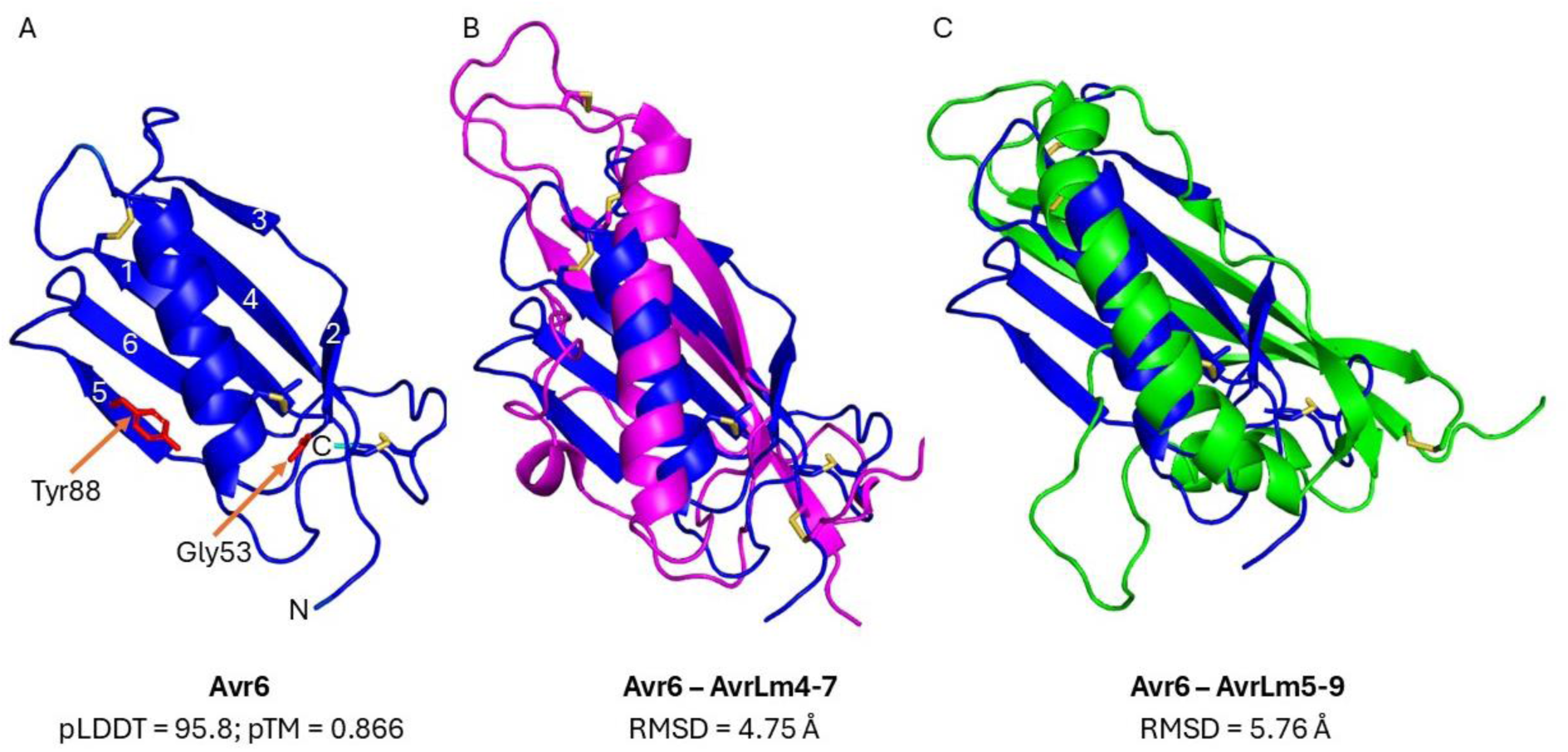
Avr6 is predicted to adopt a LARS-like fold. **A.** Predicted tertiary structure of Avr6. The tertiary structure is colored according to the predicted Local Distance Difference Test (pLDDT) score (confidence: cyan = high; dark blue = very high). Gly53 and Tyr88 are shown as sticks and are colored red. β-strands are numbered sequentially. Predicted disulfide bonds are colored yellow. N- and C-termini are shown. **B.** Alignment of Avr6 (blue) with AvrLm4-7 (magenta; RCSB Protein Data Bank (PDB) ID: 4FPR) from *L. maculans.* **C.** Alignment of Avr6 (blue) with AvrLm5-9 (green; PDB ID: 7B76) from *L. maculans*. pTM: predicted Template Modelling score. RMSD: Root Mean Square Deviation.

Foldseek analysis identified the Leptosphaeria AviRulence and Suppressing (LARS) effectors AvrLm4-7 (PDB ID: 4FPR) and AvrLm5-9 (PDB ID: 7B76) from *Leptosphaeria maculans* (Blondeau et al., 2015; Lazar et al., 2022), as the closest structural matches to Avr6, with moderate similarity (TM-scores ∼0.55) (**Figure 7**). These proteins share an overall β-sheet–rich architecture but differ in elongation and loop topology, and exhibit limited primary sequence similarity. Thus, Avr6 appears to adopt a LARS-like fold, although it forms a distinct structural variant within this effector family. The biological functions and host targets of AvrLm4-7 and AvrLm5-9 remain unknown, and further biochemical work will be needed to determine whether Avr6 shares mechanistic features with these effectors or uses this fold for entirely different virulence activities.

### Bulked segregant analysis maps *Cf-6* in Ontario 7818 to a 2 Mb interval on the short arm of chromosome 12

Currently, three distinct loci originating from different *S. pimpinellifolium* accessions that mediate a gene-for-gene relationship with *Avr6* have been reported, i.e. *Cf-Ecp5.1* on the short arm of chromosome 1 (Haanstra et al., 2000; Iakovidis et al., 2020), *Cf-Ecp5.7* on the long arm of chromosome 7, and *Cf-Ecp5.12* on the short arm of chromosome 12 (Iakovidis et al., 2020). To validate the location of *Cf-6* on the short arm of chromosome 12 in Ontario 7818 as reported by Kanwar et al. (1980), we applied a Bulked Segregant Analysis (BSA)-sequencing approach in an F2 population derived from a cross between Ontario 7818 (*Cf-6*) and MM-*Cf-0*. For this, 133 F2 seedlings were subjected to systemic PVX-based expression of *Avr6* and subsequently phenotyped based on presence or absence of a systemic HR (**Figure S6**). gDNA from 88 (out of 92) plants that underwent a systemic HR, indicating Avr6-responsiveness, was bulked into a responsive (R) pool, and from 40 (of 41) non-responders into a non-responsive (NR) pool. The expected 3R:1NR segregation pattern, under the hypothesis of *Cf-6* being a single dominant gene, was confirmed with a χ^2^ test (χ^2^=2.41, *p-*value=0.12) (**Table S6**). After short-read sequencing of the pools, we applied a Comparative Subsequence Sets analysis (CoSSa) bioinformatics pipeline (Prodhomme et al., 2019), to identify R-bulk-specific *k*-mers, which were subsequently mapped to the *S. lycopersicum* Heinz SL4.0 reference genome (Hosmani et al., 2019). We identified a small 2 Mb interval spanning from approximately 2.2 Mb to 4.2 Mb on the short arm of chromosome 12 that was significantly enriched for R-bulk-specific *k*-mers, indicating the presence of *Cf-6* (**Figure 8**). Potential resistance gene-encoding candidates for *Cf-6* within this region include eight RLP-encoding genes, divided over two distinct clusters, and two nucleotide-binding leucine-rich repeat (NB-LRR)-encoding genes. All these genes show significant allelic variation in Ontario 7818 compared to the Heinz 1706 SL4.0 reference and MM-*Cf-0*, indicating that they could be involved in recognition of Avr6 in Ontario 7818 (**Figure S7**). The region coincides with previous findings mapping *Cf-6* to the short arm of chromosome 12 (Kanwar et al., 1980), and the later finding for the two *Cf-Ecp5.12* alleles (Iakovidis et al., 2020).

**Figure 8:**
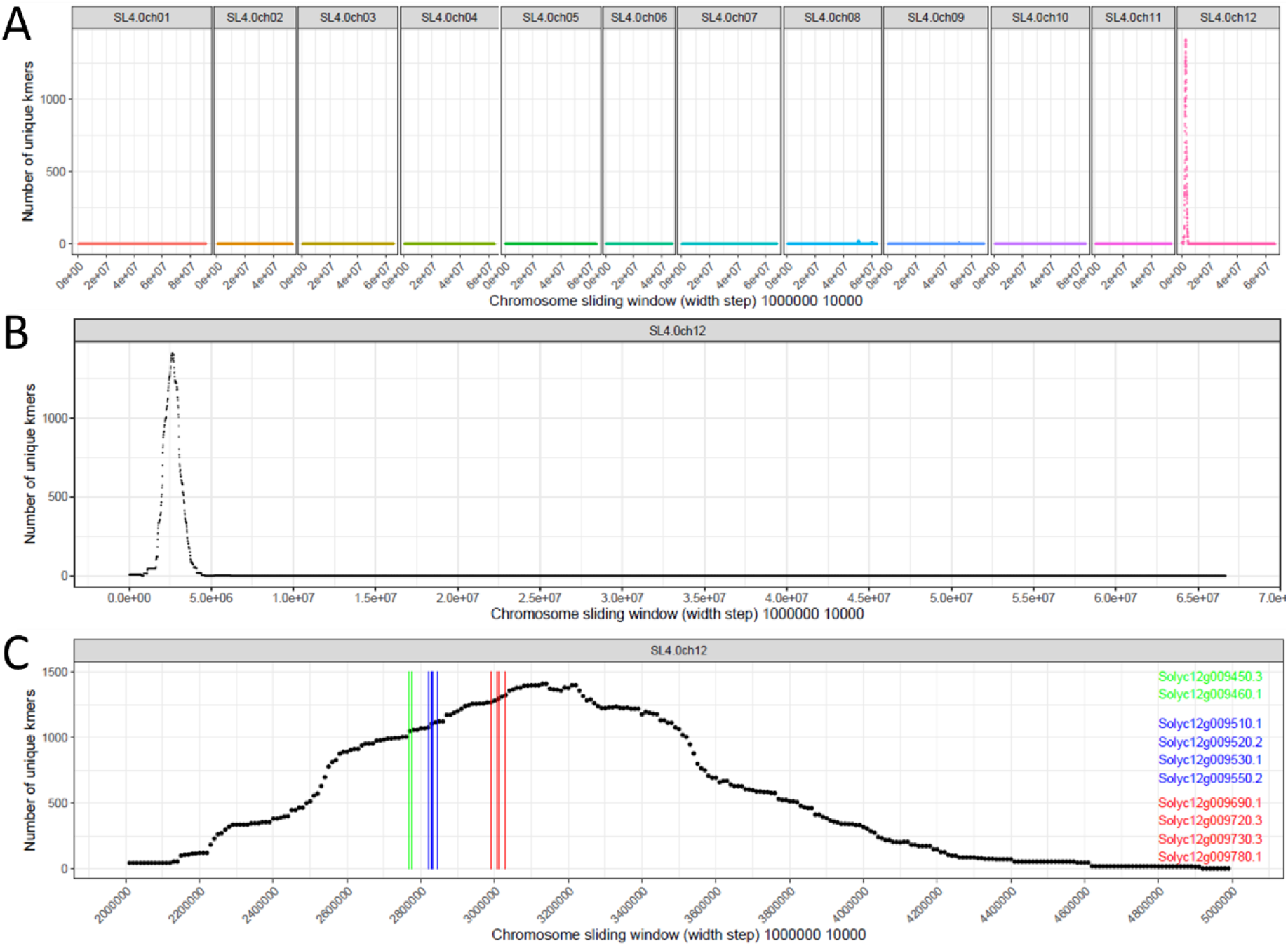
BSA-mapping of *Cf-6* (based on responsiveness to Avr6) in an Ontario 7818 x Moneymaker F2 population using Comparative Subsequence Sets analysis (CoSSa). Plots show R-pool-specific *k-*mers (resulting from intersection with Ontario 7818 specific *k-mers* and subtraction of NR-pool and MM-*Cf-0* specific *k-*mers), with respect to their position on the *S. lycopersicum* Heinz 1706 SL4.0 reference genome (Panel A). We found a strong correlation with a region spanning 2.2 to 4.2 Mb on the short arm of chromosome 12 (Panel B). One cluster containing two candidate NB-LRR-encoding genes (green lines) and two clusters both containing four RLP-encoding genes (blue and red lines, respectively) within this mapping interval are indicated (Panel C).

To determine which of these genes could be *Cf-6*, we predicted the domain structure of all ten encoded candidate Cf-6 proteins based on their predicted protein sequences from the Heinz 1706 ITAG4.0 reference annotation (**Figure S8, Tables S7 and S8**). Both NB-LRR proteins are predicted to encode for coiled-coil(CC)-NB-LRRs, whilst of the eight RLPs, only Solyc12g009510 and Solyc12g009520 are predicted to harbor an N-terminal SP for secretion, followed by an LRR domain and a transmembrane domain (TM), features that are deemed essential for RLP functionality. Interestingly, Solyc12g009510 is predicted to harbor a cytoplasmic LRR domain with a second C-terminal TM. The other RLPs do not show a complete predicted domain organization based on the reference, but because the structures of these proteins in Ontario 7818 remain unknown, they all remain candidates for Cf-6.

Finally, to verify the correlation between Avr6 recognition and resistance to *F. fulva* in the F2 population, we inoculated 52 new individuals from the same F2 population with PVX::*Avr6*, and inoculated cuttings of these individuals with the *F. fulva* race 0 strain 0WU (**Figure S9**). For every individual, we found a strict co-segregation of response to Avr6 with resistance to *F. fulva*, and of non-responsiveness with susceptibility. Again, we observed a 3R:1NR (40 R, 12 NR) segregation of the response to Avr6 and of disease resistance (χ^2^=0.103, *p-*value=0.75) (**Table S6**). This indicates that *Cf-6* is the only resistance gene providing resistance to *F. fulva* in the Ontario 7818 F2 population and further confirms that this gene confers resistance to *F. fulva* upon recognition of Avr6.

## Discussion

### Parallel loss and mutation of *Avr6* highlight repeatable paths to virulence

Avr6 encodes a 115-amino acid protein of unknown function that was originally cloned as Ecp5 (Laugé et al., 2000). Removal of its 17-amino acid N-terminal signal peptide for secretion yields a 98-amino acid mature apoplastic protein containing six cysteine residues. Avr6 expression is strongly induced *in planta* relative to expression in culture (Mesarich et al., 2014). Consistent with earlier reports (Iida et al., 2015; Stergiopoulos, de Kock, et al., 2007), we observed virtually no allelic variation in *Avr6* across a panel of recently collected Japanese strains and in all but four French race 2.6.9 strains and one additional European strain (P31). This pattern suggests that *Avr6* has largely evolved under purifying selection, with positive selection only recently imposed upon its deployment as an Avr determinant through *Cf-6*. As *Cf-6* becomes more widely deployed, it is reasonable to expect that additional virulence alleles of *Avr6* will emerge. The p.Cys30Tyr mutation in Avr6 of strain IPO 2679 is puzzling because this strain was collected in New Zealand in the 1980s (de la Rosa & Schol et al. 2024), when *Cf-6* is not known to have been commercially deployed. One possibility is that this mutation arose in a different genetic background or host context and only later encountered *Cf-6*. Interestingly, a previous study reported that an Avr6 p.Cys30Ala substitution (p.Cys13Ala in the mature protein) did not compromise the HR in the *Cf-Ecp5* donor line G1.1161 (Luderer et al., 2002). This difference suggests either that the Cf-Ecp5 receptor encoded by the G1.1161 chromosome 1 allele and Cf-6 encoded by the Ontario 7818 chromosome 12 allele differ in specificity, or that replacing Cys30 with a bulky, polar tyrosine has a more pronounced structural effect than substitution with a small, hydrophobic alanine. Distinguishing between these scenarios will require direct comparison of *Cf-Ecp5* alleles and their recognition specificities.

The diversity of polymorphisms in *Avr6* reported in Argentinian strains (Lucentini et al., 2021; Medina et al., 2015) raises questions about the origin of selection on this effector, because the sampled tomato cultivars are unlikely to harbor *Cf-6*/*Cf-Ecp5* alleles, given their susceptibility to strains expressing wild-type *Avr6*. Nevertheless, both *Ecp4* and *Avr6* were inferred to be under positive selection in the Argentinian *F. fulva* population, despite modest overall nucleotide diversity (Lucentini et al., 2021). One possible explanation is that selection on *Avr6* originated in wild tomato populations, where Cf-Ecp5-like receptors are common and widespread (Iakovidis et al., 2020), with subsequent spill-over of adapted *F. fulva* strains into cultivated tomato. A similar scenario could apply to IPO 2679, which may have acquired the p.Cys30Tyr mutation in a South American population before moving to New Zealand. However, given the current sampling bias towards cultivated tomato, we cannot rule out alternative explanations such as drift, founder effects, or selection by other, as yet unidentified host factors. A targeted sampling campaign of *F. fulva* and wild hosts in centres of tomato diversity, followed by genome- and population-level analyses (Bartoli & Roux, 2017; Lebeda & Burdon, 2023), will be required to clarify the evolutionary origin of *Avr6* variation.

### Stacked *Cf* genes can potentially be overcome by recombination between strains carrying different mutant *Avr* genes

Our data indicate that closely related *F. fulva* lineages can independently acquire virulence against different *Cf* genes via distinct mutations in multiple *Avr* genes. In particular, the clades comprising five race 2.6.9 and six race 2.9 strains appear to have overcome both *Cf-6* and *Cf-9B* through different mutations in *Avr6* and *Avr9B*: four race 2.6.9 strains carry a deletion of the *Avr6* open reading frame combined with Avr9B p.del129–133,p.L135V, the race 2.9 strains have a wild-type *Avr6* and Avr9B p.Cys107*, and one race 2.6.9 strain has Avr6 p.Gly53Val together with Avr9B p.Cys107* (**Figure 1**). These observations show that single *Cf* gene-mediated resistances can be eroded by multiple, independent effector mutations within relatively few fungal generations.

We also observe identical Avr mutations in strains with opposite mating types, including Avr2 p.Tyr41fs42, Avr2 p.Lys23fs34, Avr2 p.Asp54–Gly78del, Avr4 p.Cys109* and Avr4E p.Phe82Leu / p.Met93Thr. Similar patterns have been reported in a Turkish *F. fulva* population, where Avr alleles and both mating types occur at roughly equal frequencies (Gül et al., 2022; Gül et al., 2024). The most parsimonious explanation for these observations is that recombination occurs within *F. fulva* populations, consistent with earlier suggestions of cryptic sexual reproduction (Stergiopoulos, de Kock, et al., 2007; Stergiopoulos, Groenewald, et al., 2007; de la Rosa & Schol et al., 2024). Although we cannot yet formally distinguish sexual from parasexual recombination, either mechanism would facilitate the accumulation of multiple effector mutations in single genotypes and thereby increase the likelihood that *Cf* gene stacks are overcome over time. Nonetheless, the French strains we sequenced appear largely clonal: most cluster into a single, tight phylogenetic branch and show no evidence of recombination over the timeframe sampled. This suggests that, while recombination likely occurs in the broader species, it may be infrequent or episodic, and not necessarily detectable in all regional populations.

### Avr6 has sequence and structural similarity to putative effectors of other plant-pathogenic fungi

Sequence queries suggest that many plant-pathogenic Dothideomycete and Sordariomycete fungi produce one or more Avr6-like proteins, of which one, *Pf*Avr6, was identified in *P. fuligena*, a pathogen closely related to *F. fulva* that causes black leaf mold of tomato (Phengsintham et al., 1938). Given that the mature version of *Pf*Avr6 has a high level of sequence identity to the mature version of Avr6 (∼71.4%), including Gly53 and Tyr88 (**Figure S5**), and that there is evidence of resistance to *P. fuligena* in cultivated and wild tomato (Hartman & Wang, 1993; Wang et al., 1995), it would be interesting to determine if it is recognized by *Cf-6* tomato plants. If so, *Cf-6* could also provide protection against *P. fuligena*. However, the emergence of *Cf-6*-breaking *F. fulva* strains suggests that any such protection will likely be short-lived as *P. fuligena* may adapt in a similar way. Searches based on the predicted tertiary structure of Avr6 revealed similarity to LARS effectors, which have so far been identified in at least 13 plant-pathogenic fungi (Lazar et al., 2022). Examples include four Avr effectors from *L. maculans*, as well as *F. fulva* Ecp11-1 (Lazar et al., 2022; Mesarich et al., 2018). In the *L. maculans*–*Brassica* pathosystem, recognition of the LARS effectors AvrLm3 and AvrLm5-9 by their corresponding R proteins can be masked by another LARS effector, AvrLm4-7 (Ghanbarnia et al., 2018; Plissonneau et al., 2016). However, it is not yet known whether Avr6 (or Ecp11-1) has the capacity to mask the recognition of other *F. fulva* effectors in susceptible tomato plants. The predicted tertiary structure of Avr6 displays several notable differences to the LARS effectors described above, reflected in a lack of sequence similarity. LARS-like effectors lacking such similarity or possessing subtle differences in their predicted tertiary folds have been identified in other plant-pathogenic fungi including in *Venturia inaequalis*, which causes apple scab disease (Rocafort et al., 2022).

### *Cf-6* maps to the chromosome 12 allele of *Cf-Ecp5* at the *ANDROMEDA* locus

Initially, an HR was observed upon elicitation with the then*-*called Ecp5 protein (Laugé et al., 2000) in the introgression line G1.1161, an F_3_BC_1_ resulting from a cross between *S. pimpinellifolium* CGN15529 and MM-*Cf-0* (Haanstra et al., 2000). The *Cf-Ecp5* gene in this accession was mapped to the *AURORA* (*AU*) locus on the short arm of chromosome 1 (Haanstra et al., 2000), but a later study found that rather, it was tightly linked with *Cf-9* at the *MW* locus (Iakovidis et al., 2020). This study also reported two other independent *Cf-Ecp5* loci, i.e. *Cf-Ecp5.7* at the *CENTAURI* (*CE*) locus on the long arm of chromosome 7 in *S. pimpinellifolium* LA0722, and *Cf-Ecp5.12* at the *ANDROMEDA* (*AN*) locus located within a region on the short arm of chromosome 12 spanning from approximately 2.1 to 3.0 Mb in *S. lycopersicum* Ontario 7522 and *S. pimpinellifolium* LA2852 (Iakovidis et al., 2020). Our BSA–CoSSa mapping of *Cf-6* in Ontario 7818 to a ∼2 Mb interval on the short arm of chromosome 12 that overlaps the *Cf-Ecp5.12* mapping interval strongly suggests that *Cf-6* and *Cf-Ecp5.12* represent alleles of the same underlying gene, although allelic diversification between Ontario 7818 and the other *Cf-Ecp5.12* donors cannot yet be excluded.

The study of Iakovidis *et al*. (2020) employed a marker at 3.014 Mb, which showed 4.5% recombination with the *Cf-Ecp5.12* locus. Assuming that the *Cf-6* allele from Ontario 7818 and the *Cf-Ecp5* allele from Ontario 7522 are the same and encode an RLP, it is more likely to reside within the “blue” RLP-encoding gene cluster at 2.85 Mb, rather than at the “red” cluster at around 3.0 Mb (**Figure 8C**), which is roughly at the same location as the mentioned marker of Iakovidis et al. (2020). Although originating from the same breeding program, Ontario 7818 and 7522 do appear to have different donors, namely *S. pimpinellifolium* PI 211839 and PI 124161, respectively. PI 124161 appears to originate from Guatemala (**Table S2**), whilst the origin of PI 211839 is unknown. Since all three mapped *Cf-6/Cf-Ecp5* loci identified to date originate from *S. pimpinellifolium*, it would be valuable to determine the genomic locations of corresponding alleles in other tomato species such as *S. peruvianum* or *S. chilense*, as several accessions from these species also respond to Avr6 (Iakovidis et al., 2020). Mapping these alleles may reveal additional, distinct *Cf-6*/Cf-Ecp5 loci and thereby provide further insight into the evolution, diversification, and geographic distribution of *Cf* genes across wild tomato species.

Finally, our results help resolve earlier conflicting reports on the chromosomal position of *Cf-6*: neither chromosome 6 (Grushetskaya et al., 2007) nor chromosome 11 (Wang et al., 2007) appear to contribute to Avr6 recognition. Instead, our data align with the more recent findings of Iakovidis et al. (2020), which map *Cf-Ecp5.12*—now strongly supported to be allelic with *Cf-6*—to the short arm of chromosome 12.

The identified candidate RLP-encoding genes within our mapping region in Ontario7818 are prime candidates for *Cf-6*, particularly *Solyc12g009510* and *Solyc12g009520*, given their predicted complete protein structures in Heinz 1706, although it is also possible that *Cf-6* of Ontario 7818 is not annotated in the Heinz 1706 genome. This could be resolved by generating a *de novo* genome assembly of Ontario 7818, or by resistance gene enrichment sequencing (RENseq) (Lin et al., 2020; Witek et al., 2016), revealing any structural differences that are often found across different accessions/species in RLP-encoding genes (de Kock et al., 2005; Dixon et al., 1998; Dixon et al., 1996; Man et al., 2020). For example, the recognition of Avr9 in wild *S. chilense* accessions is highly region-dependent and the *Cf-9* locus is highly polymorphic amongst these accessions (Kahlon et al., 2020).

### Future strategies for developing novel, more durable resistance to tomato leaf mold

Future breeding efforts should focus on achieving more durable resistance against *F. fulva*, as breakdown of the recently deployed *Cf-6* gene appears to have occurred more rapidly than the breakdown of the multi-gene *Cf-9* locus (de la Rosa & Schol et al., 2024), underscoring that single *Cf* gene strategies are unlikely to be durable. Wild tomato germplasm that triggers specific HR responses to one or more of the nine newly identified Ecp effectors (ten including Ecp17-1) (Mesarich et al., 2018) likely contains an equivalent number of untapped *Cf-Ecp* genes that could be incorporated into breeding programs. However, before deployment, the durability of such genes should be carefully assessed. One promising approach is to generate *Ecp* knockout strains to determine their contribution to *F. fulva* virulence, as recently demonstrated for *Ecp20-2* (Karimi-Jashni et al., 2022). The long-standing assumption that Ecp effectors are more conserved—and therefore more essential for virulence—than Avr effectors (Ökmen & de Wit, 2012; Stergiopoulos, de Kock, et al., 2007) is challenged by our findings, which show that *Avr6*/*Ecp5* can be readily lost or mutated without substantial fitness cost.

At present, pyramiding multiple *Cf* genes appears to be the most durable strategy, as sequential loss of several effector genes is expected to occur far more slowly than loss of individual effectors, even when sporadic recombination is considered. Nevertheless, truly durable resistance will require an innovative approach, such as *Cf* genes matching effectors that are essential for pathogenicity, or the discovery of susceptibility factors that can be targeted to remove essential host functions exploited by the pathogen (Koseoglou et al., 2022; Sun KaiLe et al., 2014).

Progress toward durable resistance can be accelerated by comprehensive population-level studies of *F. fulva* across its full geographic range, including wild host populations, to identify which effectors and *Cf* genes are most constrained, which are most variable, and how rapidly new alleles arise. Such ecological and evolutionary data should be integrated with pathogen–host interaction modelling (Susi et al., 2020; Märkle et al., 2024; McMullan et al., 2025), enabling predictions of which *Cf* genes are under the strongest purifying selection and, consequently offer the greatest long-term durability.

## Acknowledgements

We would like to thank Bert Essenstam and colleagues of the Unifarm greenhouse facilities at Wageningen University for excellent plant care, Dr Pedro Ballati of the Universidad Nacional de La Plata, Buenos Aires, Argentina for kindly providing the Argentinian *F. fulva* strains, Dr Phillipe Nicot of INRAE Centre de Recherche PACA, Montfavet, France for providing a selection of *F. fulva* strains from the Southern region of France and the growers of the Cerafel association, St. Martin-des-Champs, France, for providing *F. fulva* strains from the Brittany region. This research was largely funded by the EU Horizon 2020 HARNESSTOM project (Grant agreement no.: 101000716).

## Author contributions

CRS, LF, YB, MHAJ, IS, pJGMdW and CHM designed the research; CRS, LF, AH, RJ, SdlR, BÖ, AMH, SLvZ, LDHV, CM, KM, AZZ, DE and CHM performed the research; CRS, LF, AH, AMH, SLvZ, MT, YI, AZZ, DE, A-MAW and CHM carried out the data analysis, collection or interpretation; CRS, LF and CHM wrote the manuscript. All authors reviewed the manuscript and approved it for publication.

## Data availability

All *F. fulva* sequencing data generated in this study is deposited in GenBank (PRJNA1271984 and PRJNA961892), with specific BioSample and Sequence Read Archive (SRA) accession numbers listed in **Table S3**.

## Supporting information (brief legends)

**Figure S1:** Phylogeny and sequence comparison of effector genes across a panel of 33 *F. fulva* genomes.

**Figure S2:** Novel strains of *F. fulva* can infect Ontario 7818 tomato plants harboring the *Cf-6* gene.

**Figure S3:** *Ecp5* is part of an 18 kb deletion in four Race 6 strains of *F. fulva*.

**Figure S4:** An intro-exon border mutation in *Avr6* allows *F. fulva* to partially overcome *Cf-6*-mediated resistance.

**Figure S5:** Avr6-like proteins are predicted to be produced by many plant-pathogenic Dothideomycete and Sordariomycete fungi.

**Figure S6:** The Ontario 7818 x MM-*Cf-0* F2 BSA mapping population segregates for an HR to Avr6.

**Figure S7:** Receptor-encoding candidates within the mapping interval of *Cf-6* exhibit significant numbers of R-pool-specific SNPs compared to the Heinz 1706 SL4.0 reference genome and the NR-pool.

**Figure S8:** Domain predictions of receptor proteins within the mapping interval of *Cf-6*, based on the Heinz 1706 reference annotation (ITAG 4.0).

**Figure S9:** Recognition of Avr6 correlates with tomato leaf mold disease resistance in Ontario 7818 x MM-*Cf-0* F2 individuals.

**Notes S1:** Predicted Avr6-like sequences from fungi.

**Table S1:** *F. fulva* strains used in this study for whole genome sequencing (strains with an alternate ID; as used for clarity in the manuscript) and for *Avr6* allelic variation analysis (strains without alternate ID).

**Table S2:** Detailed information regarding the plant accessions used and discussed in this study as provided by the Centrum voor Genetische Bronnen Nederland (CGN).

**Table S3:** Quality statistics for *F. fulva* genome assemblies generated in this study.

**Table S4:** Quality statistics for publicly available *F. fulva* genome assemblies.

**Table S5:** Fastp statistics for the R- and NR-pools used for BSA mapping of the *Cf-6* locus in the Ontario 7818 x MM-*Cf-0* F2 tomato mapping population.

**Table S6:** Statistical analysis of the observed segregation ratios based on PVX::*Avr6* response in the Ontario 7818 x MM-*Cf-0* F2 tomato populations by means of χ2 tests.

**Table S7:** Interpro domain prediction results for Cf-6 candidate receptor proteins, based on their Heinz 1706 ITAG4.0 annotation.

**Table S8:** Summary of Interpro domain prediction results for Cf-6 candidate receptor proteins, based on their Heinz 1706 ITAG4.0 annotation.

## References

Andrews, S. (2010) FastQC: a quality control tool for high throughput sequence data. *Available online at:* http://www.bioinformatics.babraham.ac.uk/projects/fastqc.

Andrews, S. F. (2019). V0. 11.9: A quality control tool for high throughput sequence data. Babraham Bioinformatics: Cambridge, UK. Available online: http://www.bioinformatics.babraham.ac.uk/projects/fastqc.

Bankevich, A., Nurk, S., Antipov, D., Gurevich, A. A., Dvorkin, M., Kulikov, A. S., Lesin, V. M., Nikolenko, S. I., Pham, S., & Prjibelski, A. D. (2012). SPAdes: a new genome assembly algorithm and its applications to single-cell sequencing. Journal of Computational Biology, 19(5), 455–477.

Bartoli, C., & Roux, F. (2017). Genome-wide association studies in plant pathosystems: toward an ecological genomics approach. Frontiers in Plant Science, 8, 763.

Blondeau, K., Blaise, F., Graille, M., Kale, S. D., Linglin, J., Ollivier, B., Labarde, A., Lazar, N., Daverdin, G., & Balesdent, M. H. (2015). Crystal structure of the effector AvrLm4–7 of *Leptosphaeria maculans* reveals insights into its translocation into plant cells and recognition by resistance proteins. The Plant Journal, 83(4), 610–624.

Bolger, A. M., Lohse, M., & Usadel, B. (2014). Trimmomatic: a flexible trimmer for Illumina sequence data. Bioinformatics, 30(15), 2114–2120.

Capella-Gutiérrez, S., Silla-Martínez, J. M., & Gabaldón, T. (2009). trimAl: a tool for automated alignment trimming in large-scale phylogenetic analyses. Bioinformatics, 25(15), 1972–1973.

Chen, S. (2023). Ultrafast one-pass FASTQ data preprocessing, quality control, and deduplication using fastp. Imeta, 2(2), e107.

Chen, S., Zhou, Y., Chen, Y., & Gu, J. (2018). fastp: an ultra-fast all-in-one FASTQ preprocessor. Bioinformatics, 34(17), i884–i890.

Cooke, M. (1883). New american fungi. Grevillea, 12, 22–33.

de Kock, M. J., Brandwagt, B. F., Bonnema, G., de Wit, P. J., & Lindhout, P. (2005). The tomato Orion locus comprises a unique class of *Hcr9* genes. Molecular Breeding, 15(4), 409–422.

de la Rosa, S., Schol, C. R., Ramos Peregrina, Á., Winter, D. J., Hilgers, A. M., Maeda, K., Iida, Y., Tarallo, M., Jia, R., Beenen, H. G., Rocafort, M., de Wit, P. J. G. M., Bowen, J. K., Bradshaw, R. E., Joosten, M. H. A. J., Bai, Y., & Mesarich, C. H. (2024). Sequential breakdown of the *Cf-9* leaf mould resistance locus in tomato by *Fulvia fulva*. New Phytologist, 243(4), 1522–1538.

de Wit, P., & Flach, W. (1979). Differential accumulation of phytoalexins in tomato leaves but not in fruits after inoculation with virulent and avirulent races of *Cladosporium fulvum*. Physiological Plant Pathology, 15(3), 257–267.

de Wit, P. J. (2016). *Cladosporium fulvum* effectors: weapons in the arms race with tomato. Annual Review of Phytopathology, 54, 1–23.

de Wit, P. J., Van Der Burgt, A., Ökmen, B., Stergiopoulos, I., Abd-Elsalam, K. A., Aerts, A. L., Bahkali, A. H., Beenen, H. G., Chettri, P., & Cox, M. P. et al. (2012). The genomes of the fungal plant pathogens *Cladosporium fulvum* and *Dothistroma septosporum* reveal adaptation to different hosts and lifestyles but also signatures of common ancestry. PLoS Genetics, 8(11), e1003088.

DeLano, W. L. (2002). Pymol: An open-source molecular graphics tool. CCP4 Newsletter on Protein Crystallography, 40(1), 82–92.

Dixon, M. S., Hatzixanthis, K., Jones, D. A., Harrison, K., & Jones, J. D. (1998). The tomato *Cf-5* disease resistance gene and six homologs show pronounced allelic variation in leucine-rich repeat copy number. The Plant Cell, 10(11), 1915–1925.

Dixon, M. S., Jones, D. A., Keddie, J. S., Thomas, C. M., Harrison, K., & Jones, J. D. (1996). The tomato *Cf-2* disease resistance locus comprises two functional genes encoding leucine-rich repeat proteins. Cell, 84(3), 451–459.

Edgar, R. C. (2022). Muscle5: High-accuracy alignment ensembles enable unbiased assessments of sequence homology and phylogeny. Nature Communications, 13(1), 6968.

Ghanbarnia, K., Ma, L., Larkan, N. J., Haddadi, P., Fernando, W. G. D., & Borhan, M. H. (2018). *Leptosphaeria maculans* AvrLm9: a new player in the game of hide and seek with AvrLm4-7. Molecular Plant Pathology, 19(7), 1754–1764.

Grushetskaya, Z., Lemesh, V., Poliksenova, V., & Khotyleva, L. (2007). Mapping of the *Cf-6* tomato leaf mould resistance locus using SSR markers. Russian Journal of Genetics, 43(11), 1266–1270.

Gül, E., Karakaya, A., & Ergül, A. (2022). Determination of mating types of *Fulvia fulva* isolates from Turkey. Tropical Plant Pathology, 47(3), 421–429.

Gül, E., Karakaya, A., & Ergül, A. (2024). The race distribution of the *Fulvia fulva* populations and mutations in the Avr genes in Turkey. Journal of Phytopathology, 172(1), e13273.

Gurevich, A., Saveliev, V., Vyahhi, N., & Tesler, G. (2013). QUAST: quality assessment tool for genome assemblies. Bioinformatics, 29(8), 1072–1075.

Gust, A. A., & Felix, G. (2014). Receptor like proteins associate with SOBIR1-type of adaptors to form bimolecular receptor kinases. Current Opinion in Plant Biology, 21, 104–111.

Haanstra, J., Meijer-Dekens, F., Lauge, R., Seetanah, D., Joosten, M. H., de Wit, P. J., & Lindhout, P. (2000). Mapping strategy for resistance genes against *Cladosporium fulvum* on the short arm of chromosome 1 of tomato*: Cf-ECP5* near the *Hcr9* Milky Way cluster. Theoretical and Applied Genetics, 101(4), 661–668.

Hammond-Kosack, K., Staskawicz, B., Jones, J., & Baulcombe, D. (1995). Functional expression of a fungal avirulence gene from a modified potato virus X genome. Molecular Plant-Microbe interactions, 8(1), 181–185.

Hartman, G., & Wang, T. (1993). Resistance in Lycopersicon species to black leaf mold caused by *Pseudocercospora fuligena*. Euphytica, 71(1), 125–130.

Hoang, D. T., Chernomor, O., Von Haeseler, A., Minh, B. Q., & Vinh, L. S. (2018). UFBoot2: improving the ultrafast bootstrap approximation. Molecular Biology and Evolution, 35(2), 518–522.

Hosmani, P. S., Flores-Gonzalez, M., van de Geest, H., Maumus, F., Bakker, L. V., Schijlen, E., van Haarst, J., Cordewener, J., Sanchez-Perez, G., & Peters, S. (2019). An improved de novo assembly and annotation of the tomato reference genome using single-molecule sequencing, Hi-C proximity ligation and optical maps. bioRxiv, 767764.

Huang, W. R., & Joosten, M. H. (2024). Immune signaling: receptor-like proteins make the difference. Trends in Plant Science, 30(1), 54–68.

Iakovidis, M., Soumpourou, E., Anderson, E., Etherington, G., Yourstone, S., & Thomas, C. (2020). Genes encoding recognition of the *Cladosporium fulvum* effector protein Ecp5 are encoded at several loci in the tomato genome. G3 Genes|Genomes|Genetics, 10(5), 1753–1763.

Iida, Y., van ‘t Hof, P., Beenen, H., Mesarich, C., Kubota, M., Stergiopoulos, I., Mehrabi, R., Notsu, A., Fujiwara, K., & Bahkali, A. (2015). Novel mutations detected in avirulence genes overcoming tomato *Cf* resistance genes in isolates of a Japanese population of *Cladosporium fulvum*. PLoS ONE, 10(4), e0123271.

Jones, D. A., Thomas, C. M., Hammond-Kosack, K. E., Balint-Kurti, P. J., & Jones, J. (1994). Isolation of the tomato *Cf-9* gene for resistance to *Cladosporium fulvum* by transposon tagging. Science, 266(5186), 789–793.

Joosten, M., Vogelsang, R., Cozijnsen, T. J., Verberne, M. C., & de Wit, P. (1997). The biotrophic fungus *Cladosporium fulvum* circumvents Cf-4-mediated resistance by producing unstable AVR4 elicitors. The Plant Cell, 9(3), 367–379.

Joosten, M. H., Cozijnsen, T. J., & de Wit, P. J. (1994). Host resistance to a fungal tomato pathogen lost by a single base-pair change in an avirulence gene. Nature, 367(6461), 384–386.

Joosten, M. H., & de Wit, P. J. (1989). Identification of several pathogenesis-related proteins in tomato leaves inoculated with *Cladosporium fulvum* (syn. *Fulvia fulva*) as 1, 3-β-glucanases and chitinases. Plant Physiology, 89(3), 945–951.

Jumper, J., Evans, R., Pritzel, A., Green, T., Figurnov, M., Ronneberger, O., Tunyasuvunakool, K., Bates, R., Žídek, A., & Potapenko, A. (2021). Highly accurate protein structure prediction with AlphaFold. Nature, 596(7873), 583–589.

Kahlon, P. S., Seta, S. M., Zander, G., Scheikl, D., Hückelhoven, R., Joosten, M. H., & Stam, R. (2020). Population studies of the wild tomato species *Solanum chilense* reveal geographically structured major gene-mediated pathogen resistance. Proceedings of the Royal Society B, 287(1941), 20202723.

Kalyaanamoorthy, S., Minh, B. Q., Wong, T. K., Von Haeseler, A., & Jermiin, L. S. (2017). ModelFinder: fast model selection for accurate phylogenetic estimates. Nature Methods, 14(6), 587–589.

Kang, W.-H., & Yeom, S.-I. (2018). Genome-wide identification, classification, and expression analysis of the receptor-like protein family in tomato. The Plant Pathology Journal, 34(5), 435.

Kanwar, J., Kerr, E., & Harney, P. (1980). Linkage of the *Cf-12* to *Cf-24* genes for resistance to tomato leaf mold *Cladosporium fulvum* Cke. Report of the Tomato Genetics Cooperative, 30, 22–23.

Karimi-Jashni, M., Maeda, K., Yazdanpanah, F., de Wit, P. J., & Iida, Y. (2022). An integrated omics approach uncovers the novel effector Ecp20-2 required for full virulence of *Cladosporium fulvum* on tomato. Frontiers in Microbiology, 13, 919809.

Kearse, M., Moir, R., Wilson, A., Stones-Havas, S., Cheung, M., Sturrock, S., Buxton, S., Cooper, A., Markowitz, S., & Duran, C. (2012). Geneious Basic: an integrated and extendable desktop software platform for the organization and analysis of sequence data. Bioinformatics, 28(12), 1647–1649.

Kokot, M., Długosz, M., & Deorowicz, S. (2017). KMC 3: counting and manipulating *k*-mer statistics. Bioinformatics, 33(17), 2759–2761.

Koseoglou, E., van der Wolf, J. M., Visser, R. G., & Bai, Y. (2022). Susceptibility reversed: modified plant susceptibility genes for resistance to bacteria. Trends in Plant Science, 27(1), 69–79.

Langmead, B., & Salzberg, S. L. (2012). Fast gapped-read alignment with Bowtie 2. Nature Methods, 9(4), 357–359.

Laugé, R., Goodwin, P. H., de Wit, P. J., & Joosten, M. H. (2000). Specific HR-associated recognition of secreted proteins from *Cladosporium fulvum* occurs in both host and non-host plants. The Plant Journal, 23(6), 735–745.

Lazar, N., Mesarich, C. H., Petit-Houdenot, Y., Talbi, N., Li de la Sierra-Gallay, I., Zélie, E., Blondeau, K., Gracy, J., Ollivier, B., & Blaise, F. (2022). A new family of structurally conserved fungal effectors displays epistatic interactions with plant resistance proteins. PLoS Pathogens, 18(7), e1010664.

Lebeda, A., & Burdon, J. J. (2023). Studying wild plant pathosystems to understand crop plant pathosystems: status, gaps, challenges, and perspectives. Phytopathology, 113(3), 365–380.

Li, H., & Durbin, R. (2009). Fast and accurate short read alignment with Burrows–Wheeler transform. Bioinformatics, 25(14), 1754–1760.

Lin, X., Armstrong, M., Baker, K., Wouters, D., Visser, R. G., Wolters, P. J., Hein, I., & Vleeshouwers, V. G. (2020). *RLP/K* enrichment sequencing; a novel method to identify *receptor-like protein* (*RLP*) and *receptor-like kinase* (*RLK*) genes. New Phytologist, 227(4), 1264–1276.

Lucentini, C. G., Medina, R., Franco, M. E., Saparrat, M. C., & Balatti, P. A. (2021). *Fulvia fulva* [syn. *Cladosporium fulvum, Passalora fulva*] races in Argentina are evolving through genetic changes and carry polymorphic *avr* and *ecp* gene sequences. European Journal of Plant Pathology, 159(3), 525–542.

Luderer, R., de Kock, M. J., Dees, R. H., de Wit, P. J., & Joosten, M. H. (2002). Functional analysis of cysteine residues of ECP elicitor proteins of the fungal tomato pathogen *Cladosporium fulvum*. Molecular Plant Pathology, 3(2), 91–95.

Magoč, T., & Salzberg, S. L. (2011). FLASH: fast length adjustment of short reads to improve genome assemblies. Bioinformatics, 27(21), 2957–2963.

Man, J., Gallagher, J. P., & Bartlett, M. (2020). Structural evolution drives diversification of the large LRR-RLK gene family. New Phytologist, 226(5), 1492–1505.

Manni, M., Berkeley, M. R., Seppey, M., Simão, F. A., & Zdobnov, E. M. (2021). BUSCO update: novel and streamlined workflows along with broader and deeper phylogenetic coverage for scoring of eukaryotic, prokaryotic, and viral genomes. Molecular Biology and Evolution, 38(10), 4647–4654.

Märkle, H., John, S., Metzger, L., Ansari, M. A., Pedergnana, V., & Tellier, A. (2024). Inference of host–pathogen interaction matrices from genome-wide polymorphism data. Molecular Biology and Evolution, 41(9), msae176.

McMullan, M., Percival-Alwyn, L., Kaithakottil, G. G., Gardiner, L. J., Hill, R., Yvanne, H., … & Hall, N. (2025). Developing a crop-wild-reservoir pathogen system to understand pathogen evolution and emergence. eLife, 14, e91245.

Medina, R., López, S. M., Franco, M. E., Rollan, C., Ronco, B. L., Saparrat, M. C., de Wit, P. J., & Balatti, P. A. (2015). A survey on occurrence of *Cladosporium fulvum* identifies race 0 and race 2 in tomato-growing areas of Argentina. Plant Disease, 99(12), 1732–1737.

Mesarich, C. H., Barnes, I., Bradley, E. L., de la Rosa, S., de Wit, P. J., Guo, Y., Griffiths, S. A., Hamelin, R. C., Joosten, M. H., Lu, M., McCarthy, H. M., Schol, C. R., Stergiopoulos, I., Tarallo, M., Zaccaron, A. Z., & Bradshaw, R. E. (2023). Beyond the genomes of *Fulvia fulva* (syn. *Cladosporium fulvum*) and *Dothistroma septosporum*: New insights into how these fungal pathogens interact with their host plants. Molecular Plant Pathology, 24(5), 474–494.

Mesarich, C. H., Griffiths, S. A., van der Burgt, A., Ökmen, B., Beenen, H. G., Etalo, D. W., Joosten, M. H. A. J., & de Wit, P. J. G. M. (2014). Transcriptome sequencing uncovers the *Avr5* avirulence gene of the tomato leaf mold pathogen *Cladosporium fulvum*. Molecular Plant-Microbe Interactions, 27(8), 846–857.

Mesarich, C. H., Ӧkmen, B., Rovenich, H., Griffiths, S. A., Wang, C., Karimi Jashni, M., Mihajlovski, A., Collemare, J., Hunziker, L., Deng, C. H., ven der Burgt, A., Beened, H. G., Templeton, M. D., Bradshaw, R.E., & det Wit, P. J. G. M. (2018). Specific hypersensitive response–associated recognition of new apoplastic effectors from *Cladosporium fulvum* in wild tomato. Molecular Plant-Microbe Interactions, 31(1), 145–162.

Meyer, U., & Gärber, U. (2021). Bestimmung der in Deutschland vorkommenden Pathotypen des Erregers der Samtfleckenkrankheit an Tomate, *Fulvia fulva*. Journal of Cultivated Plants/Journal für Kulturpflanzen, 73.

Minh, B. Q., Schmidt, H. A., Chernomor, O., Schrempf, D., Woodhams, M. D., Von Haeseler, A., & Lanfear, R. (2020). IQ-TREE 2: new models and efficient methods for phylogenetic inference in the genomic era. Molecular Biology and Evolution, 37(5), 1530–1534.

Mirdita, M., Schütze, K., Moriwaki, Y., Heo, L., Ovchinnikov, S., & Steinegger, M. (2022). ColabFold: making protein folding accessible to all. Nature Methods, 19(6), 679–682.

Morgulis, A., Coulouris, G., Raytselis, Y., Madden, T. L., Agarwala, R., & Schäffer, A. A. (2008). Database indexing for production MegaBLAST searches. Bioinformatics, 24(16), 1757–1764.

Murray, M., & Thompson, W. (1980). Rapid isolation of high molecular weight plant DNA. Nucleic Acids Research, 8(19), 4321–4326.

Nielsen, H. (2017). Predicting secretory proteins with SignalP. Protein Function Prediction: Methods and Protocols, 59–73.

Ökmen, B. (2013). Identification and characterization of novel effectors of Cladosporium fulvum. PhD Thesis, Wageningen University and Research.

Ökmen, B., & de Wit, P. J. (2012). *Cladosporium fulvum*–tomato pathosystem: Fungal infection strategy and plant responses. Molecular Plant Immunity, 211–224.

Parniske, M., Hammond-Kosack, K. E., Golstein, C., Thomas, C. M., Jones, D. A., Harrison, K., Wulff, B. B., & Jones, J. D. (1997). Novel disease resistance specificities result from sequence exchange between tandemly repeated genes at the *Cf-4/9* locus of tomato. Cell, 91(6), 821–832.

Paysan-Lafosse, T., Blum, M., Chuguransky, S., Grego, T., Pinto, B. L., Salazar, G. A., Bileschi, M. L., Bork, P., Bridge, A., & Colwell, L. (2023). InterPro in 2022. Nucleic Acids Research, 51(D1), D418–D427.

Penouilh-Suzette, C., Fourré, S., Besnard, G., Godiard, L., & Pecrix, Y. (2020). A simple method for high molecular-weight genomic DNA extraction suitable for long-read sequencing from spores of an obligate biotroph oomycete. Journal of Microbiological Methods, 178, 106054.

Phengsintham, P., Chukeatirote, E., McKenzie, E., Hyde, K., & Braun, U. (1938). Tropical phytopathogens 2: *Pseudocercospora fuligena*. J. Sci, 66(8).

Plissonneau, C., Daverdin, G., Ollivier, B., Blaise, F., Degrave, A., Fudal, I., Rouxel, T., & Balesdent, M. H. (2016). A game of hide and seek between avirulence genes *AvrLm4-7* and *AvrLm3* in *Leptosphaeria maculans*. New Phytologist, 209(4), 1613–1624.

Prodhomme, C., Esselink, D., Borm, T., Visser, R. G., Van Eck, H. J., & Vossen, J. H. (2019). Comparative Subsequence Sets Analysis (CoSSA) is a robust approach to identify haplotype specific SNPs; mapping and pedigree analysis of a potato wart disease resistance gene *Sen3*. Plant Methods, 15, 1–20.

Quinlan, A. R., & Hall, I. M. (2010). BEDTools: a flexible suite of utilities for comparing genomic features. Bioinformatics, 26(6), 841–842.

Ranallo-Benavidez, T. R., Jaron, K. S., & Schatz, M. C. (2020). GenomeScope 2.0 and Smudgeplot for reference-free profiling of polyploid genomes. Nature Communications, 11(1), 1432.

Robinson, J. T., Thorvaldsdóttir, H., Winckler, W., Guttman, M., Lander, E. S., Getz, G., & Mesirov, J. P. (2011). Integrative genomics viewer. Nature Biotechnology, 29(1), 24–26.

Rocafort, M., Bowen, J. K., Hassing, B., Cox, M. P., McGreal, B., de la Rosa, S., Plummer, K. M., Bradshaw, R. E., & Mesarich, C. H. (2022). The *Venturia inaequalis* effector repertoire is dominated by expanded families with predicted structural similarity, but unrelated sequence, to avirulence proteins from other plant-pathogenic fungi. BMC biology, 20(1), 246.

Rooney, H. C., Van’t Klooster, J. W., van der Hoorn, R. A., Joosten, M. H., Jones, J. D., & de Wit, P. J. (2005). *Cladosporium* Avr2 inhibits tomato Rcr3 protease required for Cf-2-dependent disease resistance. Science, 308(5729), 1783–1786.

Sangster, W. (2022). ISF Passalora fulva project tomato. https://worldseed.org/document/isf-passalora-fulva-project-tomato-final-report-2019-2022/

Stergiopoulos, I., de Kock, M. J., Lindhout, P., & de Wit, P. J. (2007). Allelic variation in the effector genes of the tomato pathogen *Cladosporium fulvum* reveals different modes of adaptive evolution. Molecular Plant-Microbe Interactions, 20(10), 1271–1283.

Stergiopoulos, I., Groenewald, M., Staats, M., Lindhout, P., Crous, P. W., & de Wit, P. J. (2007). Mating-type genes and the genetic structure of a world-wide collection of the tomato pathogen *Cladosporium fulvum*. Fungal Genetics and Biology, 44(5), 415–429.

Sun KaiLe, S. K., Zheng Zheng, Z. Z., Jacobsen, E., Visser, R. G., & Bai YuLing, B. Y. (2014). Breeding for disease resistance by editing plant susceptibility genes. CABI Reviews, (2014), 1–5.

Susi, H., Burdon, J. J., Thrall, P. H., Nemri, A., & Barrett, L. G. (2020). Genetic analysis reveals long-standing population differentiation and high diversity in the rust pathogen *Melampsora lini*. PLoS pathogens, 16(8), e1008731.

Takken, F. L., Thomas, C. M., Joosten, M. H., Golstein, C., Westerink, N., Hille, J., Nijkamp, H. J. J., de Wit, P. J., & Jones, J. D. (1999). A second gene at the tomato *Cf-4* locus confers resistance to *Cladosporium fulvum* through recognition of a novel avirulence determinant. The Plant Journal, 20(3), 279–288.

Thomas, C. M., Jones, D. A., Parniske, M., Harrison, K., Balint-Kurti, P. J., Hatzixanthis, K., & Jones, J. (1997). Characterization of the tomato *Cf-4* gene for resistance to *Cladosporium fulvum* identifies sequences that determine recognitional specificity in Cf-4 and Cf-9. The Plant Cell, 9(12), 2209–2224.

Thomma, B. P., Van Esse, H. P., Crous, P. W., & de Wit, P. J. (2005). *Cladosporium fulvum* (syn. *Passalora fulva*), a highly specialized plant pathogen as a model for functional studies on plant pathogenic Mycosphaerellaceae. Molecular Plant Pathology, 6(4), 379–393.

Van den Ackerveken, G., Van Kan, J. A., Joosten, M., Muisers, J. M., Verbakel, H. M., & de Wit, P. (1993). Characterization of two putative pathogenicity genes of the fungal tomato pathogen *Cladosporium fulvum*. Molecular Plant-Microbe Interactions, 6, 210–215.

Van Esse, H. P., Bolton, M. D., Stergiopoulos, I., de Wit, P. J., & Thomma, B. P. (2007). The chitin-binding *Cladosporium fulvum* effector protein Avr4 is a virulence factor. Molecular Plant-Microbe Interactions, 20(9), 1092–1101.

Van Kempen, M., Kim, S. S., Tumescheit, C., Mirdita, M., Lee, J., Gilchrist, C. L., Söding, J., & Steinegger, M. (2024). Fast and accurate protein structure search with Foldseek. Nature Biotechnology, 42(2), 243–246.

Videira, S., Groenewald, J., Nakashima, C., Braun, U., Barreto, R. W., de Wit, P. J., & Crous, P. (2017). Mycosphaerellaceae–chaos or clarity? Studies in Mycology, 87, 257–421.

Wang, A., Meng, F., Xu, X., Wang, Y., & Li, J. (2007). Development of molecular markers linked to *Cladosporium fulvum* resistant gene *Cf-6* in tomato by RAPD and SSR methods. HortScience, 42(1), 11–15.

Wang, T., Black, L., Hsieh, W., & Hanson, P. (1995). Inheritance of black leaf mold resistance in tomato. Euphytica, 86, 111–115.

Witek, K., Jupe, F., Witek, A. I., Baker, D., Clark, M. D., & Jones, J. D. (2016). Accelerated cloning of a potato late blight–resistance gene using RenSeq and SMRT sequencing. Nature Biotechnology, 34(6), 656–660.

Wubben, J. P., Eijkelboom, C. A., & de Wit, P. J. (1993). Accumulation of pathogenesis-related proteins in the epidermis of tomato leaves infected by *Cladosporium fulvum*. Netherlands Journal of Plant Pathology, 99, 231–239.

Zaccaron, A. Z., & Stergiopoulos, I. (2020). First draft genome resource for the tomato black leaf mold pathogen *Pseudocercospora fuligena*. Molecular Plant-Microbe Interactions, 33(12), 1441–1445.

Zaccaron, A. Z., Chen, L.-H., Samaras, A., & Stergiopoulos, I. (2022). A chromosome-scale genome assembly of the tomato pathogen *Cladosporium fulvum* reveals a compartmentalized genome architecture and the presence of a dispensable chromosome. Microbial Genomics, 8(4).

Zaccaron, A. Z., & Stergiopoulos, I. (2024). Analysis of five near-complete genome assemblies of the tomato pathogen *Cladosporium fulvum* uncovers additional accessory chromosomes and structural variations induced by transposable elements effecting the loss of avirulence genes. BMC Biology, 22(1), 25.

