## Supplementary material for "Rapid breakdown of *Cf-6*-mediated immunity in tomato through loss or mutation of the *Avr6* effector gene in *Fulvia fulva*": Figures S1-S9, Notes S1

**Supporting Information**

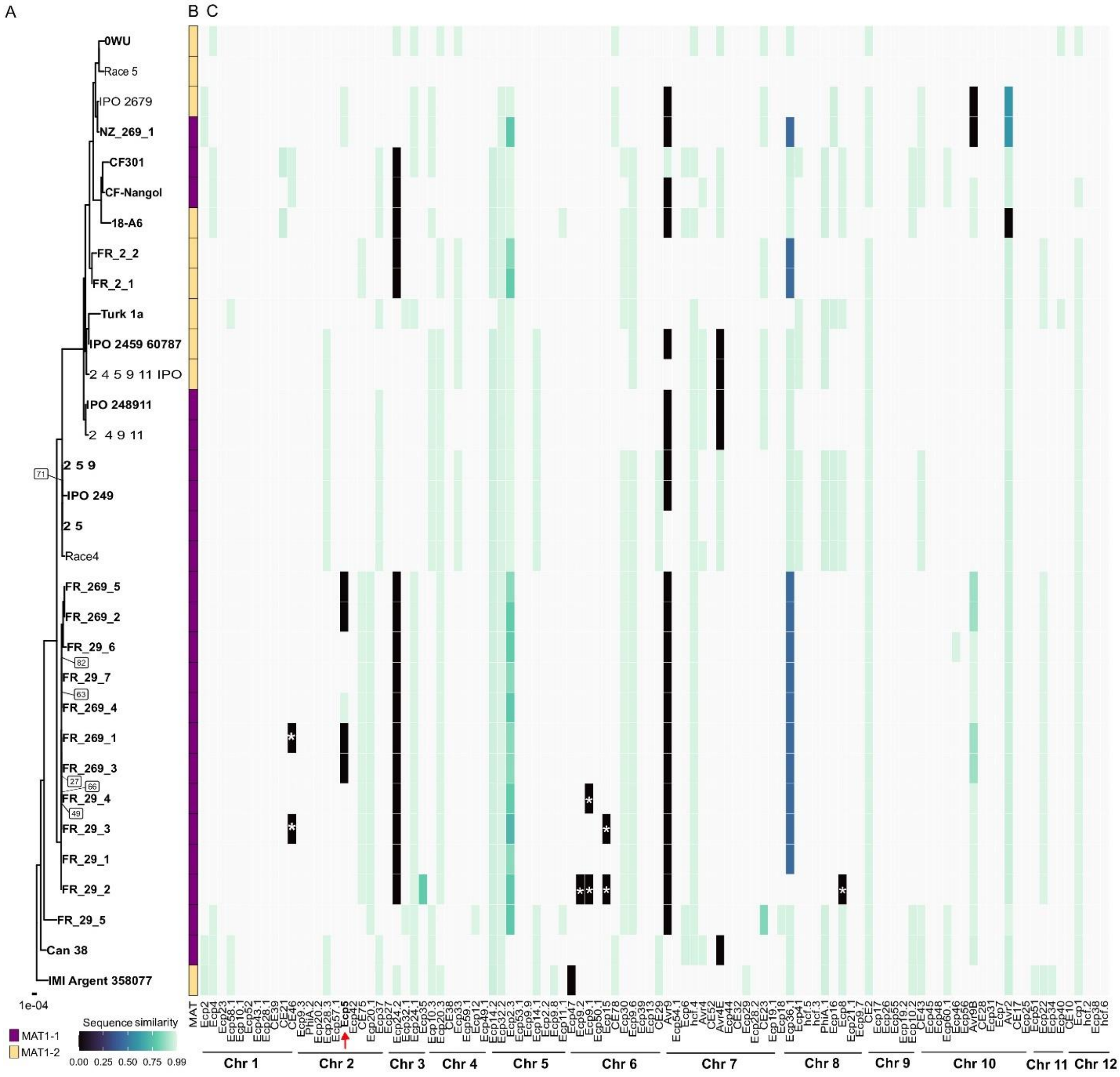


**Figure S1: Phylogeny and sequence comparison of effector genes across a panel of 33 *F. fulva* genomes.** A. Maximum-likelihood phylogeny based on a concatenated alignment of 3484 BUSCO genes, rooted with *Dothistroma septosporum* as outgroup (not shown in tree). Names of assemblies generated for this study are shown in bold. Only bootstrap values < 95 are shown in the tree. B. Genotypes of mating type (MAT) loci. C. Heatmap with sequence similarity to the Race 5 reference genome of known and predicted effectors in *F. fulva* strains. Identical sequences are indicated in white, sequences that are absent in black, and shades of green and blue indicate intermediate levels of sequence divergence. Sequences for which we found no hit in the assembly, but did find reads mapped to the reference genome at the effector locus, are indicated with an asterisk (*). *Ecp5*, the candidate for *Avr6*, is highlighted with a red arrow. Effectors are ordered according to their chromosomal location within the Race 5 genome.

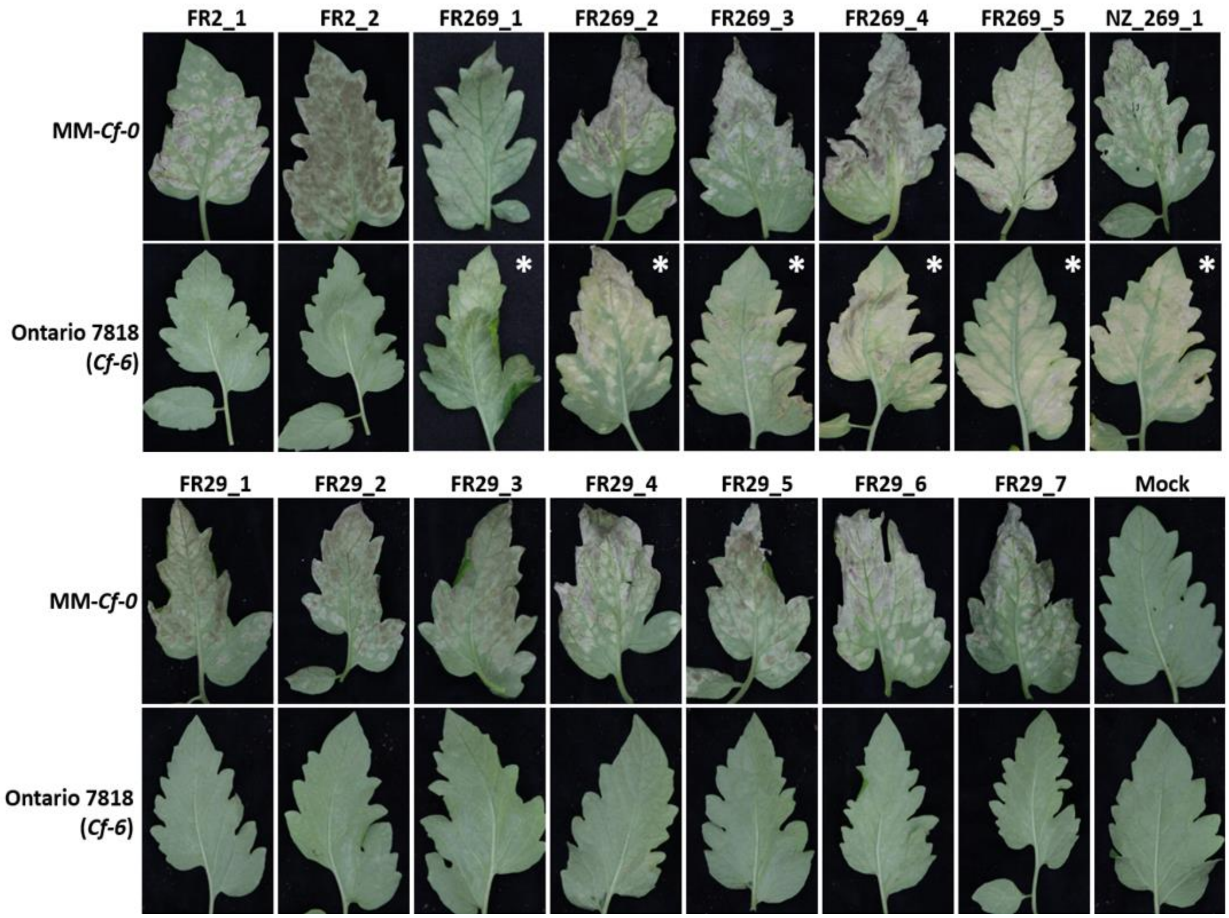


**Figure S2: Novel strains of *F. fulva* can infect Ontario 7818 tomato plants harboring the *Cf-6* gene.** *F. fulva* strains sequenced in this study were inoculated on 4-week-old Moneymaker (MM)-*Cf-0* (no *Cf* genes) and Ontario 7818 (*Cf-6*) tomato plants. At 14 days post-inoculation, representative leaves were photographed from the lower (abaxial) side. Asterisks indicate a compatible interaction on *Cf-6* tomato plants.

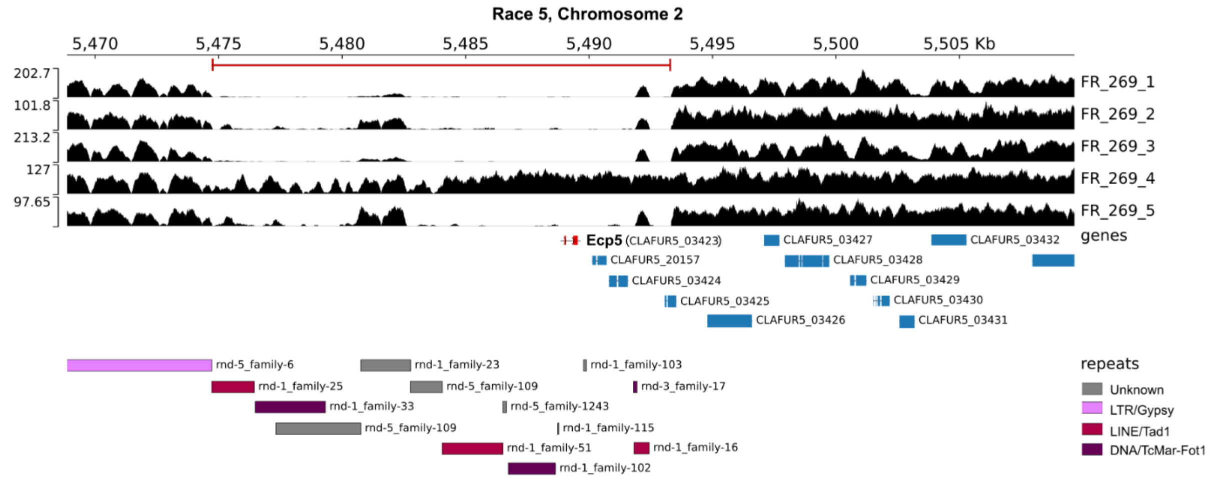


**Figure S3: *Ecp5* is part of an 18 kb deletion in four Race 6 strains of *F. fulva*.** Coverage plots of paired-end Illumina sequencing reads of five closely related Race 2.6.9 strains (of which four (FR_269_1m FR_269_2, FR_269_3 and FR_269_5) are predicted to have lost *Ecp5*), mapped to the genome of Race 5 (GCA_020509005.2). One strain (FR_269_4) has a non-synonymous mutation in *Ecp5.* In the top panel, read depth is shown in black, and only primary alignments are included. The predicted 18 kb deletion is shown with a red line at the top of the panel. The middle panel shows genes that are predicted in this region, with *Ecp5* shown in red and other genes in blue. The bottom panel shows transposable elements that are predicted by Zaccaron et al. (2022), with known superfamilies indicated in shades of pink and purple, and unknown superfamilies indicated in grey.

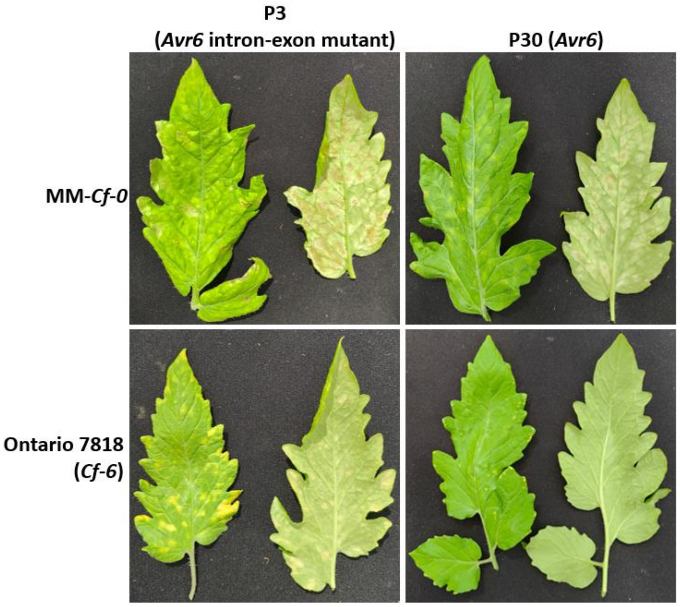


**Figure S4: An intron-exon border mutation in *Avr6* allows *F. fulva* to partially overcome *Cf-6*-mediated resistance.** Five-week-old MM-*Cf-0* (no *Cf* genes) and Ontario 7818 (*Cf-6*) tomato plants were inoculated with *F. fulva* strains P3 (*Avr6* intron-exon mutant) and P30 (wild-type *Avr6*). Leaflets were photographed at 14 days post-inoculation.

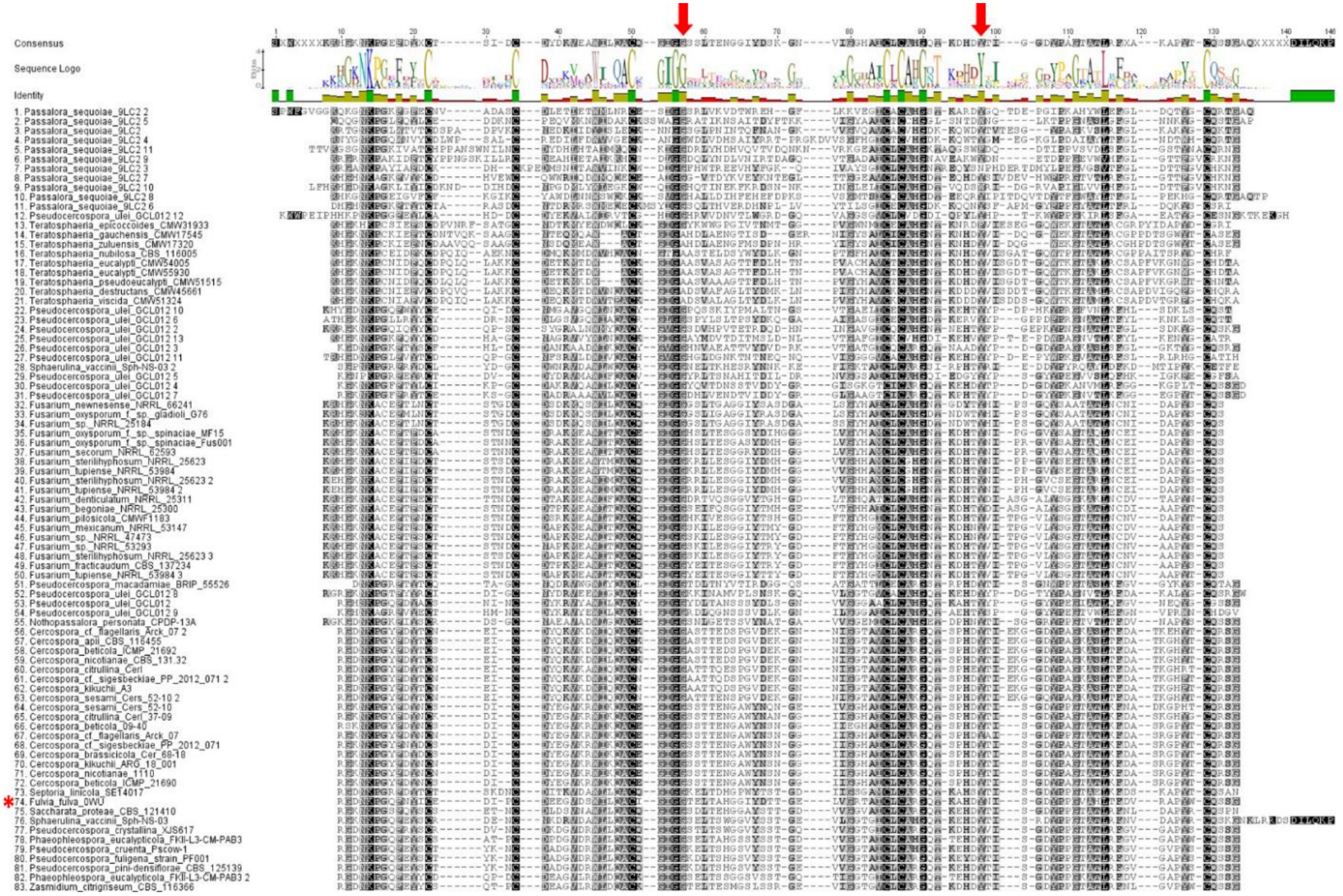

**Figure S5: Avr6-like proteins are predicted to be produced by many plant-pathogenic Dothideomycete and Sordariomycete fungi.** Only the mature sequence of the Avr6-like proteins (i.e. without their signal peptide for extracellular targeting) is aligned. Avr6 from *F. fulva* is indicated with a red asterisk. The mutated, conserved non-cysteine residues in Avr6 of *F. fulva* strains NZ_269_1 (Tyr88) and FR_269_4 (Gly53) are indicated with a red arrow on top of the alignment.

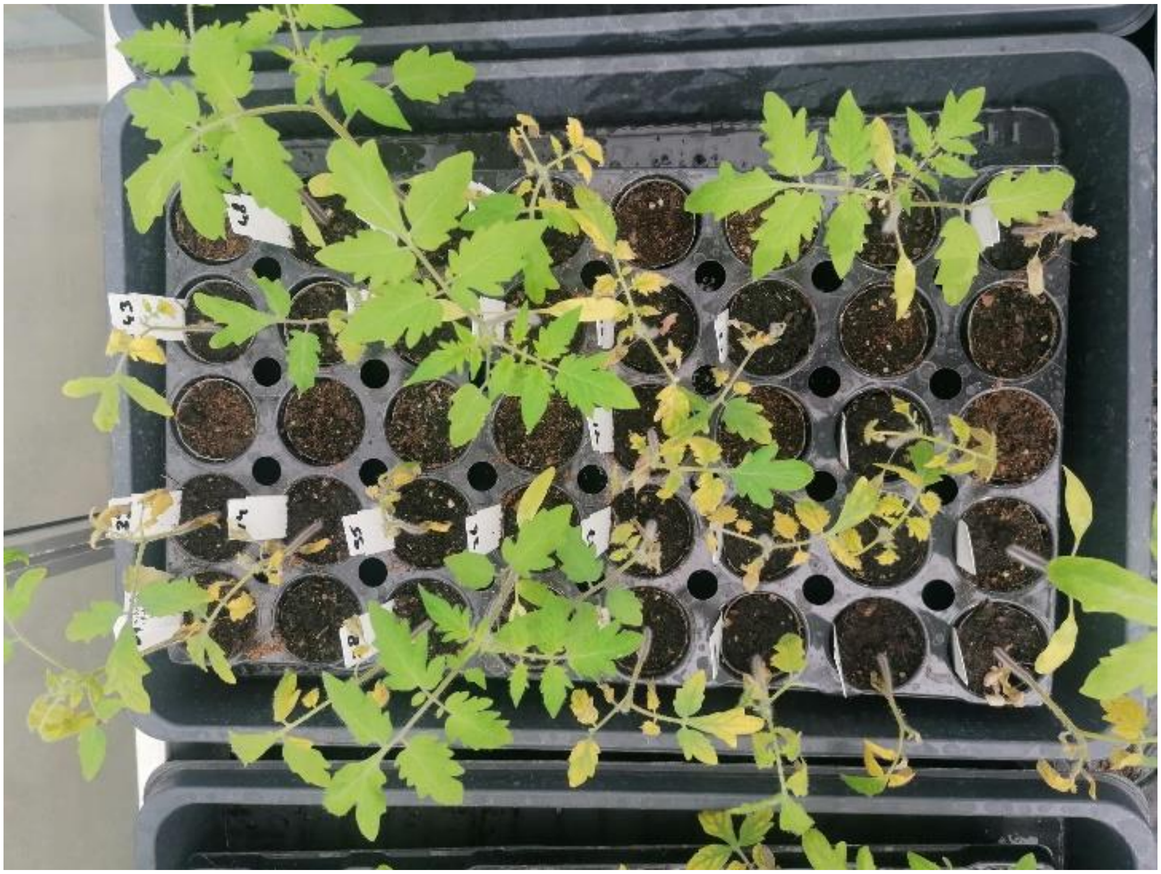

**Figure S6: The Ontario 7818 x MM-*Cf-0* F2 BSA mapping population segregates for an HR to Avr6.** Representative picture of tomato plants after systemic expression of PVX::*Avr6* mapping population. The *Agrobacterium* strain harboring PVX::*Avr6* was syringe-infiltrated in cotyledons of 16-day-old plants. Avr6-responsive plants show severe necrosis, chlorosis and stunting, whilst non-responsive plants remain unaffected. Photograph was taken at 10 days post-inoculation.

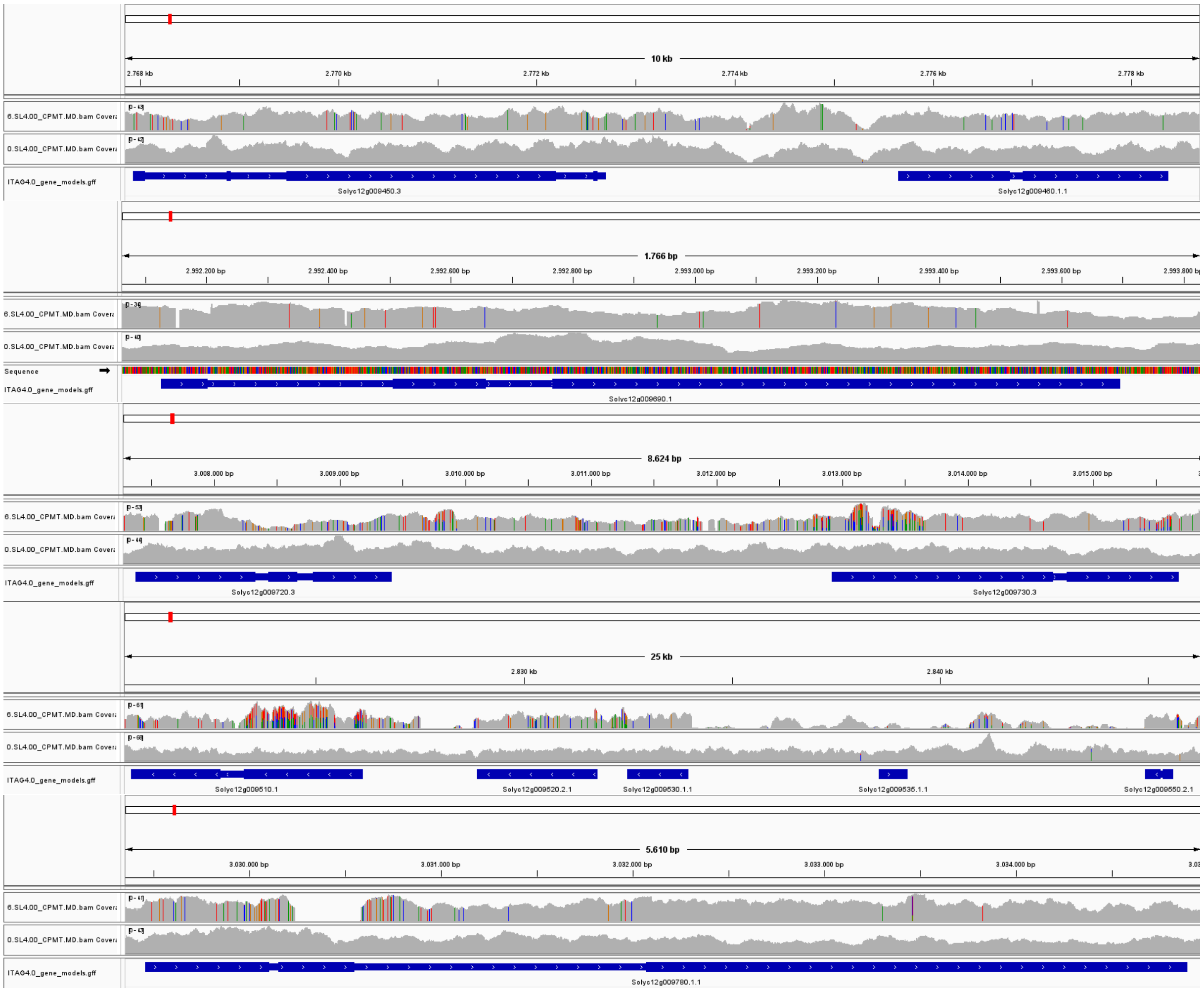

**Figure S7: Receptor-encoding candidates within the mapping interval of *Cf-6* exhibit significant numbers of R-pool-specific SNPs compared to the Heinz 1706 SL4.0 reference genome and the NR-pool.** Figures show coverage tracks of reads of the R- and NR-pools (top and bottom, respectively), visualized using the integrative genome viewer (IGV). Colored bars indicate positions with polymorphisms compared to the reference. The ITAG4.0 gene annotation is shown in blue, with the intron/exon structure shown as narrow and wide regions, respectively.

Solyc12g009450.3.1 NBS-LRR protein (AHRD V3.3 *** Q9SM52_SOLAC)

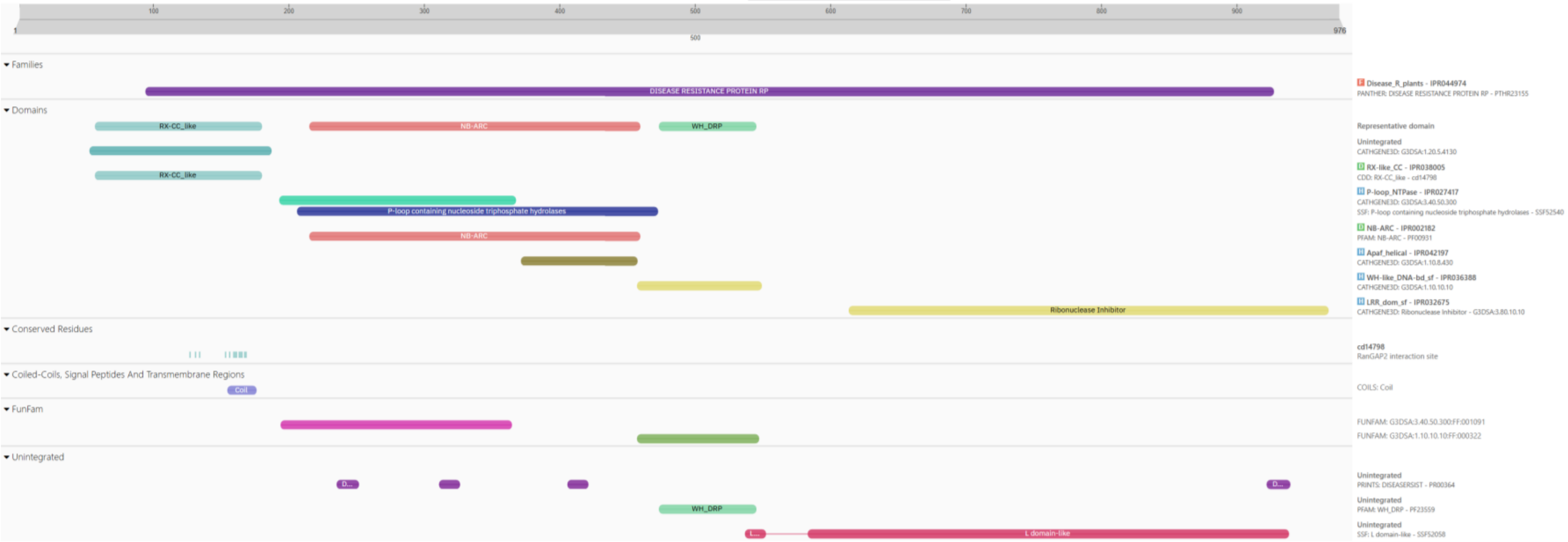


**Figure S8: Domain predictions of receptor proteins within the mapping interval of *Cf-6*, based on the Heinz 1706 reference annotation (ITAG 4.0).** Specific protein domains are indicated by colored bars, and their full descriptions can be found on the left. Predictions and visualizations were made using InterPro.

Solyc12g009460.1.1 Disease resistance protein (CC-NBS-LRR class) family (AHRD V3.3 *** AT5G35450.2)

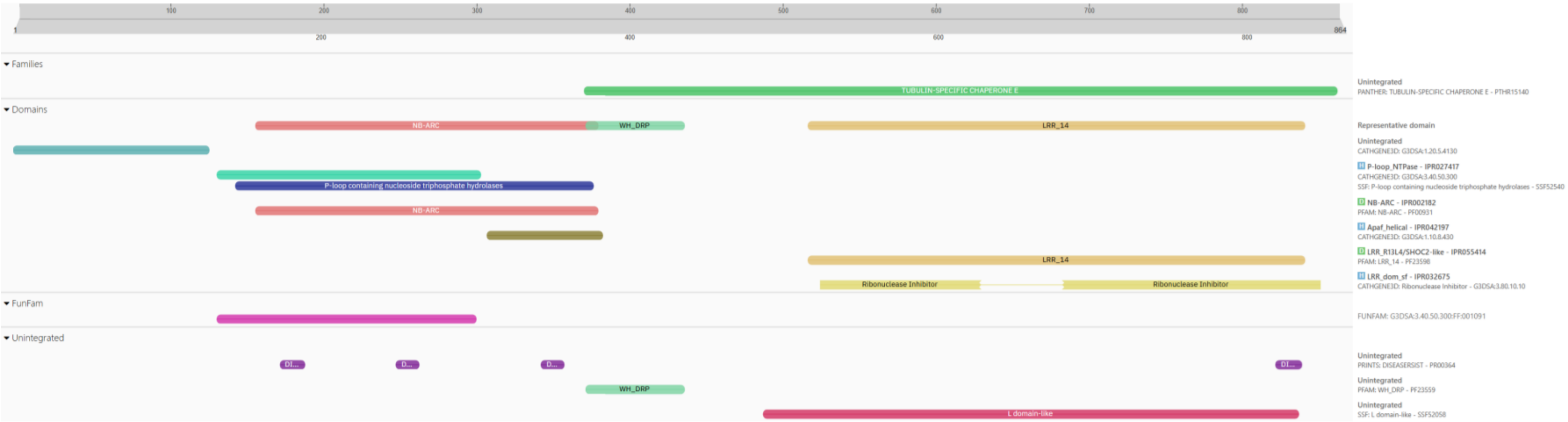

**Figure S8** (continued)

Solyc12g009510.1.1 receptor-like protein 12 (AHRD V3.3 *** A0A1S4DP57_TOBAC)

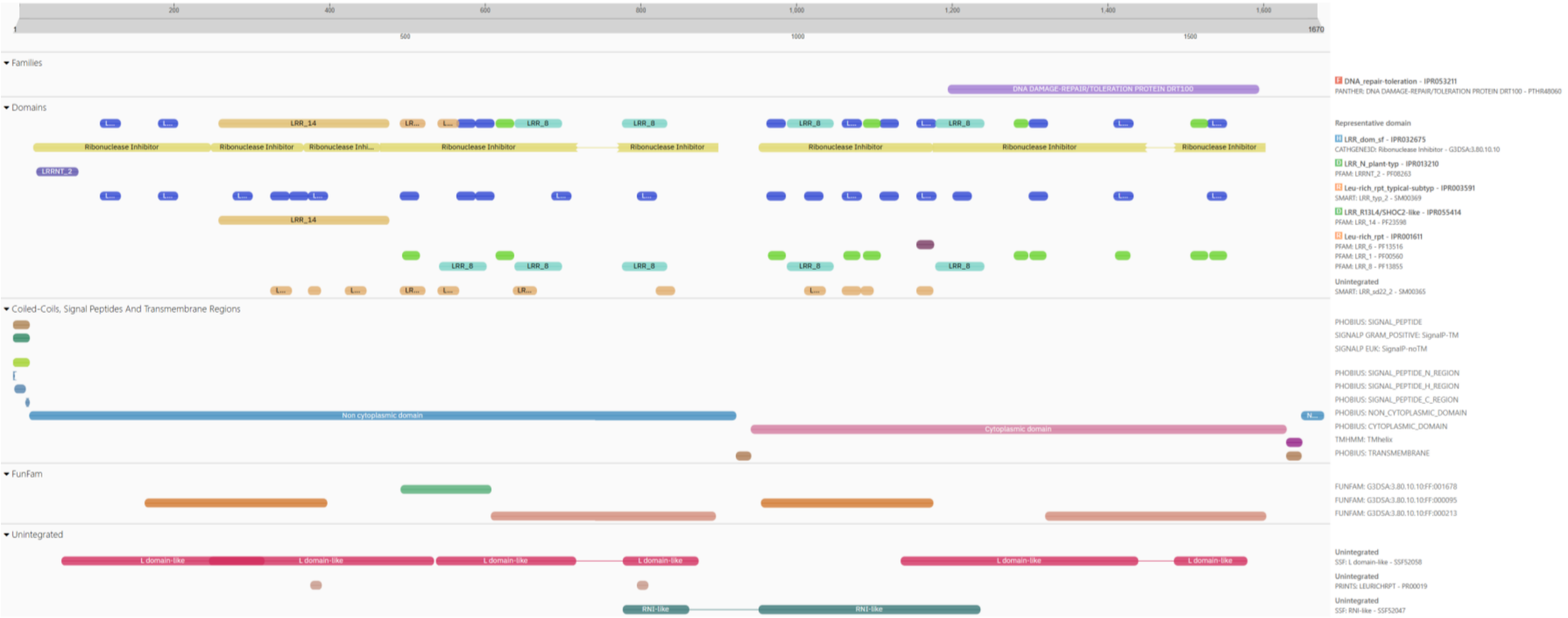


**Figure S8** (continued)

Solyc12g009520.2.1 receptor-like protein 12 (AHRD V3.3 *** A0A1S4DP57_TOBAC)

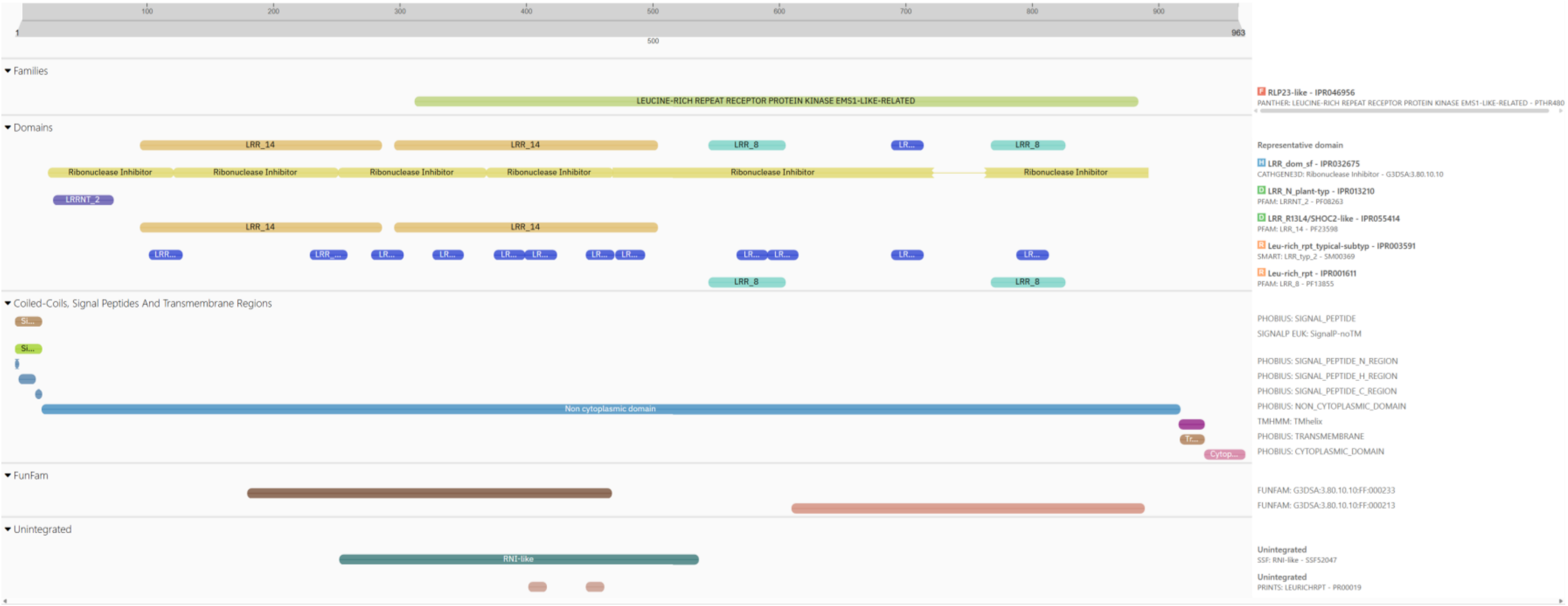


**Figure S8** (continued)

Solyc12g009530.1.1 receptor-like protein 12 (AHRD V3.3 *-* A0A1S4DP57_TOBAC)

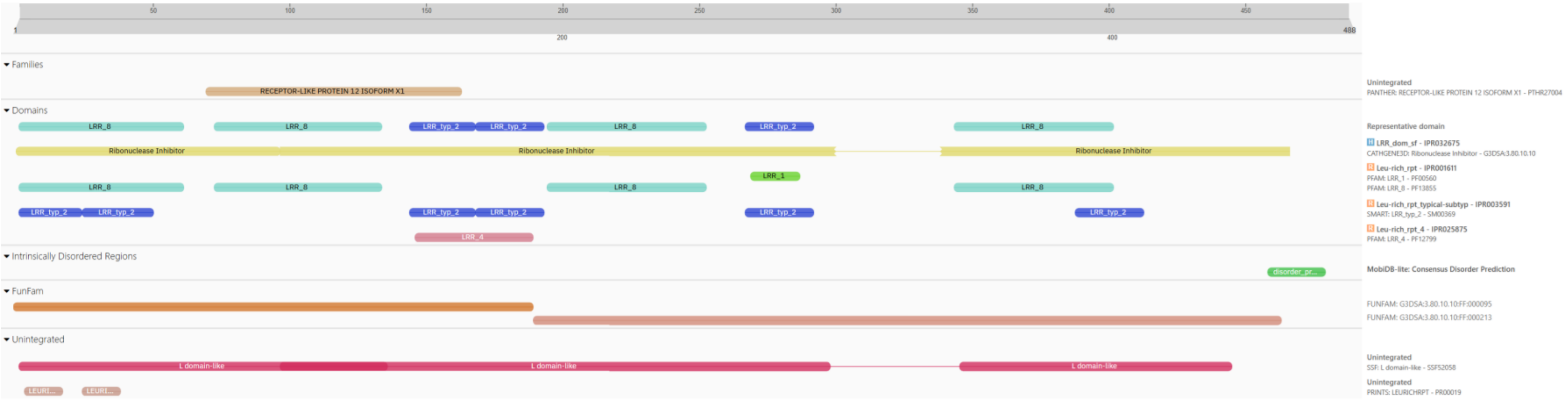


**Figure S8** (continued)

Solyc12g009550.2.1 Leucine-rich repeat receptor-like protein kinase PXL1 (AHRD V3.3 *-* A0A1D6GYD2_MAIZE)

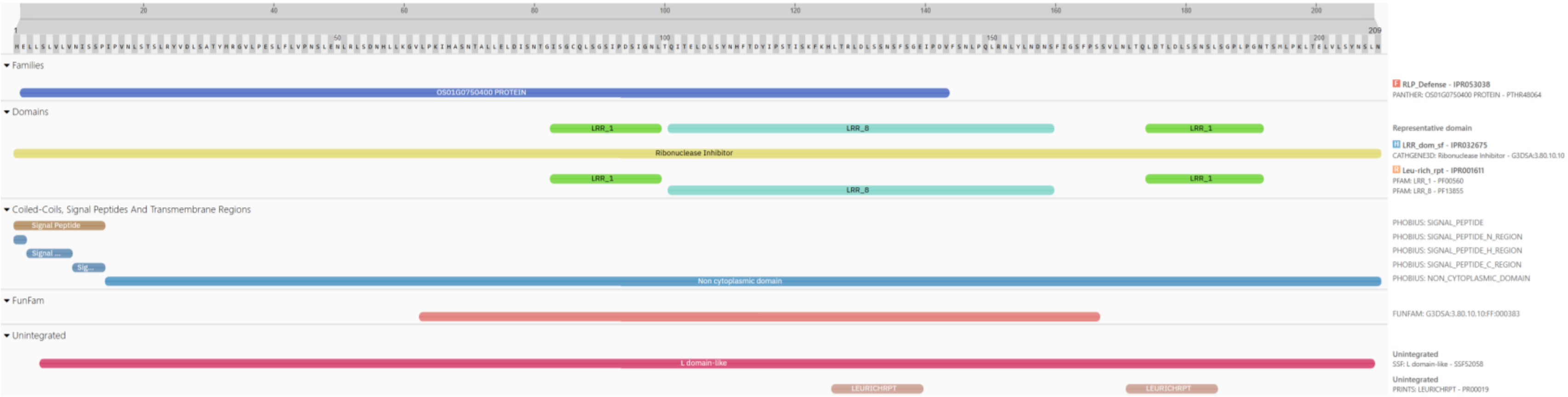


Solyc12g009690.1.1 receptor-like protein 12 (AHRD V3.3 *-* A0A1S4DP57_TOBAC)

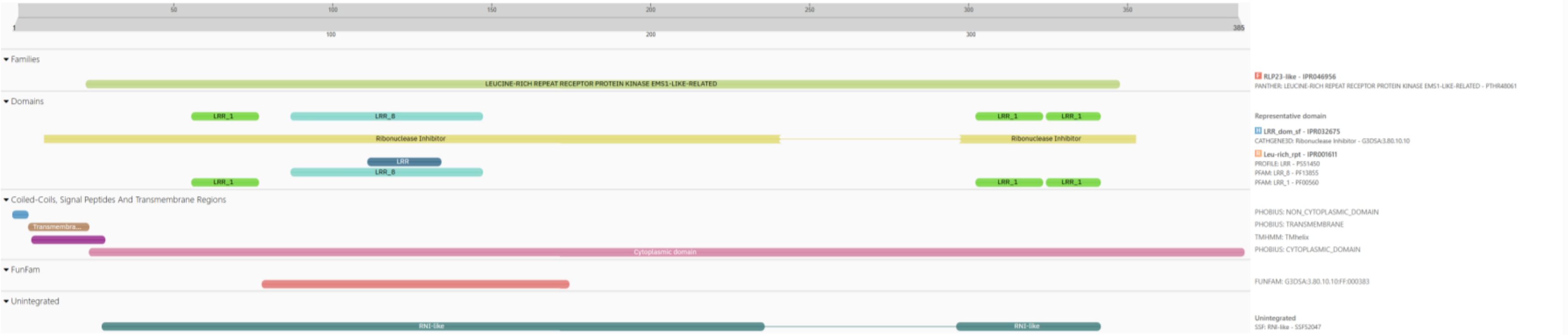


**Figure S8** (continued)

Solyc12g009720.3.1 receptor-like protein 12 (AHRD V3.3 *** A0A1U8E178_CAPAN)

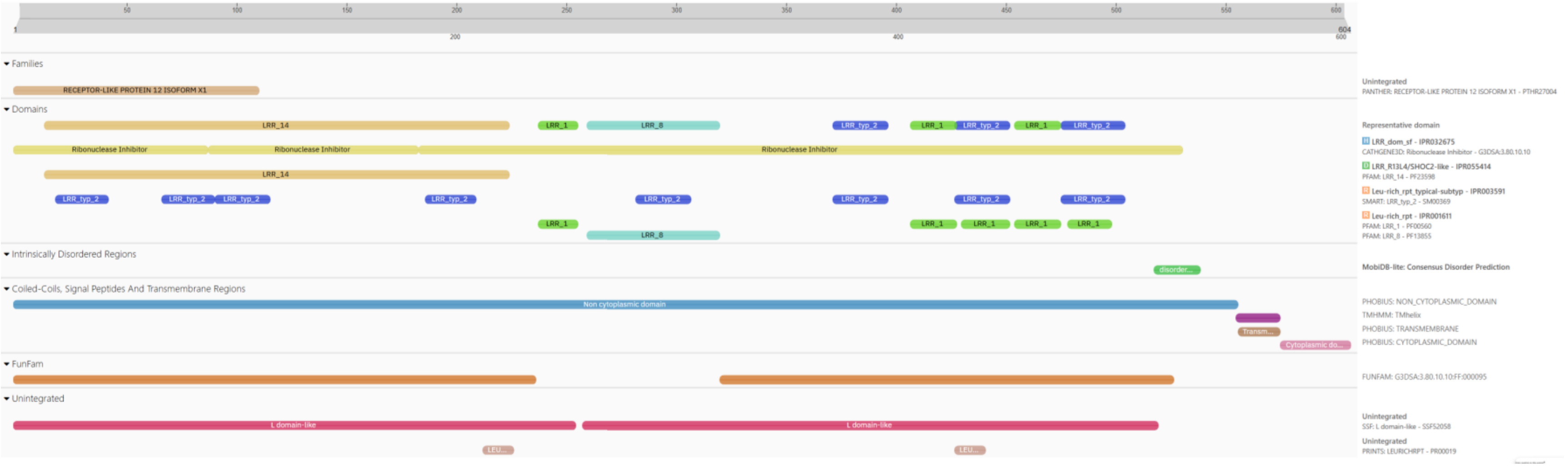


**Figure S8** (continued)

Solyc12g009730.3.1 receptor-like protein 12 (AHRD V3.3 *** A0A1S4DP57_TOBAC)

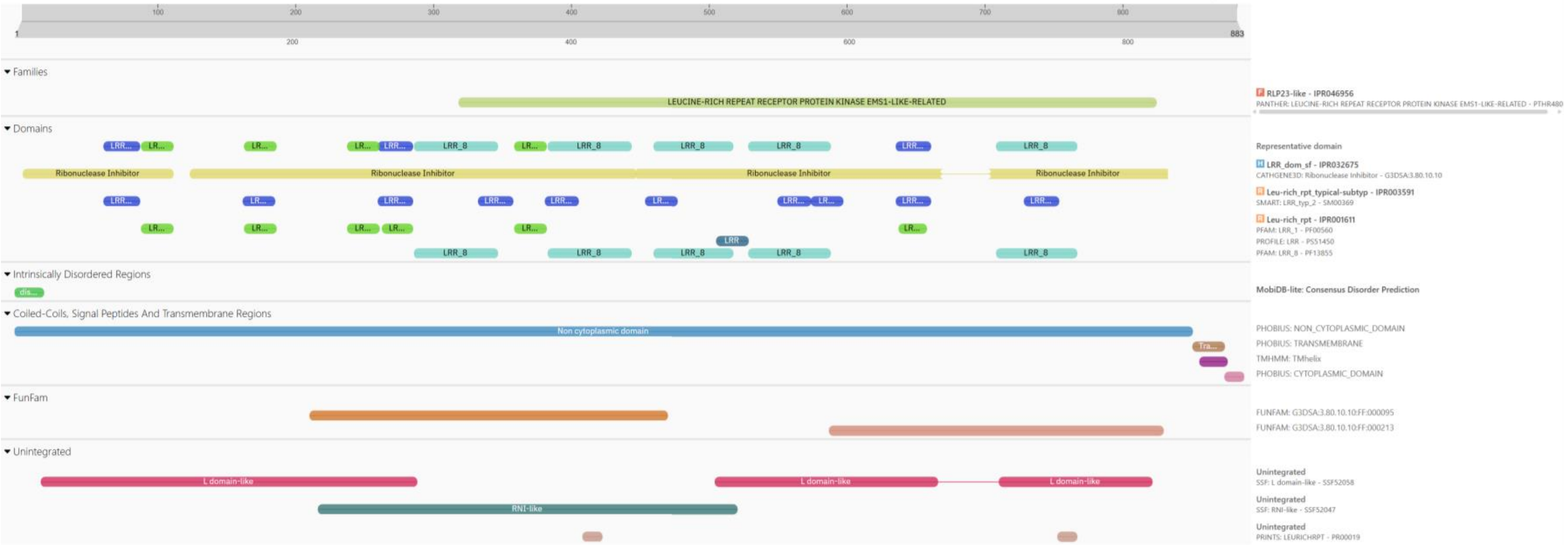


**Figure S8** (continued)

Solyc12g009780.1.1 receptor-like protein 12 (AHRD V3.3 *** A0A1S3Y203_TOBAC)

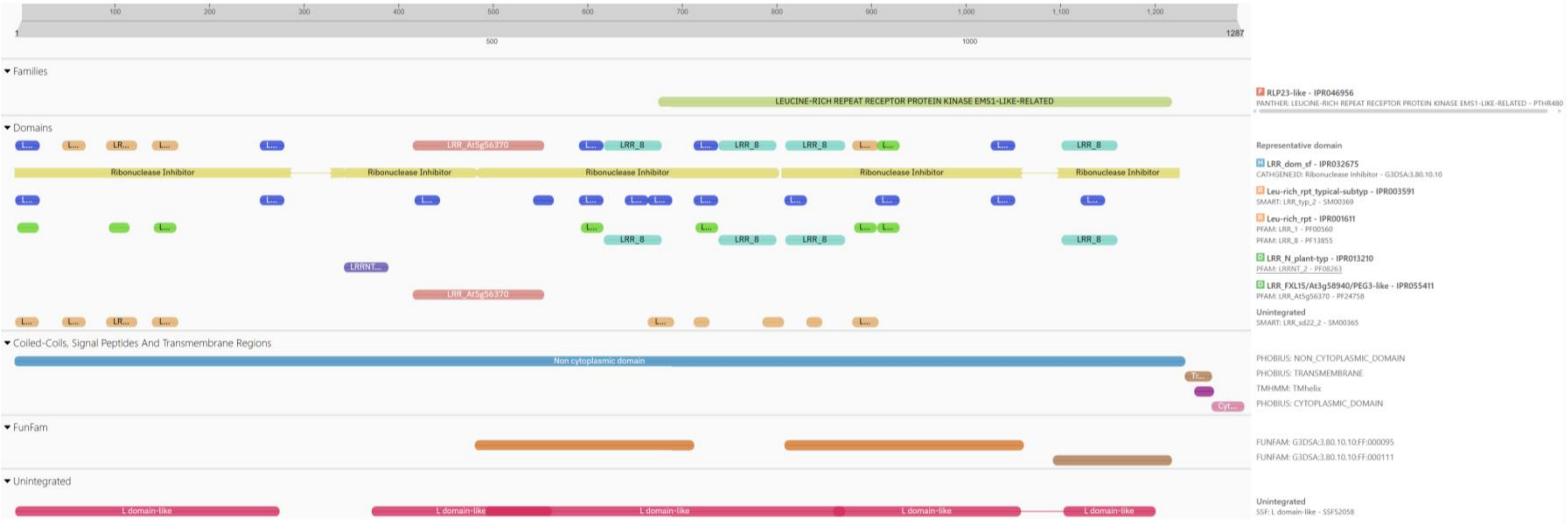


**Figure S8** (continued)

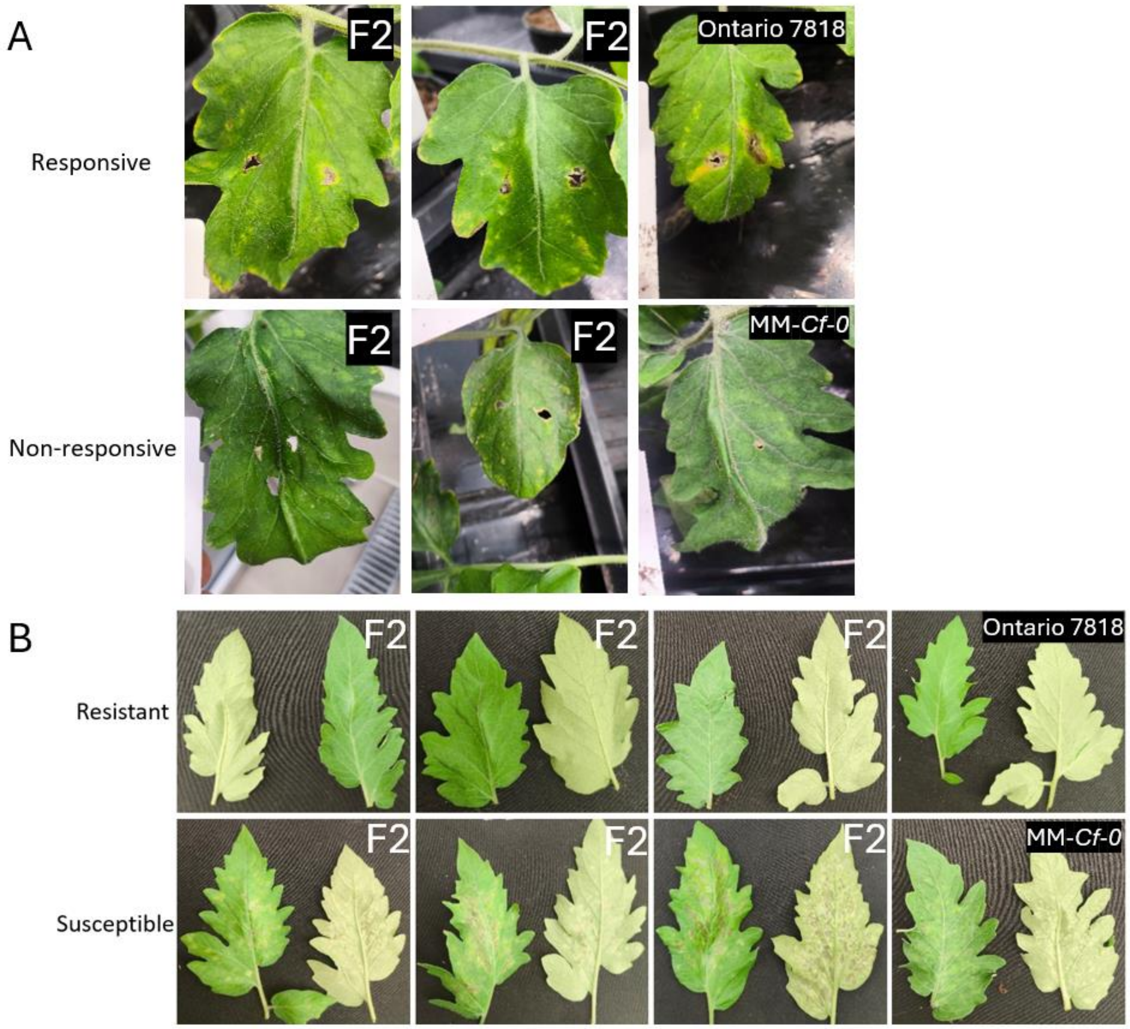


**Figure S9: Recognition of Avr6 correlates with tomato leaf mold disease resistance in Ontario 7818 x MM-*Cf-0* F2 individuals.** Representative pictures of disease symptoms after *F. fulva* inoculation and response to Avr6. Four-week-old plants were inoculated by means of toothpick inoculation of *Agrobacterium* carrying PVX::*Avr6* (Panel A). Photographs of representative F2 individuals, and MM-Cf-*0* and Ontario 7818 plants (as controls) were taken at 20 dpi. Before PVX::*Avr6* inoculation, cuttings were taken of each plant and when these were 20 days old, they were inoculated with *F. fulva* race 0 strain 0WU (Panel B). Photographs of disease symptoms were taken of representative F2 individuals, and MM-Cf-*0* and Ontario 7818 plants (as controls) at 14 days post-inoculation.

**Notes S1: Predicted Avr6-like sequences from fungi.** Predicted Avr6-like protein sequences are shown together with their corresponding gene sequences. Signal peptides are underlined. Introns are highlighted red.

>*Fulvia_fulva*_0WU_Avr6
MNTFTLLLATLPLLAYGRGDNKPGQFNYICEDIPCDEEGVDSWILQACIGIGGTELTAHGGIYDTTGELVGRTAICLCARGQTKAHDYNISGDYPPG
VAHLTFDVRAPYWCQSSG*
ATGAGCAAGTCCCGCTACATCCAACCCTTTCATATGCTCTCTAACATCCTAGCAGACACTTTTACTCTACTCCTGGCCACACTCCCACTGCTGGCAT
ATGGCAGGGGCGACAATAAGCCCGGCCAGTTCAATTACATTGTAAGTGGGTCTTCGACGCTGGATCTGCCCATACCTCAATGTCCTTTGTCACAACC
TCTCTCGCTAGAAGTCAACGGTGCCCTCGCGCGACGCGTGGCATATAGCAAACAGTGTGAGGCATCAGCGCATCGGTCCATGCTCTTTGGTCGTTGT
CAGCTCAGCGGCCCTATCGCGAATTCGGCAGCATCGACATCAGGGCGGATGGTGAGAGGCGAGTCGAGAAACGAAACATGTCACCAGACTTGCCCGT
TTCTTATGCATGAGCCGGCATTATCTCGAGATGGCGGTTCCATCTCGAGTCAGCCGCCTCACAGCAGAAACCTAGATCATTCTACCGAGAGACGTTC
CCTATTCTCTTTCCCTCTGCTCAAACACTGTACTGATTCTCTATCACTAGTACGAAGATATCCCTTGCGACGAAGAGGGGGTCGACAGCTGGATTCT
GCAAGCATGCATCGGAATTGGAGGGACTGAGCTTACAGCTCATGGAGGGATATACGACACGACTGGAGAACTCGTTGGCCGGACGGCGATCTGCTTG
TGTGCAAGGGGCCAGACGAAGGCGCATGATTATAACATCAGTGGGGATTACCCGCCAGGGGTTGCGCATTTGACGTTTGATGTTCGGGCGCCTTACT
GGTGTCAGAGTTCTGGATAG
>*Phaeophleospora_eucalypticola*_FKII-L3-CM-PAB3
MKFLVLLSLLLPALTSARGDNKPGQFEYSCSATPCDPDGVDRWLLQACQGIGGSELTAHGSSYSSTGDVVGGTAICLCAHGQTTEHDYTISGDYPPG
TATLRFGVGAPYWCQSSG*
ATGAGTGAGTCAACGTATCTTTTTCTCTGATTCCACGCTTCTAACTATGCCCTCATAGAGTTCCTCGTTCTGCTCTCCCTCCTCCTCCCGGCCCTCA
CCTCCGCCCGCGGCGACAACAAACCGGGCCAGTTCGAGTACTCCTGCAGCGCAACTCCATGCGACCCCGATGGCGTCGACCGCTGGCTGCTCCAGGC
TTGTCAAGGCATCGGCGGCAGTGAACTGACAGCCCATGGATCGAGTTACAGCTCGACGGGTGATGTGGTGGGCGGTACGGCGATTTGTTTATGTGCA
CATGGACAGACGACAGAGCATGATTATACGATTTCGGGTGATTATCCGCCGGGGACTGCGACGTTGCGGTTTGGGGTGGGGGCGCCGTATTGGTGTC
AGAGTAGTGGGTAG
>Pseudocercospora_pini-densiflorae_CBS_125139
MNAILIIPLLLTTLVSGRGDNKPGQFEYSCTYKNCDADGVDRWLLQACQGIGGKDLTSHGSVYSSTGEVVGGTAICLCARGQTKEHDYTISGDYPPG
TATLRFGVGAPYSCQSSG*
ATGAGTAACTCGCCTCCCTCTTTCCATCTCATTTCGGTCCTAACCAGTATCCAGATGCCATCCTCATCATTCCCCTCCTCCTAACAACCCTAGTCTC
AGGGCGAGGCGACAACAAACCAGGTCAATTCGAATACAGCTGCACATACAAGAATTGCGACGCAGATGGAGTTGACAGATGGCTTCTTCAAGCTTGC
CAAGGTATCGGCGGCAAAGATCTCACTTCGCATGGCAGTGTGTATAGTTCAACCGGCGAAGTCGTAGGCGGGACTGCGATTTGTTTGTGTGCTCGTG
GTCAGACCAAAGAACACGATTATACGATTTCGGGAGACTACCCTCCTGGAACGGCAACTTTGAGATTTGGTGTTGGGGCGCCGTATTCGTGTCAGAG
TTCGGGATGA
>Pseudocercospora_fuligena_strain_PF001
MNLILILPLLLTTLVSARGDNKPGQFEYSCTYKNCDADGVDRWLLQACQGIGGKDLTSHGSVYSSTGEVVGGTAICLCARGQTKEYDYTISGDYPPG
TATLRFDVGAPYSCQSSG*
ATGAGTAAACCGCCTCCTCTTTCCACCTTGATTTATCCAACATTGACCACGTTCAGATCTTATCCTCATCCTTCCCCTCCTCCTCACAACACTAGTC
TCAGCCCGAGGTGACAACAAACCAGGCCAATTCGAATACAGCTGTACATACAAGAACTGCGACGCAGATGGTGTCGACAGATGGCTGCTTCAAGCTT
GCCAAGGTATCGGCGGCAAAGATCTCACATCACACGGCAGTGTGTATAGTTCAACCGGCGAAGTCGTAGGCGGGACTGCGATTTGTTTGTGTGCTCG
TGGTCAGACCAAAGAATATGATTATACGATTTCGGGAGACTATCCTCCTGGAACGGCCACTTTGAGGTTTGATGTTGGGGCGCCGTATTCGTGTCAG
AGTTCGGGATGA
>Pseudocercospora_crystallina_XJS617MKLLSLVATATLLSLASARGDNKPGQFRYSCRDVNCDKDGVDRWLLQACQGIGGSTLTSWG
GVYSSTGQVVGGTGICLCARGQTKDHDYVISGDYPPGTATLRFGVGAPYSCQSSG*
ATGAGTACGACATGCCCATGATAACGAACACATCAAACATACTAACCCTCATCCAAAGAACTCCTCTCCCTCGTCGCTACCGCCACCCTCCTCTCCC
TCGCCTCCGCCCGCGGCGACAACAAACCAGGCCAATTCCGCTACAGCTGCAGAGATGTCAACTGCGACAAAGACGGCGTCGATCGATGGCTCTTACA
AGCCTGTCAAGGCATCGGAGGAAGTACCTTGACTTCTTGGGGTGGAGTCTACAGCTCAACTGGACAGGTCGTGGGAGGAACTGGGATTTGTCTTTGT

GCGAGGGGCCAGACGAAAGATCACGATTACGTGATCTCGGGCGATTATCCGCCGGGGACGGCGACGTTGAGGTTTGGGGTGGGGGCGCCTTATAGTT

GTCAGAGCTCGGGATAG

>Pseudocercospora_cruenta_Pscow-1

MNPILIISLLLTALVSARGDNKPGQFEYSCTYKNCDADGVDRWLLQACQGIGGKELTSHGSVYSSTGEVVGGTAICLCAHGQTKEHDYTISGDYPPG

TATLRFGVGVPYSCQSSG*

ATGAGTGAACCATCTCCTCTTTCCACCTTGATTGATCCAGAACTAACAACATTCCAGATCCCATCCTCATCATTTCCCTCCTCCTCACAGCCCTGGT

CTCAGCCCGAGGCGACAACAAGCCAGGCCAATTCGAATACAGCTGCACATACAAAAACTGTGACGCAGATGGTGTCGACAGATGGCTGCTTCAAGCT

TGCCAAGGTATCGGCGGCAAAGAACTCACATCACACGGCAGTGTGTATAGTTCAACCGGCGAAGTCGTAGGCGGGACTGCGATTTGTTTGTGTGCTC

ATGGCCAGACCAAAGAACACGATTATACGATTTCAGGCGACTATCCTCCTGGAACGGCCACTTTGAGGTTTGGTGTTGGGGTGCCGTATTCGTGTCA

GAGTTCGGGTTGA

>Zasmidium_citrigriseum_CBS_116366

MQLLSLVPALMALTPLALARGDNKPGQFEYYCSQTPCDEAGVLRWLDQACKGIGGTELTESMGSLSSRGEVIGGTGVCLCARGQTKDHDYTISGDYP

PGTATLRFNLGVPYWCQSSG*

ATGCGTAAGCCGAGCCACTCTCACAACTCCAAACAACGCCCTCTGACAACCCAAACAGAGCTGCTCTCCCTCGTTCCCGCCCTGATGGCCCTCACAC

CCCTCGCCCTCGCCCGCGGCGACAACAAACCCGGCCAATTCGAATACTACTGCTCACAAACGCCCTGCGACGAGGCCGGCGTCCTGCGCTGGCTGGA

CCAGGCCTGCAAAGGTATCGGTGGAACTGAATTGACGGAGTCGATGGGCAGCCTGAGCTCGAGGGGAGAAGTCATCGGCGGGACGGGCGTTTGTCTT

TGTGCGCGTGGCCAGACCAAGGACCATGATTATACCATTTCGGGCGATTACCCACCGGGTACTGCGACGTTGAGATTCAATTTGGGGGTGCCGTATT

GGTGTCAGAGCTCGGGGTAG

>Saccharata_proteae_CBS_121410

MKISLAFFLFLASASARGKNKPGSFQYSCSETNCDLDSVNAWLLQACQGIGGSDLTEHGGAYSTTGDVVGGVAICLCARGQTKEHDYTISGDYPPGT

ATLRFNLDAPYRCQSPN*

ATGAGTGAGTTTCGGACGACAGCTCTAATAAGCGATGATGACTAATCAGAACATCTTTCCAGAGATATCTCTCGCCTTCTTCCTCTTCCTCGCGTCT

GCCTCCGCACGGGGGAAGAACAAACCAGGCTCGTTTCAGTACAGCTGCAGCGAGACAAACTGCGACCTGGACAGCGTCAATGCATGGCTCCTTCAAG

CCTGTCAAGGAATTGGAGGCAGTGATTTGACTGAGCACGGGGGTGCCTACAGCACGACTGGTGATGTGGTGGGTGGGGTTGCGATATGCTTGTGCGC

GAGAGGGCAGACGAAGGAGCATGATTACACCATCAGTGGAGATTATCCTCCAGGGACAGCGACGCTGAGATTCAATCTCGACGCGCCATATCGGTGC

CAAAGTCCGAACTAA

>Sphaerulina_vaccinii_Sph-NS-03

MKLILPIFALLTLASARGKNKPGQFEYSCRDENCNKDRVDAWLLQACQGIGGSELTSWGGVYSSTGEVLGGTGICLCAHGQTKEHDYTISRDYPPGT

ATLRFNVDAPYRCQSKGNKLRRDSDILQKP*

ATGAGTAAGCCAGACTGCTGGTGATGTATATAAAACAGACTACTGACACTCTCCAGAGCTCATACTACCAATCTTCGCCCTTCTGACCCTGGCATCC

GCACGTGGCAAGAACAAGCCTGGCCGTATGTCTTTCAGAAAGCCTTCCATACTTCAGCACTGACGCATTCGCCAGAATTCGAATACAGCTGCAGAGA

TGAGAACTGCAACAAGGATAGAGTAGACGCCTGGTTGCTTCAGGCTTGCCAAGGAATCGGCGGCAGCGAACTCACATCGTGGGGTGGTGTCTACTCG

TCAACAGGCGAAGTCTTGGGCGGCACTGGTATCTGTCTATGCGCGCATGGTCAAACTAAAGAACACGACTACACCATCAGCCGCGACTATCCTCCAG

GAACGGCCACGTTACGATTCAATGTCGATGCACCTTATAGATGTCAGAGTAAGGGCAACAAGTTGCGCCGCGATAGTGATATTTTACAGAAGCCATA

G

>Sphaerulina_vaccinii_Sph-NS-01

MKLILPIFALLTLASARGKNKPGQFEYSCRDENCNKDRVDAWLLQACQGIGGSELTSWGGVYSSTGEVLGGTGICLCAHGQTKEHDYTISRDYPPGT

ATLRFNVDAPYRCQSKGNKLRRDSDILQKP*

ATGAGTAAGCCGAACTCCGGATGTTATAGCATAAGACGGACTACTGACACTCTCCAGAGCTCATACTACCAATCTTCGCCCTTCTGACCCTGGCATC

CGCACGTGGCAAGAACAAGCCTGGCCGTATGTCTTTCAGAAAGCCTTCCATACTTCAGCACTGACGCATTCGCCAGAATTCGAATACAGCTGCAGAG

ATGAGAACTGCAACAAGGATAGAGTAGACGCCTGGTTGCTTCAGGCTTGCCAAGGAATCGGCGGCAGCGAACTCACATCGTGGGGTGGTGTCTACTC

GTCAACAGGCGAAGTCTTGGGCGGCACTGGTATCTGTCTATGCGCGCATGGTCAAACTAAAGAACACGACTACACCATCAGCCGCGACTATCCTCCA

GGAACGGCCACGTTACGATTCAATGTCGATGCACCTTATAGATGTCAGAGTAAGGGCAACAAGTTGCGCCGCGATAGTGATATTTTACAGAAGCCAT

AG

>Phaeophleospora_eucalypticola_FKII-L3-CM-PAB3

MKQSILAMMALPFLFLARGDNKPGQFEYYCDQPDCDEAGVLRWLDQACKGIGGSELTSSGGSVSSTGQTIGGTGICLCARGQTTEHDYTISGDYPPG

KATLRFDVGAPYTCQSSG*

ATGAGTACGAGAGCAAGGTTTTATTCCAGAAAAATGCACTCACATGGTGGTAGAACAAAGCATACTGGCCATGATGGCTTTGCCCTTCCTTTTTTTA

GCCAGAGGAGACAACAAGCCCGGTCAATTCGAATACTATTGCGACCAGCCAGATTGCGATGAAGCTGGCGTACTCAGGTGGCTCGATCAAGCTTGCA

AAGGAATTGGCGGAAGTGAATTGACTTCTTCTGGTGGATCCGTCTCTTCAACAGGCCAGACAATTGGTGGCACTGGCATTTGCTTGTGCGCTCGCGG
ACAGACGACGGAGCACGACTATACGATCTCGGGCGATTATCCTCCAGGCAAGGCCACGCTGAGGTTCGACGTTGGAGCTCCGTACACGTGTCAAAGC
TCGGGATAG

>Septoria_linicola_SE14017

MNLIATLLVTILPLSLARGDNKPGRFDYTCTSKDNCDITKVDAWLLQACQGIGGKELTDHGPWYTSTGDIIGSGAMCLCARGQTKAHSYTISGDYPP
GTAELKFDSKAPYTCQSAN*

ATGAGTAAGCAGCCTCGATATCTGGAACGACAGCAGCTGGCTAACCTGACATAGATCTCATCGCGACACTCCTTGTGACCATACTGCCACTGTCGCT
GGCCCGTGGTGACAACAAACCTGGCCGTACGAAGCCTCGCTACTCCCACAGCACAACGGCCTACTGACAGCGTGCTCCTAGGATTTGACTACACCTG
CACAAGCAAAGACAACTGCGACATCACGAAGGTGGACGCTTGGCTACTACAGGCATGCCAAGGCATCGGAGGCAAGGAACTGACTGACCACGGGCCG
TGGTACACATCTACGGGCGACATTATCGGATCAGGAGCAATGTGTCTCTGCGCTAGAGGCCAGACCAAAGCGCACAGCTATACTATCAGTGGTGACT
ATCCGCCAGGAACGGCCGAGCTGAAATTCGATTCGAAAGCGCCATACACATGCCAAAGCGCGAATTGA

>Pseudocercospora_ulei_GCL012

MKFTTYSYILLTWAVLIAARGHSKPGQFDYACSNINCDYKRVYAWILQACYGIGGYDLTANSSSYDLSGNVVGGGAACLCAHGQTKAHTYYPGGYYP
EGIATLQFGVNEQYGCGSSG*

ATGAGTCAGCAATCTCCCTCGTCTATAAAGCCACCGACTTACCCGTGGACAGAATTCACTACCTACTCCTATATACTCCTGACCTGGGCTGTGTTAA
TCGCAGCTAGAGGCCATAGCAAACCCGGCCAGTTTGACTATGCCTGCTCTAATATCAACTGCGACTACAAACGAGTTTATGCTTGGATCCTCCAAGC
TTGTTACGGCATCGGCGGCTACGATCTTACAGCAAATTCTTCCTCCTACGACTTGTCCGGTAATGTCGTAGGTGGCGGAGCAGCTTGCCTATGTGCA
CATGGCCAGACGAAAGCGCACACTTATTACCCTGGTGGATATTATCCTGAAGGAATTGCTACATTGCAATTTGGTGTTAATGAGCAGTATGGCTGTG
GTAGCAGTGGCTAA

>Pseudocercospora_ulei_GCL012

MRFIALWLTLLTCAGLIAAKKRGKNKPGQIQYYCDQPSCSYGRALNWMYQACYGVGGSDVHPVTETRDQDHNINGAVGICQCAHGNTNEHTYFPGEP
KYPNGNATLTFGLSDKYGCQSKG*

ATGAGTCAGCCGCCTCCGTGGCCCGAGCAGGCGCAGCCAGAGGCTAAACCGCGGCACAGGATTCATCGCCCTCTGGCTCACACTCCTCACCTGCGCG
GGATTGATCGCCGCGAAGAAGCGCGGGAAAAACAAACCCGGCCAAATTCAGTACTACTGCGACCAACCCTCATGCAGCTACGGGCGAGCTCTGAATT
GGATGTACCAAGCTTGTTATGGCGTCGGCGGCAGTGACGTTCACCCAGTCACTGAGACCAGGGACCAGGACCACAACATCAACGGTGCCGTCGGGAT
TTGCCAATGTGCACATGGCAACACGAACGAACACACGTATTTCCCCGGGGAACCGAAATACCCCAACGGCAACGCGACTCTGACATTTGGTCTGTCG
GACAAGTATGGCTGTCAGAGCAAGGGTTGA

>Cercospora_cf._sigesbeckiae_PP_2012_071

MKFFLLVPLLASLPLAVARGKNKPGYFNYDCSDICDYEGVKRWIKQACEGIGGSSTTENGAWYNSNGGIIGGHAICLCARGQTSPHDYTISGDYPAG
TATLKFDASRGPYTCQRSG*

ATGAGTAAGCACTAGAGGAGCCATGGCATCGTAGTACTAATCTCAAGTAGAGTTCTTCCTACTCGTCCCACTCCTGGCCTCACTGCCTCTTGCCGTT
GCACGCGGCAAAAACAAACCTGGATGTAGGTCGACTGCAGTCTCACACCATACAATCGCTTCGCAGGGCTGACAATACATTCGCTGAAGACTTCAAT
TACGACTGCAGCGACATCTGCGACTACGAAGGCGTCAAACGATGGATCAAGCAAGCATGCGAGGGCATTGGAGGTTCGAGCACAACAGAAAACGGAG
CTTGGTATAACAGTAATGGCGGCATCATTGGAGGACACGCTATCTGCCTTTGCGCAAGAGGACAAACCAGCCCGCATGACTATACCATCAGTGGCGA
CTATCCTGCCGGCACAGCTACGCTCAAGTTTGATGCTTCCAGGGGTCCTTATACGTGTCAGCGCTCGGGGTAG

>Cercospora_cf._flagellaris_Arck_07

MKFFLLGLLLASLPLAIARGKNKPGYFDYNCNDICDYEGVKRWIKQACEGIGGSSTTENGAWYNSNGGIIGGHAICLCARGQTSPHDYAISGDYPPG
TATLKFDASRGPYTCQRSG*

ATGAGTAAGCACTAGAGGAGCCATGTCATCGTAGTACTAATCTCAAGTAGAATTCTTCCTACTCGGTCTACTCCTGGCCTCACTGCCTCTTGCCATT
GCACGCGGCAAAAACAAGCCAGGATGTAGGTGGACCGTCATCCTCCGTCAGACTGCACACCCAGACTCACCGGAAACGTCATAGACTTCGATTACAA
CTGCAACGACATTTGCGACTACGAAGGCGTCAAACGATGGATCAAGCAAGCATGCGAGGGCATTGGAGGTTCAAGCACAACAGAAAATGGAGCTTGG
TACAACAGTAATGGCGGCATCATTGGAGGACACGCTATCTGCCTTTGCGCGAGAGGACAAACCAGTCCGCATGACTATGCCATCAGTGGCGACTATC
CTCCCGGCACAGCTACGCTCAAGTTTGATGCTTCCAGGGGTCCCTACACTTGTCAACGCTCAGGGTAG

>Cercospora_brassicicola_Cer_68-18

MKFFLLVPLLASLPLAVARGKNKPGYFDYDCSDICDYEGVKRWIKQACEGIGGSSTTENGAWYNSNGGIIGGHAICLCARGQTSPHDYTISGDYPAG
TATLKFDASRGPYTCQRSG*

ATGAGTAAGCACTAGAGGAGCCATGGCATCGTAGTACTAATCTCAAGTAGAGTTCTTCCTACTCGTCCCACTCCTGGCCTCACTGCCTCTTGCCGTT

GCACGCGGCAAAAACAAACCTGGATGTAGGTCGACTGCAGTCTCACACCATACAATCGCTTCGCAGGGCTGACAATACATTCGCTGAAGACTTCGAT
TACGACTGCAGCGACATCTGCGACTACGAAGGCGTCAAACGATGGATCAAGCAAGCATGCGAGGGCATTGGAGGTTCGAGCACAACAGAAAACGGAG
CTTGGTATAACAGTAATGGCGGCATCATTGGAGGACACGCTATCTGCCTTTGCGCAAGAGGACAAACCAGCCCGCATGACTATACCATCAGTGGCGA
CTATCCTGCCGGCACAGCTACGCTCAAGTTTGATGCTTCCAGGGGTCCTTATACGTGTCAGCGCTCGGGGTAG
>Cercospora_kikuchii_ARG_18_001
MKFFLLVPLLASLPLAIARGKNKPGYFDYNCNDICDYEGVKRWIKQACEGIGGSSTTENGAWYNSNGGIIGGHAICLCARGQTSPHDYTISGDYPAG
TATLKFDASRGPYTCQRSG*

ATGAGTAAGCACTAGAGCAGCCATGTCATCGTAGTACTAATCTCAAGTAGAGTTCTTCCTACTCGTCCCACTCCTGGCCTCACTGCCTCTTGCCATT
GCACGCGGCAAAAACAAGCCAGGATGTAGGTGGACCGCAGTCTCACACCATACCATCGCTTCGCAGGGCTGACAATACATTCGCTGAAGACTTCGAT
TACAACTGCAACGACATCTGCGACTACGAAGGCGTTAAACGATGGATCAAGCAAGCATGCGAGGGTATTGGCGGATCGAGCACGACTGAAAATGGAG
CTTGGTATAACAGTAATGGCGGCATCATTGGAGGACACGCTATCTGCCTTTGCGCAAGAGGACAAACCAGCCCGCATGACTATACCATCAGTGGCGA
CTATCCTGCCGGCACAGCTACGCTCAAGTTTGATGCTTCCAGGGGTCCTTATACGTGTCAGCGCTCGGGGTAG
>Cercospora_kikuchii_A3
MKFFLLVPLLASLPLAIARGKNKPGYFDYNCNDICDYEGVKRWIKQACEGIGGSSTTENGAWYNSNGGIIGGHAICLCARGQTSPHDYTISGDYPAG
TATLKFDASRGPYTCQRSG*
ATGAGTAAGCACTAGAGCAGCCATGTCATCGTAGTACTAATCTCAAGTAGAGTTCTTCCTACTCGTCCCACTCCTGGCCTCACTGCCTCTTGCCATT
GCACGCGGCAAAAACAAGCCAGGATGTAGGTGGACCGCAGTCTCACACCATACCATCGCTTCGCAGGGCTGACAATACATTCGCTGAAGACTTCGAT
TACAACTGCAACGACATCTGCGACTACGAAGGCGTTAAACGATGGATCAAGCAAGCATGCGAGGGTATTGGCGGATCGAGCACGACTGAAAATGGAG
CTTGGTATAACAGTAATGGCGGCATCATTGGAGGACACGCTATCTGCCTTTGCGCAAGAGGACAAACCAGCCCGCATGACTATACCATCAGTGGCGA
CTATCCTGCCGGCACAGCTACGCTCAAGTTTGATGCTTCCAGGGGTCCTTATACGTGTCAGCGCTCGGGGTAG
>Cercospora_kikuchii_DNA
MKFFLLVPLLASLPLAIARGKNKPGYFDYNCNDICDYEGVKRWIKQACEGIGGSSTTENGAWYNSNGGIIGGHAICLCARGQTSPHDYTISGDYPAG
TATLKFDASRGPYTCQRSG*
ATGAGTAAGCACTAGAGCAGCCATGTCATCGTAGTACTAATCTCAAGTAGAGTTCTTCCTACTCGTCCCACTCCTGGCCTCACTGCCTCTTGCCATT
GCACGCGGCAAAAACAAGCCAGGATGTAGGTGGACCGCAGTCTCACACCATACCATCGCTTCGCAGGGCTGACAATACATTCGCTGAAGACTTCGAT
TACAACTGCAACGACATCTGCGACTACGAAGGCGTTAAACGATGGATCAAGCAAGCATGCGAGGGTATTGGCGGATCGAGCACGACTGAAAATGGAG
CTTGGTATAACAGTAATGGCGGCATCATTGGAGGACACGCTATCTGCCTTTGCGCAAGAGGACAAACCAGCCCGCATGACTATACCATCAGTGGCGA
CTATCCTGCCGGCACAGCTACGCTCAAGTTTGATGCTTCCAGGGGTCCTTATACGTGTCAGCGCTCGGGGTAG
>Cercospora_nicotianae_1110
MKFFLLVPLLASLPLAIARGKNKPGYFDYNCNDICDYEGVKRWIKQACEGIGGSSTTENGAWYNSNGGIIGGHAICLCARGQTSPHDYTISGDYPAG
TATLKFDASRGPYTCQRSG*
ATGAGTAAGCACTAGAGCAGCCATGTCATCGTAGTACTAATCTCAAGTAGAGTTCTTCCTACTCGTCCCACTCCTGGCCTCACTGCCTCTTGCCATT
GCACGCGGCAAAAACAAGCCAGGATGTAGGTGGACCGCAGTCTCACACCATACCATCGCTTCGCAGGGCTGACAATACATTCGCTGAAGACTTCGAT
TACAACTGCAACGACATCTGCGACTACGAAGGCGTTAAACGATGGATCAAGCAAGCATGCGAGGGTATTGGCGGATCGAGCACGACTGAAAATGGAG
CTTGGTATAACAGTAATGGCGGCATCATTGGAGGACACGCTATCTGCCTTTGCGCAAGAGGACAAACCAGCCCGCATGACTATACCATCAGTGGCGA
CTATCCTGCCGGCACAGCTACGCTCAAGTTTGATGCTTCCAGGGGTCCTTATACGTGTCAGCGCTCGGGGTAG
>Cercospora_sesami_Cers_52-10
MKLFLIAPLLTSLPLAIARGKNKPGYFDYNCSDICDYEGVKRWIKQACEGIGGSSTTENGAWYNQNGEIVGGHAICLCARGQTSPHDYTISGDYPPG
TATLKFNADKGPHTCQRSG*
ATGAGTAAGCACTAGAGCAGTCATGTCATCGTAGTACTGACCCAAGACAGAGCTCTTCTTGATCGCACCCCTCTTGACCTCACTGCCTCTTGCCATC
GCGAGAGGCAAAAACAAGCCAGGATGTAAGTTGACTGTCACTTGCGTTAGACTGCACTTTCAGACTTACCAGAAATGGTACAGATTTCGATTACAAC
TGCAGCGACATTTGTGACTACGAAGGCGTCAAACGATGGATCAAGCAAGCATGTGAGGGCATTGGGGGCTCGAGTACGACAGAAAATGGAGCTTGGT
ATAATCAGAATGGCGAAATTGTTGGAGGACATGCTATTTGTCTTTGTGCCCGCGGGCAAACCAGCCCGCACGACTATACTATCAGTGGCGACTACCC
TCCCGGCACAGCTACGCTGAAGTTTAACGCCGACAAGGGTCCTCATACGTGCCAGCGCTCGGGGTAG
>Cercospora_sesami_Cers_52-10
MKLFLIAPLLTSLPLAIARGKNKPGYFDYNCSDICDYEGVKRWIKQACEGIGGSSTTENGAWYNQNGEIVGGHAICLCARGQTSPHDYTISGDYPPG
TATLKFNADKGPHTCQRSG*

ATGAGTAAGCACTAGAGCAGTCATGTCATCGTAGTACTGACCCAAGACAGAGCTCTTCTTGATCGCACCCCTCTTGACCTCACTGCCTCTTGCCATC

GCGAGAGGCAAAAACAAGCCAGGATGTAAGTTGACTGTCACTTGCGTTAGACTGCACTTTCAGACTTACCAGAAATGGTACAGATTTCGATTACAAC

TGCAGCGACATTTGTGACTACGAAGGCGTCAAACGATGGATCAAGCAAGCATGTGAGGGCATTGGGGGCTCGAGTACGACAGAAAATGGAGCTTGGT

ATAATCAGAATGGCGAAATTGTTGGAGGACATGCTATTTGTCTTTGTGCCCGCGGGCAAACCAGCCCGCACGACTATACTATCAGTGGCGACTACCC

TCCCGGCACAGCTACGCTGAAGTTTAACGCCGACAAGGGTCCTCATACGTGCCAGCGCTCGGGGTAG

>Cercospora_beticola_ICMP_21690

MKFFLLVPLLASLPLAIARGKNKPGYFDYNCSDICDYEGVKRWIKQACEGIGGSSTTENGAWYNSNGGIIGGHAICLCARGQTSPHDYTISGDYPAG

TATLKFDASRGPYTCQRSG*

ATGAGTAAGCACTAGAGCAGCCGTGTCCTCATGGTACTAACCTCAAGTAGAGTTCTTCCTACTCGTTCCACTCCTGGCCTCACTGCCTCTTGCCATT

GCACGCGGCAAAAACAAGCCTGGATGTAGGTCGACCGCGATCTCACGTCATACAATCGCTTCGCGGAGCTAAACGATAAATTCGCTGAAGACTTCGA

TTACAACTGCAGCGACATCTGCGACTACGAAGGCGTCAAACGATGGATCAAGCAAGCATGCGAGGGCATTGGAGGTTCGAGCACAACAGAAAACGGA

GCTTGGTATAACAGTAATGGCGGCATCATTGGAGGACACGCTATCTGCCTTTGCGCGAGAGGACAAACCAGCCCGCATGACTATACCATCAGTGGCG

ACTATCCTGCCGGCACAGCTACGCTCAAGTTTGATGCTTCCAGGGGTCCCTATACGTGTCAGCGCTCGGGGTAG

>Cercospora_beticola_ICMP_21692

MKFFLLVPLLASLPLAIARGKNKPGYFDYNCSDICDYEGVKRWIKQACEGIGGSSTTENGAWYNSNGGIIGGHAICLCARGQTSPHDYTISGDYPAG

TATLKFDASRGPYTCQRSG*

ATGAGTAAGCACTAGAGCAGCCGTGTCCTCATGGTACTAACCTCAAGTAGAGTTCTTCCTACTCGTTCCACTCCTGGCCTCACTGCCTCTTGCCATT

GCACGCGGCAAAAACAAGCCTGGATGTAGGTCGACCGCGATCTCACGTCATACAATCGCTTCGCGGAGCTAAACGATAAATTCGCTGAAGACTTCGA

TTACAACTGCAGCGACATCTGCGACTACGAAGGCGTCAAACGATGGATCAAGCAAGCATGCGAGGGCATTGGAGGTTCGAGCACAACAGAAAACGGA

GCTTGGTATAACAGTAATGGCGGCATCATTGGAGGACACGCTATCTGCCTTTGCGCGAGAGGACAAACCAGCCCGCATGACTATACCATCAGTGGCG

ACTATCCTGCCGGCACAGCTACGCTCAAGTTTGATGCTTCCAGGGGTCCCTATACGTGTCAGCGCTCGGGGTAG

>Pseudocercospora_macadamiae_BRIP_55526

MKTTSIIPLLIFLASAKADNKPGRFTYTCQTAGCNQDRVWGWLYQACLGIGGTDLTNYVTIRDGGQSATGATTLCECARGSTRPHDYTISGNWPPGT

ASLKFGVGYKYSCQTAG*

ATGAGTAAGGCACTCTTTCAACCATATCAAGTATCATCATCAAGCTGATTTCACCCCAGAGACCACTTCCATAATCCCACTCCTAATCTTCCTCGCT

TCAGCCAAAGCTGACAACAAACCTGGCCGATTCACCTACACCTGCCAAACTGCCGGCTGTAATCAGGACCGAGTCTGGGGATGGCTTTATCAAGCCT

GTCTAGGTATCGGAGGCACTGATCTGACGAACTACGTGACGATAAGAGACGGAGGTCAATCTGCAACTGGAGCAACTACACTCTGTGAGTGCGCGCG

CGGCTCGACTAGACCACATGATTATACTATTTCGGGCAATTGGCCTCCTGGGACGGCATCTCTGAAGTTCGGCGTCGGATACAAATATTCGTGTCAG

ACTGCCGGATAG

>Cercospora_beticola_09-40

MKFFLLVPLLASLPLAIARSKNKPGYFDYNCSDICDYEGVKRWIKQACEGIGGSSTTENGAWYNSNGGIIEGHAICLCARGQTSPHDYTISGDYPAG

TATLKFDASRGPYTCQRSG*

ATGAGTAAGCACTAGAGCAGCCGTGTCCTCATGGTACTAACCTCAAGTAGAGTTCTTCCTACTCGTTCCACTCCTGGCCTCACTGCCTCTTGCCATT

GCACGCAGCAAAAACAAGCCTGGATGTAGGTCGACCGCGATCTCACGTCATACAATCGCTTCGCGGAGCTAAACGATAAATTCGCTGAAGACTTCGA

TTACAACTGCAGCGACATCTGCGACTACGAAGGCGTCAAACGATGGATCAAGCAAGCATGCGAGGGCATTGGAGGTTCGAGCACAACAGAAAACGGA

GCTTGGTATAACAGTAATGGCGGCATCATTGAAGGACACGCTATCTGCCTTTGCGCGAGAGGACAAACCAGCCCGCATGACTATACCATCAGTGGCG

ACTATCCTGCCGGCACAGCTACGCTCAAGTTTGATGCTTCCAGGGGTCCCTATACGTGTCAGCGCTCGGGGTAG

>Cercospora_citrullina_Cerl_37-09

MKFFLLVPLLAPLPLAIARGKNKPGYFDYNCKDICDYEGVKRWIKQACEGIGGSSTTENGAWYNANGGIIGGHAICLCARGQTSPHDYTISGDYPAG

TATLKFDASKGHYGCQRSG*

ATGAGTAAGCACTAGAGGAGCCACGTCATCGTAGTACTAATCTCAAGTAGAGTTCTTCCTACTCGTTCCACTCCTGGCCCCGCTGCCTCTTGCCATT

GCACGCGGCAAAAACAAGCCAGGGTGTAAGTCGACTGCGGTCTCACCACACACTTCCGCCTCGCAAGACTAACAATAAATCCGCCAAAGACTTCGAT

TACAACTGCAAGGACATCTGTGACTACGAGGGCGTCAAAAGATGGATCAAGCAAGCATGCGAGGGCATTGGCGGCTCCAGCACGACAGAGAATGGAG

CTTGGTACAACGCAAATGGCGGCATCATTGGAGGACACGCAATTTGCCTTTGCGCCAGGGGACAAACCAGCCCGCACGACTATACCATCAGTGGCGA

CTACCCTGCCGGCACAGCTACACTTAAGTTTGATGCTTCCAAAGGTCATTATGGGTGTCAACGCTCGGGGTAG

>Pseudocercospora_ulei_GCL012

MRFITLVPTLLTWAAVSTTVHGKGDNKPGKFTYSCFHLDCDANKAADWIKQACYGVGGHNVAEATTVYDVDRKVTGATGICQCARGQTNAADYYPDG

DYPAGKASLSFGLGKTYGCQSRG*

ATGAGTCAGCCATCTGCGTTGCCCGAGAAGGAGCAGCGGACGACTAACCCGCTGCACAGGATTCATCACCCTCGTCCCTACACTCCTCACCTGGGCG
GCGGTGTCGACCACCGTCCACGGCAAGGGCGACAACAAACCGGGCAAATTCACATACAGTTGCTTCCATCTCGACTGCGACGCCAACAAAGCCGCGG
ACTGGATCAAGCAAGCTTGCTACGGCGTCGGCGGCCATAACGTTGCAGAGGCCACGACCGTCTACGACGTGGATAGAAAAGTCACCGGCGCCACGGG
GATTTGCCAGTGTGCGCGTGGTCAGACGAACGCAGCCGATTATTACCCCGATGGAGATTACCCCGCCGGAAAGGCGTCGTTGAGTTTCGGTCTGGGC
AAGACCTATGGCTGCCAGAGCAGAGGTTGA
>Pseudocercospora_ulei_GCL012
MKSITLLATLLALTGLSSGKGKNKPGFFTYLCIKPGCDAKRVQAWILQACYGIGGYQVTDNSSTVDDYGRGISGKGTCICARGQTKEHDYTPDGDYP
KANVMLRFGGKGPLTCQSSGD*
ATGAGTGAGCGATCTCGGTTCCCCGAGAAGCCAACGACTAAACTGCCGCACAGAATCTATCACCCTCCTCGCCACGCTCCTCGCCTTGACCGGGTTG
AGTTCAGGCAAAGGCAAAAACAAACCCGGATTTTTTACCTATCTCTGCATAAAGCCCGGCTGCGACGCCAAGCGAGTCCAGGCGTGGATACTCCAAG
CTTGTTACGGCATCGGCGGCTATCAGGTGACCGATAATAGTTCGACCGTCGACGACTATGGCCGAGGCATATCCGGCAAGGGGACTTGCATATGCGC
ACGCGGTCAGACGAAAGAACACGATTATACCCCCGATGGAGACTATCCGAAAGCAAATGTGATGTTGAGATTTGGTGGGAAGGGGCCGTTGACCTGT
CAGAGCAGTGGTGATTGA
>Pseudocercospora_ulei_GCL012
MKFITSYPLLLTWAVLVAAKGNPKPGRFEYRCTDVCDYDRVMAWIFQACVGIGGYRLTSNAHITDILDRNVGGSAICICAHGQIEDGYYYPSGYYPE
GVVSLQFHKIGKFECGFSA*
ATGAGTTAGCAATCTCCGTCGTCTATAAAAGCCACACACTGACCCGCTGCACAGAATTCATCACCTCCTACCCACTACTCCTTACATGGGCGGTGTT
AGTCGCAGCCAAAGGCAATCCCAAACCCGGCAGGTTTGAGTATCGCTGCACCGATGTCTGCGACTACGACCGAGTTATGGCTTGGATCTTCCAAGCT
TGTGTCGGCATCGGCGGCTACCGTCTTACATCAAATGCTCATATCACCGACATTCTGGACAGGAACGTAGGTGGCAGTGCAATTTGCATATGTGCAC
ATGGCCAGATCGAAGATGGCTATTATTACCCGAGTGGATATTATCCTGAAGGAGTGGTATCGTTGCAATTTCACAAGATCGGGAAGTTCGAATGTGG
CTTCAGTGCCTGA
>Pseudocercospora_ulei_GCL012
MKFIALFPTLLTYAVLITAATHGKNKPGLLRYYCYDRNCDLGSVGQWIAQACRGLGGSPYLSLTPSYDKQGAAIGASGICECAHGDTKERVYYPGPP
DFPRGNATLTFYLKNKLSCQTT*
ATGAGTTGGTAGTCTTCGTTGCCCGTGAAGCAGCAGCTATAGACTAACCCGCTGTACAGAATTTATTGCCCTCTTCCCTACACTCCTTACCTACGCG
GTGTTGATTACCGCCGCGACTCACGGTAAAAATAAACCAGGCCTGCTTCGGTATTATTGCTATGACCGCAACTGCGACCTCGGTTCAGTTGGGCAGT
GGATAGCTCAAGCTTGTCGCGGCCTCGGCGGCAGTCCCTATCTTTCACTTACTCCTTCCTATGACAAGCAAGGCGCAGCCATAGGTGCATCTGGGAT
TTGCGAATGTGCGCATGGCGACACGAAAGAACGCGTTTATTACCCCGGACCACCTGATTTCCCGAGAGGAAACGCAACTCTGACATTTTATCTGAAG
AACAAGCTTAGCTGTCAGACCACATAA
>Pseudocercospora_ulei_GCL012
MKSITLFATLLALTGLSTGRGKNKPGFFRYTCEKSGCDADRAAAWVLQACHGIGGDHLVENDTVIDDYGRGLGAAGTCICARGQTKPHTYYPDGYYP
KGNVTLKFGGEGPYTCQSSGD*
ATGAGTGAGCAACCTCCGTTCCCCCGAGAAGCCATCGACTAAACTGCTGCACAGAATCCATTACCCTCTTCGCTACACTCCTCGCCTTGACTGGGTT
GAGTACAGGCAGAGGCAAAAACAAACCAGGCTTTTTTCGGTATACCTGCGAGAAGTCCGGCTGCGACGCAGACCGAGCCGCCGCCTGGGTCCTCCAA
GCTTGTCACGGCATCGGCGGCGACCACTTGGTCGAAAATGATACGGTTATCGACGACTATGGCAGAGGCTTGGGTGCCGCGGGGACTTGCATATGCG
CACGCGGTCAGACGAAACCGCACACTTATTACCCCGATGGATACTATCCTAAAGGAAATGTGACGTTGAAGTTTGGTGGGGAAGGGCCGTATACCTG
TCAGAGCAGTGGCGACTGA
>Pseudocercospora_ulei_GCL012
MRLLTHVSALLTWAVLITARGRGKNKPGTFYYRCIDIGCNYDRVEEWIGQACHGIGGKKINAMVPLSNSKGQVLGGTGVCACAHGQTKEHDYYPDGT
YYPAGNVTLRFDAKALYGCQSRGW*
ATGAGTCAGCCGCTCTCCGTTGCCCGAGAAGGAGACGCGATGGACTAAACCGCTGCACAGGACTCCTCACCCACGTCTCCGCCCTCCTCACCTGGGC
GGTGTTGATCACCGCGCGTGGCCGCGGCAAAAACAAGCCCGGCACATTCTACTACCGGTGCATCGACATCGGCTGCAACTACGACCGAGTCGAGGAG
TGGATCGGCCAAGCTTGTCACGGCATCGGCGGCAAAAAAATCAATGCCATGGTTCCTCTGAGCAACTCGAAAGGCCAAGTCCTCGGTGGCACGGGCG
TGTGCGCGTGTGCGCATGGTCAGACGAAAGAACACGACTATTACCCCGATGGAACATATTATCCGGCCGGGAACGTGACGCTGAGGTTTGATGCGAA
AGCGCTCTATGGCTGTCAGAGCCGTGGTTGGTAG
>Nothopassalora_personata_CPDP-13A
MIALTFFLAALPPLAYGRGKGDNKPGKFIYSCRDSGCNAEAVADWLGQACEGIGGSNLGETSSVYNATGAVVGGEMICLCAHGETDPHNYTISGGRY
PPGTVTLTFDVNAPYSCQSSG*

ATGAGTAGGTCTCTATACATTCGGAATCTACGCCGCTATCACTGACGTTCTCGTAGTTGCGCTTACCTTCTTCCTCGCCGCACTTCCTCCGCTAGCT

TATGGCAGGGGCAAAGGGGACAATAAACCAGGCAAATTCATATACAGTGTAAGTAGATAAATCTAGGCTTTGGAGGATTTACAAAATCCCAAGGTTG

TTTACAATTACATTATTGTCCTCTTAGTGTAGCCCTCTGTCTCGCCTCCTCTTATCTCCTAACCCCTTCCTAATTTCCTCTATCGCCTTACCCTAGA

ATCGAATTATACTAACGCCCTCGACCTAGTGCCGAGACTCGGGCTGCAACGCAGAAGCCGTCGCAGACTGGCTAGGGCAAGCCTGTGAAGGCATTGG

TGGCTCGAACCTAGGCGAGACCAGTTCAGTATATAACGCGACGGGTGCCGTTGTAGGCGGAGAGATGATTTGCTTGTGTGCGCATGGCGAGACGGAC

CCGCATAATTATACGATTAGTGGTGGACGTTATCCACCTGGCACGGTGACTTTGACGTTTGACGTTAATGCGCCTTATTCGTGTCAGAGTTCGGGTT

AG

>Cercospora_kikuchii_A3

MKLLLTTALLASLPLSYARGDNKPGYFDYTCNEICDYQKVKDWIQQACNGIGGAATTQDSPGVDEKGNNIGGTAVCLCAKGQTSPHDYTIEKGGDYP

AGKASLTFDATKGHFTCQRSG*

ATGAGTAAGCATCGTAGCTGCCATAGTGGCCTCTTGTCACTGACCCTAGACAGAGCTCCTCCTAACTACGGCCCTACTGGCCTCCCTACCTCTATCC

TACGCTCGCGGAGACAACAAGCCAGGATGTAAGAACACCACTCCTTGACAACAAATGACAAATCTGACAAAACAGCTCCCAGACTTCGACTATACCT

GCAACGAAATCTGTGATTATCAGAAAGTCAAAGACTGGATTCAACAAGCATGTAACGGCATCGGCGGTGCGGCCACTACTCAGGACTCACCTGGAGT

AGACGAGAAGGGTAACAATATCGGGGGAACAGCTGTCTGCCTTTGTGCGAAAGGTCAAACCAGTCCGCATGATTATACGATTGAGAAAGGTGGAGAT

TATCCTGCTGGTAAAGCTTCTCTTACGTTCGATGCTACGAAGGGGCACTTTACTTGTCAGCGTTCGGGGTGA

>Cercospora_cf._sigesbeckiae_PP_2012_071

MKLLLTTALLASLPLSYARGDNKPGYFDYTCNEICDYQKVKDWIQQACNGIGGAATTQDSPGVDEKGNNIGGTAVCLCAKGQTSPHDYTIEKGGDYP

AGKASLTFDATKGHFTCQRSG*

ATGAGTAAGCATCGTAGCTGCCATAGTGGCCTCTTGTCACTGACCCTAGACAGAGCTCCTCCTAACTACGGCCCTACTGGCCTCCCTACCTCTATCC

TACGCTCGCGGAGACAACAAGCCAGGATGTAAGAACACCACTCCTTGACAACAAATGACAAATCTGACAAAACAGCTCCCAGACTTCGACTATACCT

GCAACGAAATCTGTGATTATCAGAAAGTCAAAGACTGGATTCAACAAGCATGTAACGGCATCGGCGGTGCGGCCACTACTCAGGACTCACCTGGAGT

AGACGAGAAGGGTAACAATATCGGGGGAACAGCTGTCTGCCTTTGTGCGAAAGGTCAAACCAGTCCGCATGATTATACGATTGAGAAAGGTGGAGAT

TATCCTGCTGGTAAAGCTTCTCTTACGTTCGATGCTACGAAGGGGCACTTTACTTGTCAGCGTTCGGGGTGA

>Cercospora_kikuchii_DNA

MKLLLTTALLASLPLSYARGDNKPGYFDYTCNEICDYQKVKDWIQQACNGIGGAATTQDSPGVDEKGNNIGGTAVCLCAKGQTSPHDYTIEKGGDYP

AGKASLTFDATKGHFTCQRSG*

ATGAGTAAGCATCGTAGCTGCCATAGTGGCCTCTTGTCACTGACCCTAGACAGAGCTCCTCCTAACTACGGCCCTACTGGCCTCCCTACCTCTATCC

TACGCTCGCGGAGACAACAAGCCAGGATGTAAGAACACCACTCCTTGACAACAAATGACAAATCTGACAAAACAGCTCCCAGACTTCGACTATACCT

GCAACGAAATCTGTGATTATCAGAAAGTCAAAGACTGGATTCAACAAGCATGTAACGGCATCGGCGGTGCGGCCACTACTCAGGACTCACCTGGAGT

AGACGAGAAGGGTAACAATATCGGGGGAACAGCTGTCTGCCTTTGTGCGAAAGGTCAAACCAGTCCGCATGATTATACGATTGAGAAAGGTGGAGAT

TATCCTGCTGGTAAAGCTTCTCTTACGTTCGATGCTACGAAGGGGCACTTTACTTGTCAGCGTTCGGGGTGA

>Cercospora_sesami_Cers_52-10

MKLLLTTAILVSLPLVYARGDNKPGYFDYTCNEICDYQKVKDWIQQACNGIGGATTTENSPGVDEKGNNIGGTAVCLCAKGQTSPHDYTIEKGGQYP

AGKASLTFDATKGHYTCQRSG*

ATGAGTAAGCATCGAAGCTGCCATGATGCCCTCGTGTTACTGACCCTGTGCAGAACTTCTTCTGACTACAGCAATCTTGGTATCCCTACCTCTAGTC

TACGCTCGCGGAGACAACAAGCCAGGATGTGAGAACACCGCTCCTTGACAACATGTTACAAATCTGACAAAAGAGTCCTCAGACTTTGATTACACAT

GCAATGAAATCTGTGATTACCAAAAAGTCAAAGACTGGATTCAACAAGCATGTAACGGCATCGGTGGTGCAACCACTACCGAAAATTCACCTGGGGT

AGACGAGAAGGGTAACAATATTGGAGGAACAGCTGTCTGCCTTTGTGCGAAAGGTCAAACCAGTCCGCATGATTATACGATTGAGAAGGGTGGACAG

TATCCTGCTGGGAAAGCGTCTCTTACGTTTGATGCTACGAAGGGACACTATACTTGTCAGCGTTCGGGGTGA

>Fusarium_secorum_NRRL_62593

MQILSIIVFAMLGLASAKKHGKNKACEFTFDCASTNNCDRAKMEAWITQACEGIGGHSLTESGGRYDMHGGLVGYHAMCLCAHGNTKDHTYNIPRGV

YSAGTAQLNCEIDAPYSCQS*

ATGCGTATGTTATTTCGAATGTTCACAGTTCGTTATACAGCGTGACTACCTTTAAACTAACCACTCTTAGAAATCTTATCTATCATTGTTTTTGCGA

TGCTAGGCCTTGCCAGCGCAAAGAAACACGGAAAAAACAAGGCGTGCGAATTCACTTTTGACTGCGCTTCGACAAATAATTGCGATCGCGCAAAGAT

GGAAGCTTGGATTACGCAGGCTTGTGAAGGTATTGGTGGACATAGCCTCACAGAGTCGGGCGGCAGGTATGATATGCATGGTGGCCTTGTTGGGTAC

CATGCAATGTGTTTGTGCGCCCATGGTAACACCAAAGATCACACCTATAACATCCCACGCGGCGTATATTCTGCGGGAACGGCTCAATTGAATTGTG

AAATTGATGCTCCGTATAGCTGCCAGTCTTAG

>Fusarium secorum_CBS 175.32

MQILSIIVFAMLGLASAKKHGKNKACEFTFDCASTNNCDRAKMEAWITQACEGIGGHSLTESGGRYDMHGGLVGYHAMCLCAHGNTKDHTYNIPRGV
YSAGTAQLNCEIDAPYSCQS*
ATGCGTATGTTATTTCGAATGTTCACAGTTCGTTATACAGCGTGACTACCTTTAAACTAACCACTCTTAGAAATCTTATCTATCATTGTTTTTGCGA
TGCTAGGCCTTGCCAGCGCAAAGAAACACGGAAAAAACAAGGCGTGCGAATTCACTTTTGACTGCGCTTCGACAAATAATTGCGATCGCGCAAAGAT
GGAAGCTTGGATTACGCAGGCTTGTGAAGGTATTGGTGGACATAGCCTCACAGAGTCGGGCGGCAGGTATGATATGCATGGTGGCCTTGTTGGGTAC
CATGCAATGTGTTTGTGCGCCCATGGTAACACCAAAGATCACACCTATAACATCCCACGCGGCGTATATTCTGCGGGAACGGCTCAATTGAATTGTG
AAATTGATGCTCCGTATAGCTGCCAGTCTTAG
>Cercospora citrullina_Cerl
MKLLLTTALLASLPLSYARGDNKPGYFDYTCNEICDYQKVKAWIQQACNGIGGATTTQESPGVDEKGNNIGGTAVCLCARGQTSPHEYTIEKGGDYP
AGKASLTFDATKGHRTCQRSG*
ATGAGTAAGCATCGTAGCTGGCATAGTGTCCTCTTATTACTGACCCTAGACAGAGCTCCTCCTGACTACAGCCCTTCTGGCCTCCCTACCTCTATCC
TACGCTCGCGGAGACAACAAGCCAGGATGTAAGAACACCACTCCGCGGCATTATGTTAGGAATCTGACAAAACAGCTCCCAGACTTCGACTATACCT
GCAACGAAATCTGTGATTATCAGAAAGTCAAAGCCTGGATTCAACAAGCATGTAACGGCATCGGCGGTGCGACCACTACTCAGGAATCACCTGGAGT
AGACGAGAAGGGTAACAACATCGGAGGAACAGCTGTCTGCCTTTGTGCGAGAGGACAGACCAGCCCGCACGAGTACACGATCGAGAAAGGTGGGGAC
TATCCTGCTGGGAAGGCTTCTCTTACGTTTGATGCTACGAAAGGTCACCGTACTTGTCAGCGTTCGGGGTGA
>Sphaerulina_vaccinii_Sph-NS-03
MKLLIALLPFALASGPNKPGRFRYDCLYDGCDWNRVDAWLWQACRGIGGNELTKHESRYNNKKEFIGKTAICLCARGSIRPHTYRIEDDYPYRQAYL
DFKDMTIPYKCETFE*
ATGAGTACAATATATCCCGATCCATCGCTTTCATCACGCCAAGTCCCTAACTTTCTTCATTAGAACTCCTCATCGCCCTCTTACCCTTTGCACTTGC
AAGTGGCCCAAACAAGCCTGGCCGTAAGCATAACGCGCCATACATACATACATACCTTCTCACCATTCAGCGCTGATATAACATCACGATTAACAGG
CTTCCGATACGACTGCCTCTACGACGGCTGCGACTGGAACCGCGTCGATGCCTGGCTATGGCAAGCCTGTCGAGGCATTGGCGGAAACGAATTAACA
AAGCACGAATCCAGATACAATAACAAGAAAGAATTTATAGGCAAGACGGCGATCTGTCTATGCGCAAGAGGCTCGATACGTCCGCACACTTATCGCA
TTGAAGACGACTATCCTTATAGGCAGGCATATTTGGACTTCAAGGATATGACTATTCCGTATAAGTGTGAGACTTTCGAGTAG
>Sphaerulina_vaccinii_Sph-NS-01
MKLLIALLPFALASGPNKPGRFRYDCLYDGCDWNRVDAWLWQACRGIGGNELTKHESRYNNKKEFIGKTAICLCARGSIRPHTYRIEDDYPYRQAYL
DFKDMTIPYKCETFE*
ATGAGTACAATATATCCCGATCCATCGCTTTCATCACGCCAAGTCCCTAACTTTCTTCATTAGAACTCCTCATCGCCCTCTTACCCTTTGCACTTGC
AAGTGGCCCAAACAAGCCTGGCCGTAAGCATAACGCGCCATACATACATACATACCTTCTCACCATTCAGCGCTGATATAACATCACGATTAACAGG
CTTCCGATACGACTGCCTCTACGACGGCTGCGACTGGAACCGCGTCGATGCCTGGCTATGGCAAGCCTGTCGAGGCATTGGCGGAAACGAATTAACA
AAGCACGAATCCAGATACAATAACAAGAAAGAATTTATAGGCAAGACGGCGATCTGTCTATGCGCAAGAGGCTCGATACGTCCGCACACTTATCGCA
TTGAAGACGACTATCCTTATAGGCAGGCATATTTGGACTTCAAGGATATGACTATTCCGTATAAGTGTGAGACTTTCGAGTAG
>Cercospora_nicotianae_CBS_131.32
MRLLLAAALVASLPLSYARGDNKPGYFDYTCSEICDYQKVKDWILQACKGIGGASTTEDSPGVDEKGNNIGGTAVCLCARGQTSPHDYTIEKGGDYP
AGKASLTFDATKGHYTCQRSG*
ATGAGTAAGCATTGTAGGTGCCGTATCGTGCTCTCCTTACTGATGATAGTCAGGATTACTCCTAGCCGCAGCCCTTGTGGCCTCTCTACCTCTATCT
TACGCTCGGGGCGACAACAAACCAGGATGTAAGAAAACCACTCCTTGACAACATACTACCAATCTCTAAAACGATTCTCAGATTTCGACTACACCTG
CAGCGAAATCTGTGATTACCAAAAAGTCAAAGACTGGATTCTGCAAGCATGTAAAGGTATCGGCGGCGCATCCACTACAGAGGACTCACCTGGAGTG
GACGAGAAAGGCAATAACATCGGAGGAACAGCTGTCTGCCTTTGTGCGAGAGGACAGACCAGCCCGCATGATTATACGATCGAGAAAGGTGGAGACT
ATCCTGCTGGGAAGGCTTCTCTTACGTTTGATGCTACGAAGGGGCACTATACTTGTCAGCGTTCGGGGTGA
>Cercospora nicotianae_1110
MRLLLAAALVASLPLSYARGDNKPGYFDYTCSEICDYQKVKDWILQACKGIGGASTTEDSPGVDEKGNNIGGTAVCLCARGQTSPHDYTIEKGGDYP
AGKASLTFDATKGHYTCQRSG*
ATGAGTAAGCATTGTAGGTGCCGTATCGTGCTCTCCTTACTGATGATAGTCAGGATTACTCCTAGCCGCAGCCCTTGTGGCCTCTCTACCTCTATCT
TACGCTCGGGGCGACAACAAACCAGGATGTAAGAAAACCACTCCTTGACAACATACTACCAATCTCTAAAACGATTCTCAGATTTCGACTACACCTG
CAGCGAAATCTGTGATTACCAAAAAGTCAAAGACTGGATTCTGCAAGCATGTAAAGGTATCGGCGGCGCATCCACTACAGAGGACTCACCTGGAGTG
GACGAGAAAGGCAATAACATCGGAGGAACAGCTGTCTGCCTTTGTGCGAGAGGACAGACCAGCCCGCATGATTATACGATCGAGAAAGGTGGAGACT
ATCCTGCTGGGAAGGCTTCTCTTACGTTTGATGCTACGAAGGGGCACTATACTTGTCAGCGTTCGGGGTGA
>Fusarium_oxysporum_f._sp._spinaciae_Fus001

MQILSIIVFAMLGLASAKKHGKNKACEFTFDCASTNNCDRAKMEAWITQACEGIGGHSLTESGASYDMHGGLVGYHAMCLCAHGNTKDHTYNIPRGV
YSAGTAQLNCEIDAPYSCQS*
ATGCGTATGTTATTTCGAATGTTCACAGTTCGTTATACGGCGTGACTACCTTTCAACTAACCACTCTTAGAAATCTTATCTATCATTGTTTTTGCGA
TGCTAGGCCTTGCCAGCGCAAAGAAACACGGAAAAAACAAGGCGTGCGAATTCACTTTTGACTGCGCTTCGACAAATAATTGCGATCGCGCAAAGAT
GGAAGCTTGGATTACGCAGGCTTGTGAAGGTATTGGTGGACATAGCCTCACGGAGTCGGGCGCCAGCTATGATATGCATGGCGGCCTTGTTGGGTAC
CATGCAATGTGTTTGTGCGCCCATGGTAACACCAAAGATCACACCTATAACATCCCACGCGGCGTATATTCTGCGGGAACGGCTCAATTGAATTGTG
AAATTGATGCTCCGTATAGCTGCCAGTCTTAG
>Fusarium_oxysporum_f._sp._spinaciae_Fus173
MQILSIIVFAMLGLASAKKHGKNKACEFTFDCASTNNCDRAKMEAWITQACEGIGGHSLTESGASYDMHGGLVGYHAMCLCAHGNTKDHTYNIPRGV
YSAGTAQLNCEIDAPYSCQS*
ATGCGTATGTTATTTCGAATGTTCACAGTTCGTTATACGGCGTGACTACCTTTCAACTAACCACTCTTAGAAATCTTATCTATCATTGTTTTTGCGA
TGCTAGGCCTTGCCAGCGCAAAGAAACACGGAAAAAACAAGGCGTGCGAATTCACTTTTGACTGCGCTTCGACAAATAATTGCGATCGCGCAAAGAT
GGAAGCTTGGATTACGCAGGCTTGTGAAGGTATTGGTGGACATAGCCTCACGGAGTCGGGCGCCAGCTATGATATGCATGGCGGCCTTGTTGGGTAC
CATGCAATGTGTTTGTGCGCCCATGGTAACACCAAAGATCACACCTATAACATCCCACGCGGCGTATATTCTGCGGGAACGGCTCAATTGAATTGTG
AAATTGATGCTCCGTATAGCTGCCAGTCTTAG
>Fusarium_oxysporum_f._sp._spinaciae_MF15
MQILSIIVFAMLGLASAKKHGKNKACEFTFDCASTNNCDRAKMEAWITQACEGIGGHSLTESGASYDMHGGLVGYHAMCLCAHGNTKDHTYNIPRGV
YSAGTAQLNCEIDAPYSCQS*
ATGCGTATGTTATTTCGAATGTTCACAGTTCGTTATACGGCGTGACTACCTTTCAACTAACCACTCTTAGAAATCTTATCTATCATTGTTTTTGCGA
TGCTAGGCCTTGCCAGCGCAAAGAAACACGGAAAAAACAAGGCGTGCGAATTCACTTTTGACTGCGCTTCGACAAATAATTGCGATCGCGCAAAGAT
GGAAGCTTGGATTACGCAGGCTTGTGAAGGTATTGGTGGACATAGCCTCACGGAGTCGGGCGCCAGCTATGATATGCATGGCGGCCTTGTTGGGTAC
CATGCAATGTGTTTGTGCGCCCATGGTAACACCAAAGATCACACCTATAACATCCCACGCGGCGTATATTCTGCGGGAACGGCTCAATTGAATTGTG
AAATTGATGCTCCGTATAGCTGCCAGTCTTAG
>Fusarium_oxysporum_f._sp._spinaciae_MF42
MQILSIIVFAMLGLASAKKHGKNKACEFTFDCASTNNCDRAKMEAWITQACEGIGGHSLTESGASYDMHGGLVGYHAMCLCAHGNTKDHTYNIPRGV
YSAGTAQLNCEIDAPYSCQS*
ATGCGTATGTTATTTCGAATGTTCACAGTTCGTTATACGGCGTGACTACCTTTCAACTAACCACTCTTAGAAATCTTATCTATCATTGTTTTTGCGA
TGCTAGGCCTTGCCAGCGCAAAGAAACACGGAAAAAACAAGGCGTGCGAATTCACTTTTGACTGCGCTTCGACAAATAATTGCGATCGCGCAAAGAT
GGAAGCTTGGATTACGCAGGCTTGTGAAGGTATTGGTGGACATAGCCTCACGGAGTCGGGCGCCAGCTATGATATGCATGGCGGCCTTGTTGGGTAC
CATGCAATGTGTTTGTGCGCCCATGGTAACACCAAAGATCACACCTATAACATCCCACGCGGCGTATATTCTGCGGGAACGGCTCAATTGAATTGTG
AAATTGATGCTCCGTATAGCTGCCAGTCTTAG
>Fusarium_oxysporum_f._sp._spinaciae_Fus167
MQILSIIVFAMLGLASAKKHGKNKACEFTFDCASTNNCDRAKMEAWITQACEGIGGHSLTESGASYDMHGGLVGYHAMCLCAHGNTKDHTYNIPRGV
YSAGTAQLNCEIDAPYSCQS*
ATGCGTATGTTATTTCGAATGTTCACAGTTCGTTATACGGCGTGACTACCTTTCAACTAACCACTCTTAGAAATCTTATCTATCATTGTTTTTGCGA
TGCTAGGCCTTGCCAGCGCAAAGAAACACGGAAAAAACAAGGCGTGCGAATTCACTTTTGACTGCGCTTCGACAAATAATTGCGATCGCGCAAAGAT
GGAAGCTTGGATTACGCAGGCTTGTGAAGGTATTGGTGGACATAGCCTCACGGAGTCGGGCGCCAGCTATGATATGCATGGCGGCCTTGTTGGGTAC
CATGCAATGTGTTTGTGCGCCCATGGTAACACCAAAGATCACACCTATAACATCCCACGCGGCGTATATTCTGCGGGAACGGCTCAATTGAATTGTG
AAATTGATGCTCCGTATAGCTGCCAGTCTTAG
>Cercospora_beticola_ICMP_21692
MRLLLAAALVASLPLSYARGDNKPGYFDYTCSEICDYQKVKDWIQQACKGIGGASTTEDSPGVDEKGNNIGGTAVCLCARGQTSPHDYTIEKGGDYP
AGKASLTFDATKGHYTCQRSG*
ATGAGTAAGCATTGTAGGTGCCGTATCGTGCTCTCCTTACTGATGATAGTCAGGATTACTCCTAGCTGCAGCCCTTGTGGCCTCCCTACCTCTATCT
TACGCTCGGGGCGACAATAAACCAGGGTGTAAGGCAGCCATTCGCCGACGGAAACCTTGCAAAACTAACAAAAGAGCTCCCAGACTTCGATTACACC
TGCAGCGAAATCTGTGATTACCAAAAAGTCAAAGACTGGATCCAGCAAGCATGTAAAGGTATCGGCGGCGCATCAACTACAGAGGACTCACCTGGAG
TGGACGAGAAAGGCAATAACATCGGAGGAACAGCTGTCTGCCTTTGTGCGAGAGGACAGACCAGCCCGCATGATTATACGATCGAGAAAGGTGGAGA
CTATCCTGCTGGGAAGGCTTCTCTTACGTTTGATGCTACGAAGGGGCACTATACTTGTCAGCGGTCGGGCTGA
>Cercospora_apii_CBS_116455

MRLLLAAALVASLPLSYARGDNKPGYFDYTCSEICDYQKVKDWIQQACKGIGGASTTEDSPGVDEKGNNIGGTAVCLCARGQTSPHDYTIEKGGDYP
AGKASLTFDATKGHYTCQRSG*
ATGAGTAAGCATTGTAGGTGCCGTATCGTGCTCTCCTTACTGATGACAGTCAGGATTACTCCTAGCTGCAGCCCTTGTGGCCTCCCTACCTCTATCT
TACGCTCGGGGCGACAATAAACCAGGGTGTAAGGCAGCCATTCGCCGACGGAAACCTTGCAAAACTAACAAAAGAGCTCCCAGACTTCGATTACACC
TGCAGCGAAATCTGTGATTACCAAAAAGTCAAAGACTGGATCCAGCAAGCATGTAAAGGTATCGGCGGCGCATCAACTACAGAGGACTCACCTGGAG
TGGACGAGAAAGGCAATAACATCGGAGGAACAGCTGTCTGCCTTTGTGCGAGAGGACAGACCAGCCCGCATGATTATACGATCGAGAAAGGTGGAGA
CTATCCTGCTGGGAAGGCTTCTCTTACGTTTGATGCTACGAAGGGGCACTATACTTGTCAGCGGTCGGGCTGA
>Cercospora_beticola_ICMP_21690
MRLLLAAALVASLPLSYARGDNKPGYFDYTCSEICDYQKVKDWIQQACKGIGGASTTEDSPGVDEKGNNIGGTAVCLCARGQTSPHDYTIEKGGDYP
AGKASLTFDATKGHYTCQRSG*
ATGAGTAAGCATTGTAGGTGCCGTATCGTGCTCTCCTTACTGATGATAGTCAGGATTACTCCTAGCTGCAGCCCTTGTGGCCTCCCTACCTCTATCT
TACGCTCGGGGCGACAATAAACCAGGGTGTAAGGCAGCCATTCGCCGACGGAAACCTTGCAAAACTAACAAAAGAGCTCCCAGACTTCGATTACACC
TGCAGCGAAATCTGTGATTACCAAAAAGTCAAAGACTGGATCCAGCAAGCATGTAAAGGTATCGGCGGCGCATCAACTACAGAGGACTCACCTGGAG
TGGACGAGAAAGGCAATAACATCGGAGGAACAGCTGTCTGCCTTTGTGCGAGAGGACAGACCAGCCCGCATGATTATACGATCGAGAAAGGTGGAGA
CTATCCTGCTGGGAAGGCTTCTCTTACGTTTGATGCTACGAAGGGGCACTATACTTGTCAGCGGTCGGGCTGA
>Cercospora_apii_QCYBC
MRLLLAAALVASLPLSYARGDNKPGYFDYTCSEICDYQKVKDWIQQACKGIGGASTTEDSPGVDEKGNNIGGTAVCLCARGQTSPHDYTIEKGGDYP
AGKASLTFDATKGHYTCQRSG*
ATGAGTAAGCATTGTAGGTGCCGTATCGTGCTCTCCTTACTGATGACAGTCAGGATTACTCCTAGCTGCAGCCCTTGTGGCCTCCCTACCTCTATCT
TACGCTCGGGGCGACAATAAACCAGGGTGTAAGGCAGCCATTCGCCGACGGAAACCTTGCAAAACTAACAAAAGAGCTCCCAGACTTCGATTACACC
TGCAGCGAAATCTGTGATTACCAAAAAGTCAAAGACTGGATCCAGCAAGCATGTAAAGGTATCGGCGGCGCATCAACTACAGAGGACTCACCTGGAG
TGGACGAGAAAGGCAATAACATCGGAGGAACAGCTGTCTGCCTTTGTGCGAGAGGACAGACCAGCCCGCATGATTATACGATCGAGAAAGGTGGAGA
CTATCCTGCTGGGAAGGCTTCTCTTACGTTTGATGCTACGAAGGGGCACTATACTTGTCAGCGGTCGGGCTGA
>Cercospora_beticola_09-40
MRLLLAAALVASLPLSYARGDNKPGYFDYTCSEICDYQKVKDWIQQACKGIGGASTTEDSPGVDEKGNNIGGTAVCLCARGQTSPHDYTIEKGGDYP
AGKASLTFDATKGHYTCQRSG*
ATGAGTAAGCATTGTAGGTGCCGTATCGTGCTCTCCTTACTGATGATAGTCAGGATTACTCCTAGCTGCAGCCCTTGTGGCCTCCCTACCTCTATCT
TACGCTCGGGGCGACAATAAACCAGGGTGTAAGGCAGCCATTCGCCGACGGAAACCTTGCAAAACTAACAAAAGAGCTCCCAGACTTCGATTACACC
TGCAGCGAAATCTGTGATTACCAAAAAGTCAAAGACTGGATCCAGCAAGCATGTAAAGGTATCGGCGGCGCATCAACTACAGAGGACTCACCTGGAG
TGGACGAGAAAGGCAATAACATCGGAGGAACAGCTGTCTGCCTTTGTGCGAGAGGACAGACCAGCCCGCATGATTATACGATCGAGAAAGGTGGAGA
CTATCCTGCTGGGAAGGCTTCTCTTACGTTTGATGCTACGAAGGGGCACTATACTTGTCAGCGGTCGGGCTGA
>Cercospora_cf._flagellaris_Arck_07
MRLLLATALVASLPLSYARGDNKPGYFDYTCNEICDYQKVKDWIQQACKGIGGASTTEDSPGVDEKGNNIGGTAVCLCARGQTSPHDYTIEKGGDYP
AGKASLTFDATKEHYTCQRSG*
ATGAGTAAGCATTATAGGTGCCGTATCGTGCTCTCCTTACTGATGATAGTCAGGATTACTCCTAGCTACAGCCCTCGTGGCTTCCCTACCTCTATCT
TACGCTCGGGGCGACAACAAACCAGGATGTAAGGCAGCCATTCGCTGACGGAACCCTTGCAAAACTAACAAAAGAGCTCCCAGACTTCGATTACACC
TGCAACGAAATCTGTGATTACCAAAAAGTCAAAGACTGGATTCAGCAAGCATGTAAAGGTATCGGCGGCGCATCCACTACAGAGGACTCACCTGGAG
TGGACGAGAAAGGCAATAATATCGGAGGAACAGCTGTCTGCCTTTGTGCGAGAGGACAGACCAGCCCGCATGATTATACGATCGAGAAAGGTGGAGA
CTATCCTGCTGGGAAGGCTTCTCTTACGTTTGATGCTACGAAGGAGCACTATACTTGTCAGCGGTCGGGGTGA
>Fusarium begoniae_NRRL_25300
MQILPIITLAMLGHASAKKHGKNKACEFTFSCTSTNDCDTPKMEAWLTQACQGIGGSEIFQSGGIYTMHGEVTGHHAICLCAHGNTKDHTYDIASGA
LYSGGKATLNCDIAAPYTCQS*
ATGCGTAAGTTACTTCAAATGTTTATGATTAGCTATGCTAACTAGTTATAGAAATCTTGCCTATTATTACTCTTGCCATGCTGGGCCACGCCAGTGC
GAAGAAACATGGAAAGAACAAAGCGTGCGAATTCACTTTCAGTTGCACTTCTACAAACGATTGCGATACCCCAAAGATGGAAGCTTGGCTTACTCAG
GCTTGTCAAGGAATTGGGGGGTCTGAGATCTTTCAGTCTGGAGGTATATATACTATGCATGGTGAAGTCACTGGGCACCATGCAATTTGTTTGTGTG
CCCATGGTAATACCAAGGATCACACCTACGACATCGCATCTGGAGCATTGTATTCTGGAGGGAAGGCTACGCTTAATTGCGACATTGCTGCCCCATA
CACCTGCCAGTCTTAG
>Fusarium_sterilihyphosum_NRRL_25623

MQISSILVFATLGLASTKKHGKNKACEFIFDCTSTSDCDRAKMEAWTMQACQGIGGKRLLESGGIYDMHGGVVGHHAMCLCGHGNTKDHTYNIPPGV
YSEGTARLNCEIDAPYGCQS*
ATGCGTATGTTATTCTAAACGTTCACGGTTCGTCATGCAGTATGACTACCTTCAAGCTGACCATTCTTAGAAATTTCATCTATCCTTGTTTTTGCGA
CGTTAGGCCTTGCTAGCACGAAGAAACACGGCAAGAACAAGGCGTGCGAATTCATCTTTGATTGTACTTCAACAAGTGATTGCGATCGCGCGAAAAT
GGAGGCCTGGACTATGCAGGCTTGTCAAGGTATTGGCGGAAAAAGGCTCTTGGAGTCGGGCGGGATTTATGATATGCATGGTGGTGTTGTTGGGCAC
CACGCAATGTGTTTGTGCGGTCACGGAAACACCAAGGATCACACCTACAACATTCCACCTGGCGTATATTCCGAGGGAACAGCCCGGTTGAATTGCG
AGATTGATGCCCCATACGGCTGTCAGTCCTAG
>Fusarium tupiense_NRRL_53984
MQISSILVFATLGLASTKKHGKNKACEFIFDCTSTSDCDRAKMEAWTMQACQGIGGKRLLESGGIYDMHGGVVGHHAMCLCGHGNTKDHTYNIPPGV
YSEGTARLNCEIDAPYGCQS*
ATGCGTATGTTATTCTAAACGTTCACGGTTCGTCATGCAGTATGACTACCTTCAAGCTGACCATTCTTAGAAATTTCATCTATCCTTGTTTTTGCGA
CGTTAGGCCTTGCTAGCACGAAGAAACACGGCAAGAACAAGGCGTGCGAATTCATCTTTGATTGTACTTCAACAAGTGATTGCGATCGCGCGAAAAT
GGAGGCCTGGACTATGCAGGCTTGTCAAGGTATTGGCGGAAAAAGGCTCTTGGAGTCGGGCGGGATTTATGATATGCATGGTGGTGTTGTTGGGCAC
CACGCAATGTGTTTGTGCGGTCACGGAAACACCAAGGATCACACCTACAACATTCCACCTGGCGTATATTCCGAGGGAACAGCCCGGTTGAATTGCG
AGATTGATGCCCCATACGGCTGTCAGTCCTAG
>Fusarium_oxysporum_f._sp._spinaciae_MF15
MQILSIIVFAMLGLASAKKHGKNKACEFTFDCASTNDCDRAKMEAWITQACEGIGGHSLTESGGSYDIHGGGLVGYHAMCLCAHGNTKDHTYNIPGG
VYSAGTAQLNCEIDAPYSCQS*
ATGCGTATGTTATTTCGAATGTTCACAGTTCGTTATACAGCGTGACTACCTTTAAACTAACCACTCTTAGAAATCTTATCTATCATTGTTTTTGCGA
TGCTAGGCCTTGCCAGCGCAAAGAAACACGGAAAAAACAAGGCGTGCGAATTCACTTTTGACTGCGCTTCGACAAATGATTGCGATCGCGCAAAGAT
GGAAGCTTGGATTACGCAGGCTTGTGAAGGTATTGGTGGACATAGCCTCACAGAGTCGGGCGGCAGCTATGATATACATGGTGGTGGCCTTGTTGGG
TACCATGCAATGTGTTTGTGCGCCCATGGTAACACCAAAGATCACACCTATAACATCCCAGGCGGCGTATATTCTGCGGGAACGGCTCAATTGAATT
GTGAAATTGATGCTCCGTATAGCTGCCAGTCTTAG
>Fusarium_oxysporum_f._sp._spinaciae_MF34
MQILSIIVFAMLGLASAKKHGKNKACEFTFDCASTNDCDRAKMEAWITQACEGIGGHSLTESGGSYDIHGGGLVGYHAMCLCAHGNTKDHTYNIPGG
VYSAGTAQLNCEIDAPYSCQS*
ATGCGTATGTTATTTCGAATGTTCACAGTTCGTTATACAGCGTGACTACCTTTAAACTAACCACTCTTAGAAATCTTATCTATCATTGTTTTTGCGA
TGCTAGGCCTTGCCAGCGCAAAGAAACACGGAAAAAACAAGGCGTGCGAATTCACTTTTGACTGCGCTTCGACAAATGATTGCGATCGCGCAAAGAT
GGAAGCTTGGATTACGCAGGCTTGTGAAGGTATTGGTGGACATAGCCTCACAGAGTCGGGCGGCAGCTATGATATACATGGTGGTGGCCTTGTTGGG
TACCATGCAATGTGTTTGTGCGCCCATGGTAACACCAAAGATCACACCTATAACATCCCAGGCGGCGTATATTCTGCGGGAACGGCTCAATTGAATT
GTGAAATTGATGCTCCGTATAGCTGCCAGTCTTAG
>Fusarium_oxysporum_f._sp._spinaciae_Fus001
MQILSIIVFAMLGLASAKKHGKNKACEFTFDCASTNDCDRAKMEAWITQACEGIGGHSLTESGGSYDIHGGGLVGYHAMCLCAHGNTKDHTYNIPGG
VYSAGTAQLNCEIDAPYSCQS*
ATGCGTATGTTATTTCGAATGTTCACAGTTCGTTATACAGCGTGACTACCTTTAAACTAACCACTCTTAGAAATCTTATCTATCATTGTTTTTGCGA
TGCTAGGCCTTGCCAGCGCAAAGAAACACGGAAAAAACAAGGCGTGCGAATTCACTTTTGACTGCGCTTCGACAAATGATTGCGATCGCGCAAAGAT
GGAAGCTTGGATTACGCAGGCTTGTGAAGGTATTGGTGGACATAGCCTCACAGAGTCGGGCGGCAGCTATGATATACATGGTGGTGGCCTTGTTGGG
TACCATGCAATGTGTTTGTGCGCCCATGGTAACACCAAAGATCACACCTATAACATCCCAGGCGGCGTATATTCTGCGGGAACGGCTCAATTGAATT
GTGAAATTGATGCTCCGTATAGCTGCCAGTCTTAG
>Fusarium_oxysporum_f._sp._spinaciae_VPRI44294
MQILSIIVFAMLGLASAKKHGKNKACEFTFDCASTNDCDRAKMEAWITQACEGIGGHSLTESGGSYDIHGGGLVGYHAMCLCAHGNTKDHTYNIPGG
VYSAGTAQLNCEIDAPYSCQS*
ATGCGTATGTTATTTCGAATGTTCACAGTTCGTTATACAGCGTGACTACCTTTAAACTAACCACTCTTAGAAATCTTATCTATCATTGTTTTTGCGA
TGCTAGGCCTTGCCAGCGCAAAGAAACACGGAAAAAACAAGGCGTGCGAATTCACTTTTGACTGCGCTTCGACAAATGATTGCGATCGCGCAAAGAT
GGAAGCTTGGATTACGCAGGCTTGTGAAGGTATTGGTGGACATAGCCTCACAGAGTCGGGCGGCAGCTATGATATACATGGTGGTGGCCTTGTTGGG
TACCATGCAATGTGTTTGTGCGCCCATGGTAACACCAAAGATCACACCTATAACATCCCAGGCGGCGTATATTCTGCGGGAACGGCTCAATTGAATT
GTGAAATTGATGCTCCGTATAGCTGCCAGTCTTAG
>Fusarium_oxysporum_f._sp._spinaciae_Fus322

MQILSIIVFAMLGLASAKKHGKNKACEFTFDCASTNDCDRAKMEAWITQACEGIGGHSLTESGGSYDIHGGGLVGYHAMCLCAHGNTKDHTYNIPGG
VYSAGTAQLNCEIDAPYSCQS*
ATGCGTATGTTATTTCGAATGTTCACAGTTCGTTATACAGCGTGACTACCTTTAAACTAACCACTCTTAGAAATCTTATCTATCATTGTTTTTGCGA
TGCTAGGCCTTGCCAGCGCAAAGAAACACGGAAAAAACAAGGCGTGCGAATTCACTTTTGACTGCGCTTCGACAAATGATTGCGATCGCGCAAAGAT
GGAAGCTTGGATTACGCAGGCTTGTGAAGGTATTGGTGGACATAGCCTCACAGAGTCGGGCGGCAGCTATGATATACATGGTGGTGGCCTTGTTGGG
TACCATGCAATGTGTTTGTGCGCCCATGGTAACACCAAAGATCACACCTATAACATCCCAGGCGGCGTATATTCTGCGGGAACGGCTCAATTGAATT
GTGAAATTGATGCTCCGTATAGCTGCCAGTCTTAG
>Fusarium_oxysporum_f._sp._spinaciae_Fus059
MQILSIIVFAMLGLASAKKHGKNKACEFTFDCASTNDCDRAKMEAWITQACEGIGGHSLTESGGSYDIHGGGLVGYHAMCLCAHGNTKDHTYNIPGG
VYSAGTAQLNCEIDAPYSCQS*
ATGCGTATGTTATTTCGAATGTTCACAGTTCGTTATACAGCGTGACTACCTTTAAACTAACCACTCTTAGAAATCTTATCTATCATTGTTTTTGCGA
TGCTAGGCCTTGCCAGCGCAAAGAAACACGGAAAAAACAAGGCGTGCGAATTCACTTTTGACTGCGCTTCGACAAATGATTGCGATCGCGCAAAGAT
GGAAGCTTGGATTACGCAGGCTTGTGAAGGTATTGGTGGACATAGCCTCACAGAGTCGGGCGGCAGCTATGATATACATGGTGGTGGCCTTGTTGGG
TACCATGCAATGTGTTTGTGCGCCCATGGTAACACCAAAGATCACACCTATAACATCCCAGGCGGCGTATATTCTGCGGGAACGGCTCAATTGAATT
GTGAAATTGATGCTCCGTATAGCTGCCAGTCTTAG
>Fusarium_oxysporum_f._sp._spinaciae_Fus173
MQILSIIVFAMLGLASAKKHGKNKACEFTFDCASTNDCDRAKMEAWITQACEGIGGHSLTESGGSYDIHGGGLVGYHAMCLCAHGNTKDHTYNIPGG
VYSAGTAQLNCEIDAPYSCQS*
ATGCGTATGTTATTTCGAATGTTCACAGTTCGTTATACAGCGTGACTACCTTTAAACTAACCACTCTTAGAAATCTTATCTATCATTGTTTTTGCGA
TGCTAGGCCTTGCCAGCGCAAAGAAACACGGAAAAAACAAGGCGTGCGAATTCACTTTTGACTGCGCTTCGACAAATGATTGCGATCGCGCAAAGAT
GGAAGCTTGGATTACGCAGGCTTGTGAAGGTATTGGTGGACATAGCCTCACAGAGTCGGGCGGCAGCTATGATATACATGGTGGTGGCCTTGTTGGG
TACCATGCAATGTGTTTGTGCGCCCATGGTAACACCAAAGATCACACCTATAACATCCCAGGCGGCGTATATTCTGCGGGAACGGCTCAATTGAATT
GTGAAATTGATGCTCCGTATAGCTGCCAGTCTTAG
>Fusarium_oxysporum_f._sp._spinaciae_Fus057
MQILSIIVFAMLGLASAKKHGKNKACEFTFDCASTNDCDRAKMEAWITQACEGIGGHSLTESGGSYDIHGGGLVGYHAMCLCAHGNTKDHTYNIPGG
VYSAGTAQLNCEIDAPYSCQS*
ATGCGTATGTTATTTCGAATGTTCACAGTTCGTTATACAGCGTGACTACCTTTAAACTAACCACTCTTAGAAATCTTATCTATCATTGTTTTTGCGA
TGCTAGGCCTTGCCAGCGCAAAGAAACACGGAAAAAACAAGGCGTGCGAATTCACTTTTGACTGCGCTTCGACAAATGATTGCGATCGCGCAAAGAT
GGAAGCTTGGATTACGCAGGCTTGTGAAGGTATTGGTGGACATAGCCTCACAGAGTCGGGCGGCAGCTATGATATACATGGTGGTGGCCTTGTTGGG
TACCATGCAATGTGTTTGTGCGCCCATGGTAACACCAAAGATCACACCTATAACATCCCAGGCGGCGTATATTCTGCGGGAACGGCTCAATTGAATT
GTGAAATTGATGCTCCGTATAGCTGCCAGTCTTAG
>Fusarium_oxysporum_f._sp._spinaciae_Fus254
MQILSIIVFAMLGLASAKKHGKNKACEFTFDCASTNDCDRAKMEAWITQACEGIGGHSLTESGGSYDIHGGGLVGYHAMCLCAHGNTKDHTYNIPGG
VYSAGTAQLNCEIDAPYSCQS*
ATGCGTATGTTATTTCGAATGTTCACAGTTCGTTATACAGCGTGACTACCTTTAAACTAACCACTCTTAGAAATCTTATCTATCATTGTTTTTGCGA
TGCTAGGCCTTGCCAGCGCAAAGAAACACGGAAAAAACAAGGCGTGCGAATTCACTTTTGACTGCGCTTCGACAAATGATTGCGATCGCGCAAAGAT
GGAAGCTTGGATTACGCAGGCTTGTGAAGGTATTGGTGGACATAGCCTCACAGAGTCGGGCGGCAGCTATGATATACATGGTGGTGGCCTTGTTGGG
TACCATGCAATGTGTTTGTGCGCCCATGGTAACACCAAAGATCACACCTATAACATCCCAGGCGGCGTATATTCTGCGGGAACGGCTCAATTGAATT
GTGAAATTGATGCTCCGTATAGCTGCCAGTCTTAG
>Fusarium_sterilihyphosum_NRRL_25623
MQIASVLVFATLGLASTKEHGKNKACEFIFDCTSTNDCDRAKMEAWIMQACQGIGGRRLLESGGIYDMHGGVVGHHAMCLCGHGNTKDHTYNIPHGV
CSEGTARLNCEIDAPYSCQS*
ATGCGTATGTTATTCTAAACGTTCACAGTTCGTTATGCAGTATGACTACCTTCAAGCTGACCATTCTTAGAAATTGCATCTGTCCTTGTTTTTGCGA
CGTTAGGCCTTGCTAGCACGAAGGAACACGGCAAGAACAAGGCGTGCGAATTCATCTTTGATTGTACTTCAACAAATGATTGCGATCGCGCGAAAAT
GGAGGCTTGGATTATGCAGGCTTGTCAAGGTATTGGTGGAAGAAGGCTCTTGGAGTCGGGCGGGATTTATGATATGCATGGTGGTGTTGTTGGGCAC
CACGCGATGTGTTTGTGCGGTCACGGAAACACCAAGGATCACACCTACAACATTCCACATGGCGTATGTTCCGAGGGAACAGCCCGGTTGAATTGCG
AGATTGATGCCCCATACAGCTGTCAGTCCTAG
>Fusarium_tupiense_NRRL_53984

MQIASVLVFATLGLASTKEHGKNKACEFIFDCTSTNDCDRAKMEAWIMQACQGIGGRRLLESGGIYDMHGGVVGHHAMCLCGHGNTKDHTYNIPHGV
CSEGTARLNCEIDAPYSCQS*
ATGCGTATGTTATTCTAAACGTTCACAGTTCGTTATGCAGTATGACTACCTTCAAGCTGACCATTCTTAGAAATTGCATCTGTCCTTGTTTTTGCGA
CGTTAGGCCTTGCTAGCACGAAGGAACACGGCAAGAACAAGGCGTGCGAATTCATCTTTGATTGTACTTCAACAAATGATTGCGATCGCGCGAAAAT
GGAGGCTTGGATTATGCAGGCTTGTCAAGGTATTGGTGGAAGAAGGCTCTTGGAGTCGGGCGGGATTTATGATATGCATGGTGGTGTTGTTGGGCAC
CACGCGATGTGTTTGTGCGGTCACGGAAACACCAAGGATCACACCTACAACATTCCACATGGCGTATGTTCCGAGGGAACAGCCCGGTTGAATTGCG
AGATTGATGCCCCATACAGCTGTCAGTCCTAG
>Fusarium_denticulatum_NRRL_25311
MQILPIVVLAMLGQASAKRHGKNKACEFTFECTTTNDCDTAKMEAWIFQACQGIGGDRTVQSGGVYTGHGDLTGHQATCLCAHGNTKDHTYDIASGA
LYSGGKAKLNCDVTAPYNCQS*
ATGCGTAAGTTACTTCGAATGCTTATAAATAGCTATGCTAACTAGTTATAGAAATCTTGCCTATTGTTGTTCTTGCCATGCTGGGCCAAGCCAGTGC
GAAGAGACATGGAAAGAACAAGGCGTGCGAATTCACTTTCGAATGCACTACTACGAACGATTGCGATACCGCAAAGATGGAAGCTTGGATTTTTCAG
GCTTGTCAAGGAATTGGAGGGGATAGGACCGTTCAGTCTGGAGGTGTATACACCGGTCATGGTGACCTCACTGGGCACCAAGCAACTTGTTTATGTG
CCCATGGTAATACCAAGGATCACACCTACGACATCGCATCTGGAGCATTGTATTCTGGAGGGAAGGCTAAGCTGAATTGCGACGTTACTGCCCCATA
CAACTGCCAGTCTTAG
>Fusarium_tupiense_NRRL_53984
MQILPIITLAMLGHASAKKHGKNKACEFTFSCTSTNDCDAPKMEAWLTQACQGIGGYEITESGGIYTTYGDFTGYHGMCLCAHGNTKDHTYVITPGV
LYSGGTATLNCNVAAPYTCQS*
ATGCGTAAGTTACTTCAAGTGTTTATGATTAGCTATGCTAACTGGTTATAGAAATCTTGCCTATTATTACTCTTGCCATGCTGGGCCACGCCAGTGC
GAAGAAACATGGAAAGAACAAAGCATGCGAATTCACTTTCAGTTGCACTTCTACAAACGATTGCGATGCCCCAAAGATGGAAGCGTGGCTTACTCAG
GCTTGTCAAGGAATTGGGGGGTATGAGATCACTGAGTCTGGAGGTATATATACTACGTATGGTGACTTCACTGGGTACCATGGAATGTGTTTGTGTG
CCCATGGTAATACCAAAGACCACACCTACGTCATCACACCTGGAGTATTGTATTCTGGAGGGACGGCTACGTTGAATTGCAACGTTGCTGCCCCATA
CACCTGCCAGTCTTAG
>Pseudocercospora_ulei_GCL012
MKFTTYSSVLLTWAVFIAAKGHNKAGRFVYECTHMNCDYKRVYEWILQACHGIGGRDLQGNSSSVDLKGNILGGGAICLCAHGQTAEHTYSPGGYYP
EGTVWLEFGNVPRWNCHDGV*
ATGAGTTAGCCATCTCCCTCGTCTATCAAGCTACAGTCTGACCCGCTGCACAAAATTCACCACCTACTCCTCTGTACTCCTGACCTGGGCCGTGTTC
ATCGCAGCCAAAGGCCATAACAAAGCCGGCAGGTTTGTCTATGAATGCACCCATATGAACTGCGACTACAAACGAGTTTATGAATGGATCCTCCAAG
CATGTCACGGCATCGGCGGCCGCGATCTTCAAGGAAATTCCAGCTCCGTCGACTTGAAGGGCAATATCTTGGGGGGCGGAGCGATTTGCCTGTGTGC
ACATGGCCAGACCGCAGAGCATACTTATTCCCCGGGTGGATATTATCCTGAAGGAACGGTCTGGTTGGAATTTGGCAACGTCCCGCGGTGGAACTGT
CACGACGGTGTCTAA
>Fusarium_fracticaudum_CBS_137234
MQILPIITLAMLGHASAKKHGKNKACEFTFSCTSTNDCDTPKMEAWITQACQGIGGYEITESGGIYTRYGDFTGYHGICLCAHGNTKDHTYVITPGV
LYSGGTATLNCNVAAPYSCQS*
ATGCGTCAGTTACTTCAAATGTTTATGAGTAGCTATGCTAACTAGTTATAGAAATCTTGCCTATTATTACTCTTGCCATGCTGGGCCACGCCAGTGC
TAAGAAACATGGAAAGAATAAAGCGTGCGAATTCACTTTCAGTTGCACTTCTACAAACGATTGCGATACCCCAAAGATGGAAGCTTGGATTACTCAG
GCTTGTCAAGGAATTGGGGGGTATGAGATCACTGAGTCTGGAGGTATATATACTAGGTATGGTGACTTCACTGGGTACCATGGAATTTGTTTGTGTG
CCCATGGTAATACCAAAGATCACACCTACGTCATCACACCTGGTGTATTGTATTCTGGAGGGACGGCTACGTTGAATTGCAACGTTGCTGCCCCATA
CAGCTGCCAGTCTTAG
>Fusarium_sp._NRRL_47473
MQILPIITLAMLGHASAKKHGKNKACEFTFSCTSTNDCDAPKMEAWITQACQGIGGSKILESGGIYTMYGDFTGYHGMCLCAHGNTKDHNYVITPGV
LYSGGTATLNCDVAAPYSCQS*
ATGCGTAAGTTACTTCAAATGTTTATGATTAGCTATGCTAACTAGTTATAGAAATCTTGCCTATTATTACTCTTGCCATGCTGGGCCACGCCAGTGC
GAAGAAACATGGAAAGAACAAAGCGTGCGAATTCACTTTCAGTTGCACTTCTACAAACGATTGCGATGCCCCAAAGATGGAAGCTTGGATTACTCAG
GCTTGTCAAGGAATTGGGGGGTCTAAGATCCTTGAGTCTGGAGGTATATATACTATGTATGGTGACTTCACTGGGTACCATGGAATGTGTTTGTGTG
CCCATGGTAATACCAAAGATCACAACTACGTCATCACACCTGGAGTATTGTATTCTGGAGGGACGGCTACGTTGAATTGCGACGTTGCTGCCCCATA
CAGCTGCCAGTCCTAG
>Fusarium_mexicanum_NRRL_53147

MQILPIITLAMLGHASAKKHGKNKACEFTFSCTSTNDCDAPKMEAWITQACQGIGGSKILESGGIYTMYGDFTGYHGMCLCAHGNTKDHNYVITPGV

LYSGGTATLNCDVAAPYSCQS*

ATGCGTAAGTTACTTCAAATGTTTATGATTAGCTATGCTAACTAGTTATAGAAATCTTGCCTATTATTACTCTTGCCATGCTGGGCCACGCCAGTGC

GAAGAAACATGGAAAGAACAAAGCGTGCGAATTCACTTTCAGTTGCACTTCTACAAACGATTGCGATGCCCCAAAGATGGAAGCTTGGATTACTCAG

GCTTGTCAAGGAATTGGGGGGTCTAAGATCCTTGAGTCTGGAGGTATATATACTATGTATGGTGACTTCACTGGGTACCATGGAATGTGTTTGTGTG

CCCATGGTAATACCAAAGATCACAACTACGTCATCACACCTGGAGTATTGTATTCTGGAGGGACGGCTACGTTGAATTGCGACGTTGCTGCCCCATA

CAGCTGCCAGTCCTAG

>Fusarium_pilosicola_CMWF1183

MQILPIIILAMLGHASAKRHGKNKACEFTFNCTSTNDCDSRKMEAWLTQACQGIGGHKIVESGGIYTSHGDFTGYHGICLCAHGNTKDHTYVITPGV

LYSGGTATLNCNVDAPYTCQS*

ATGCGTAAGTTACTTTAAATGTTTATAATTAGCTATGCTAACTAGTTATAGAAATCTTGCCAATTATTATTCTTGCCATGCTGGGCCATGCCAGTGC

GAAGAGACATGGAAAGAATAAAGCGTGCGAATTCACTTTCAATTGCACTTCTACAAACGATTGCGATAGCCGAAAGATGGAAGCTTGGCTTACTCAG

GCTTGTCAAGGTATTGGGGGTCATAAGATCGTTGAATCTGGAGGTATATATACAAGCCATGGCGACTTCACTGGGTACCATGGAATTTGTTTGTGTG

CCCATGGTAATACCAAAGATCACACCTACGTCATCACACCTGGAGTATTGTATTCTGGAGGGACGGCTACGTTGAATTGCAACGTTGATGCCCCATA

CACCTGCCAGTCTTAG

>Fusarium_sterilihyphosum_NRRL_25623

MQILPIITLAMLGHASAKKHGKNKACEFTFSCTSTSDCDAPKMEAWITQACQGIGGSKILESGGIYTRYGDFTGYHGMCLCAHGNTKDHTYVITPGV

LYSGGTATLNCNVAAPYSCQS*

ATGCGTCAGTTACTTCAAATGTTTATGAGTAGCTATGCTAACTAGTTATAGAAATCTTGCCTATTATTACTCTTGCCATGCTGGGCCATGCCAGTGC

GAAGAAACATGGAAAGAACAAAGCGTGCGAATTCACTTTCAGTTGCACTTCTACAAGCGATTGCGATGCCCCAAAGATGGAAGCTTGGATTACTCAG

GCTTGTCAAGGAATTGGGGGGTCTAAGATCCTTGAGTCTGGAGGTATATATACTAGGTATGGTGACTTCACTGGGTACCATGGAATGTGTTTGTGTG

CCCATGGTAATACCAAAGATCACACCTACGTCATCACACCTGGAGTATTGTATTCTGGAGGGACGGCTACGTTGAATTGCAACGTTGCTGCCCCATA

CAGCTGCCAGTCTTAG

>Fusarium_sp._NRRL_53293

MQLLPIITLAMLGHASAKKHGKNKACEFTFSCTSTNDCNAPKMEAWITQACQGIGGSKILESGGIYTRYGDFTGYHGMCLCAHGNTKDHTYVITPGV

LYSGGTATLNCNVAAPYSCQS*

ATGCGTAAGTTACTTCAAATGTTTATGATTAGCTATGCTAACTGGTTATAGAACTCTTGCCTATTATTACTCTTGCCATGCTGGGCCACGCCAGTGC

GAAGAAACATGGAAAGAACAAAGCATGCGAATTCACTTTCAGTTGCACTTCTACAAACGATTGCAATGCCCCAAAGATGGAAGCTTGGATTACTCAG

GCTTGTCAAGGAATCGGGGGGTCTAAGATCCTTGAGTCTGGAGGTATATATACTAGGTATGGTGACTTCACTGGGTACCATGGAATGTGTTTGTGTG

CCCATGGTAATACCAAAGATCACACCTACGTCATCACACCTGGAGTATTGTATTCTGGAGGGACGGCTACGTTGAATTGCAACGTTGCTGCCCCATA

CAGCTGCCAGTCTTAG

>Pseudocercospora_ulei_GCL012

MNFIALFSTPLTYAVLITAKHYGDNKPGQFWYYCEQIDCNMGAVGQWLNQACYGIGGPQSSKIYPMALTNGSVTGAELICECAHGSTKEHTYYPDPH

KYPRGFVNLKFHLSDKLSCQST*

ATGAGTCAGCAGTCTCCATTGCCTAAGAAGTAGCAACTATTGACTAAACCGCTGTACAGATTTCATTGCCCTGTTCTCTACACCCCTTACCTACGCA

GTGTTGATCACCGCCAAACACTACGGTGACAACAAACCAGGCCAGTTTTGGTATTATTGCGAACAAATCGACTGCAACATGGGGGCAGTCGGGCAGT

GGCTAAATCAAGCTTGTTACGGCATCGGCGGCCCTCAGTCGAGTAAAATTTATCCTATGGCTCTTACCAACGGCTCAGTCACAGGTGCCGAACTGAT

TTGCGAATGTGCACATGGCAGCACGAAAGAACACACTTATTACCCCGATCCACACAAATACCCGAGAGGATTCGTAAACCTGAAATTTCATCTGTCT

GACAAGCTTAGCTGTCAAAGCACATAG

>Pseudocercospora_ulei_GCL012

MRLITLVSTLLAWAVTADGTRHGDNKPGLFVYTCDQPGCNFSRVLDWIVQACHGVGGHGLDGNKTNTNEQNQVIGGGGVCACAHGNTKEHDYYPDEP

YWPKGVATLRFSLRLRHGCATIH*

ATGAGTCAGCAGTTTCCGTTGCCCGAGAAGAAGAAGCTGTAGACTAAACCGCTGCACAGGATTGATCACCCTAGTCTCTACACTCCTTGCCTGGGCT

GTTACAGCCGACGGCACCCGCCACGGCGACAACAAACCGGGCTTATTTGTTTATACCTGTGACCAGCCCGGTTGCAACTTCAGCCGCGTTTTAGACT

GGATTGTCCAAGCTTGCCACGGCGTCGGCGGGCACGGCCTTGATGGAAATAAAACAAACACCAACGAGCAAAATCAAGTTATAGGTGGCGGAGGGGT

TTGCGCATGTGCGCATGGTAACACGAAAGAACACGATTATTACCCTGATGAACCTTATTGGCCGAAAGGAGTTGCGACGTTAAGGTTTAGTCTGCGT

CTGAGGCATGGCTGTGCGACCATACATTGA

>Pseudocercospora_ulei_GCL012

MRVTALLFSLAVLATAKWWPEIPHHKPNKPGGFEYLCAHDDCDYEKVALWIRVTCGHIGGHRVVDNVTLWGRDGQVAGGSGVCLCVDGDIQPYLYHP
TKWYPPGKIRLSFGAEATYGCESNGKTKEKGH*
ATGAGTAAGCCACCTCCGGTGGCCCCACCCCCGCCGAAGAGGAAGTGCCGAAACTAAAACCCGCGTGCGCGCACAGGAGTCACGGCCTTGCTTTTCT
CCCTGGCGGTGTTGGCCACGGCCAAATGGTGGCCCGAGATCCCCCACCACAAACCCAACAAACCGGGGGGATTCGAATACCTCTGCGCCCACGACGA
CTGCGACTACGAGAAAGTGGCGTTGTGGATCCGGGTGACGTGCGGCCACATCGGCGGCCACCGCGTCGTGGACAACGTGACCCTCTGGGGCCGCGAC
GGCCAGGTCGCGGGCGGCTCCGGCGTGTGCCTGTGTGTGGACGGTGACATCCAACCATACCTCTATCACCCGACCAAATGGTACCCTCCGGGAAAGA
TCAGGTTGAGCTTCGGCGCGGAAGCGACCTATGGCTGCGAGAGCAATGGGAAGACGAAGGAGAAGGGGCATTAG
>Fusarium_newnesense_NRRL_66241
MQISSIFILTILGFASAKRHGKNKACEFTLNCTSTGDCDSDKMQSWILQACQGIGGGSLTGAGGIYSASDGALSGYHAICLCAHGNTGDYTYHIPSG
QYSAATATLNCNIDAPYSCQS*
ATGCGTATGTTGACAGTCCGTTATGTAATACAGCTATATTCCAACTGACTACTCTTAGAAATCTCTTCTATCTTTATTCTTACAATACTTGGCTTTG
CTAGCGCAAAGAGACACGGAAAGAACAAGGCTTGCGAGTTTACGCTCAATTGCACTTCGACAGGAGACTGCGATTCCGACAAGATGCAAAGCTGGAT
CCTTCAGGCTTGTCAAGGTATTGGTGGGGGTTCACTCACAGGGGCGGGCGGTATATACAGCGCGAGTGATGGTGCATTGTCTGGGTATCATGCGATA
TGTTTGTGTGCTCACGGCAACACTGGAGATTACACGTATCACATCCCATCCGGCCAATATTCTGCAGCAACGGCCACGTTGAACTGCAACATCGATG
CTCCGTACAGCTGTCAGTCTTAA
>Fusarium_sp._NRRL_25184
MQISSIFILTILGFASAKRHGKNKACEFTLNCTSTGDCDSDKMQSWILQACQGIGGGSLTGAGGIYRASDGASSGYHAICLCAHGDTNDWTYHIPSG
QYSAATATLNCNIDAPYSCQS*
ATGCGTATGTTAACAGTCCATTATGTAATACAGCTGTATTCCAACTGACTACTCTTAGAAATCTCTTCTATCTTTATTCTTACAATACTTGGCTTTG
CTAGCGCAAAGAGACATGGAAAAAACAAGGCTTGCGAGTTCACGCTCAATTGCACTTCGACAGGAGACTGCGATTCCGACAAGATGCAAAGCTGGAT
CCTTCAGGCTTGTCAAGGTATTGGCGGGGGTTCACTCACAGGGGCGGGCGGTATATACAGAGCGAGTGACGGTGCATCGTCTGGGTATCATGCGATA
TGTTTGTGTGCCCACGGCGACACCAACGATTGGACGTATCACATCCCATCCGGTCAATATTCTGCAGCAACGGCCACGTTGAACTGCAACATCGATG
CTCCGTACAGCTGTCAGTCTTAA
>Passalora_sequoiae_9LC2
MKLTTISLLALLGLAEAKRTGNKPGLWTVTCDSPADPVKCNEDKIIDYISLECKNNGGSGLPNINTQFNANGKVVGVQAVCACVHGDKKQWDYTVTE
SGYPAGKVSLRFGLKAKYGCQKTG*
ATGAGTTAGTATTCCTGCAAATCTCTCGAGGGTCGATTTAGACTGAAAACACCTAGAACTCACTACCATATCTCTCCTGGCCCTGCTCGGCCTCGCC
GAAGCCAAAAGGACAGGAAATAAGCCCGGCCTTTGGACCGTAACGTGCGATTCTCCGGCTGACCCAGTCAAATGCAACGAAGATAAGATCATAGATT
ACATTTCGCTCGAATGCAAGAACAACGGCGGATCGGGTTTACCAAACATAAATACCCAATTCAATGCGAATGGCAAGGTGGTTGGAGTACAAGCCGT
CTGCGCATGTGTTCATGGGGACAAGAAACAGTGGGATTACACGGTGACAGAGTCAGGATATCCAGCGGGCAAAGTTTCCCTGAGGTTCGGGCTGAAA
GCCAAGTATGGCTGTCAGAAGACGGGATAG
>Passalora_sequoiae_9LC2
MRLRIVFLSLLGVAAASPWFGVGGKQKGNKPGKFGFECNVADASCDLETIETWMLNRCESIGGSRLVKVDTWREHGFLKKVEGICACAHGSTKARDY
GQTDEPGIPKAHYWLEFGLDQTYGCQRTGAQ*
ATGAGTCAGTGCTTCTACACAACTCTCGATGAGCTTGTTTACTAAATCCTCGACTCTGTCTAGGGCTCAGAATCGTCTTCCTAAGCCTACTTGGCGT
CGCCGCAGCGTCCCCTTGGTTTGGGGTTGGTGGGAAGCAGAAAGGCAACAAGCCGGGCAAGTTTGGCTTCGAATGCAACGTTGCGGACGCCAGTTGC
GACCTCGAAACTATTGAGACATGGATGTTAAATCGATGTGAATCGATCGGCGGATCGAGGTTGGTCAAAGTCGATACTTGGCGGGAGCACGGCTTCC
TGAAGAAGGTTGAGGGCATCTGCGCGTGTGCACACGGAAGCACGAAAGCGAGAGACTACGGGCAAACCGATGAGCCGGGCATTCCCAAGGCTCATTA
CTGGCTCGAGTTCGGGCTTGATCAAACGTACGGCTGTCAGAGAACTGGAGCTCAGTAG
>Passalora_sequoiae_9LC2
MKLSIVLTALSASLVTAKRGANKPAYIAFDCKDHCKPEDMSNITAWVINKCWDIGGFHWTREEVHYFGKQIVAYSGICLCAHGNTREQYSNPHDERT
DMYLPEGVGSVTFGLDTTFGVCHKNG*
ATGAGTTCGTCGTTCCATCTGCCACCTCTCGGATACATTCTGACACCCTGCGCTGACCTATGCATAGAACTCTCCATCGTCCTCACAGCCCTTTCGG
CCTCCCTAGTCACCGCTAAGCGCGGCGCGAACAAGCCCGCCTACATCGCTTTCGACTGCAAGGATCATTGCAAGCCTGAAGATATGTCGAATATTAC
TGCTTGGGTCATCAACAAATGTTGGGACATTGGCGGATTCCACTGGACCAGGGAAGAGGTTCATTACTTTGGGAAACAAATTGTGGCTTATTCTGGG
ATTTGTTTGTGTGCACATGGGAATACGAGGGAGCAGTACTCGAATCCGCATGACGAGAGGACGGATATGTATCTCCCGGAGGGGGTTGGGAGTGTGA
CGTTTGGATTGGACACGACGTTTGGGGTCTGTCATAAGAATGGGTAG
>Pseudocercospora_ulei_GCL012

MRSIALLPTLLAYAGLITAAKHGKNKPGKFQYYCDHAGCNAGRVVYWLNQACKGIGGAYMDVTDITMSLDNLVTGAFGICKCVHGDIKEHVYFPTEP
DWPAGNVTLKFYLKENYGCATR*
ATGAGTCAGCCATCTCCGTTGCCCGAGAAGGAGTAGCTCTAGACTAAACCGCCGTACAGGATCCATCGCCCTCCTCCCCACACTCCTTGCCTACGCG
GGATTGATCACCGCTGCGAAACACGGGAAAAACAAACCCGGCAAATTTCAGTACTATTGCGACCATGCCGGGTGCAATGCCGGGCGCGTTGTATACT
GGCTGAACCAAGCCTGTAAAGGCATCGGCGGCGCTTACATGGACGTAACCGATATCACTATGAGCCTCGACAACCTAGTGACAGGTGCCTTCGGGAT
TTGCAAATGCGTCCACGGCGACATAAAAGAACACGTTTATTTCCCCACCGAGCCGGATTGGCCGGCAGGCAACGTGACTCTCAAATTCTATCTGAAA
GAAAATTATGGTTGTGCGACCAGATGA
>Passalora_sequoiae_9LC2
MKIMTATSLLAVTGFVTARNYGNKPGQWNVYCDLWPSALCREDILFDWVVGECKANGGWDLVDHSAIYRRTPRGKDVVSGFHGICTCVHGDKKQWTY
GMEGKGLPDAIAYLTFGLNDTWHCQRTG*
ATGAGTTAGTACTCCCTTACCATTAAAGACTTGACCTGTACTGAGCTTCGCACAGAAATCATGACTGCCACATCCCTCCTGGCTGTGACTGGCTTCG
TCACGGCGAGAAACTATGGAAATAAGCCTGGGCAATGGAACGTCTACTGCGATCTATGGCCATCCGCGCTCTGTAGGGAAGACATACTCTTTGATTG
GGTTGTGGGGGAATGTAAGGCCAACGGCGGGTGGGACCTTGTAGACCACAGTGCGATTTACCGCCGCACGCCCAGAGGAAAAGATGTTGTGAGTGGA
TTCCATGGCATCTGTACATGTGTGCATGGCGACAAGAAGCAGTGGACCTACGGGATGGAGGGGAAAGGGTTACCAGATGCCATCGCATACTTGACAT
TTGGACTGAATGACACGTGGCATTGCCAAAGGACCGGTTAG
>Fusarium_oxysporum_f._sp._gladioli_G76
MQISSIFILTMLGFASAKRHGKSKACEFMLNCTSTGDCDSDKMQSWILQACQGIGGGSLIGAGGIYRASDGALSGYHAICLCAHGDTNDWTYHIPSG
QYSAATATLNCNIDAPYSCQS*
ATGCGTATGTTGACAGTCCATTATGTAATACAGCAATATTCCAACTGACTACTCTTAGAAATATCTTCCATCTTTATTCTTACAATGCTTGGCTTTG
CTAGCGCAAAGAGACATGGAAAAAGCAAGGCTTGCGAGTTCATGCTCAATTGCACTTCGACAGGAGACTGCGATTCCGACAAGATGCAAAGCTGGAT
CCTTCAGGCTTGTCAAGGTATTGGCGGGGGTTCACTCATAGGGGCGGGCGGTATATACAGAGCGAGTGACGGTGCATTGTCTGGGTATCATGCGATA
TGTTTGTGTGCCCACGGCGACACCAACGATTGGACGTATCACATCCCATCCGGCCAATATTCTGCAGCAACGGCCACGTTGAACTGCAACATCGATG
CTCCGTACAGCTGTCAGTCTTAA
>Fusarium_oxysporum_f._sp._gladioli_G14
MQISSIFILTMLGFASAKRHGKSKACEFMLNCTSTGDCDSDKMQSWILQACQGIGGGSLIGAGGIYRASDGALSGYHAICLCAHGDTNDWTYHIPSG
QYSAATATLNCNIDAPYSCQS*
ATGCGTATGTTGACAGTCCATTATGTAATACAGCAATATTCCAACTGACTACTCTTAGAAATATCTTCCATCTTTATTCTTACAATGCTTGGCTTTG
CTAGCGCAAAGAGACATGGAAAAAGCAAGGCTTGCGAGTTCATGCTCAATTGCACTTCGACAGGAGACTGCGATTCCGACAAGATGCAAAGCTGGAT
CCTTCAGGCTTGTCAAGGTATTGGCGGGGGTTCACTCATAGGGGCGGGCGGTATATACAGAGCGAGTGACGGTGCATTGTCTGGGTATCATGCGATA
TGTTTGTGTGCCCACGGCGACACCAACGATTGGACGTATCACATCCCATCCGGCCAATATTCTGCAGCAACGGCCACGTTGAACTGCAACATCGATG
CTCCGTACAGCTGTCAGTCTTAA
>Passalora_sequoiae_9LC2
MKVVVLHLLAVVGIVTAQQQGNKPGLFQYLCEDDKNCPEQVMKNWIDAKCKSSWAGGRATIKNSAITDYFTKKVIGYAAICTCCHGGLSNTDWNGLK
TPPGNASLGFGLNNKYGCQSTGAP*
ATGAGTGAGTCTTCATAACTCGCCATCAAGGTCGTACACTGAGATTTGCTTAGAAGTCGTTGTCCTCCACCTCCTAGCTGTGGTCGGTATCGTCACA
GCTCAGCAACAAGGGAACAAGCCCGGGTTATTCCAGTACCTATGCGAGGACGATAAAAACTGCCCCGAACAGGTTATGAAGAACTGGATCGATGCTA
AATGCAAATCGAGTTGGGCCGGCGGCCGGGCGACCATCAAGAACAGTGCTATCACCGATTATTTCACCAAGAAAGTGATTGGATATGCAGCCATCTG
CACATGTTGCCATGGCGGCCTAAGCAACACGGACTGGAATGGACTCAAGACTCCACCAGGCAATGCTTCGCTGGGTTTCGGGCTTAATAACAAGTAT
GGGTGTCAATCAACGGGAGCCCCATGA
>Passalora_sequoiae_9LC2
MNFSIFSLLAFKLLALAVADHGHNKPGKFTYTCTARASDCNTDRMRSWMEQECKMSYIGGSQLTHVERDHNPLLVYTILSGICECAHGDKKQQNYSF
APMGYFPPGYATLTFRLDQKYSCSRI*
ATGAGTTAGTATTTCCATATCTTCGTCGAAATCTCGTCCTAACCAGCTCTGCCCAGATTTTAGTATCTTCTCCTTGCTAGCCTTCAAGCTGCTCGCC
CTCGCCGTAGCTGATCATGGCCACAATAAGCCAGGCAAATTCACATACACATGCACTGCTCGAGCGTCAGACTGCAACACCGATCGCATGCGAAGCT
GGATGGAGCAAGAATGCAAAATGAGCTACATCGGCGGTAGTCAATTGACTCATGTCGAACGGGACCACAACCCGCTACTAGTGTACACCATCTTGTC
TGGAATATGCGAGTGTGCGCACGGGGACAAAAAGCAGCAGAATTACAGTTTCGCGCCAATGGGTTACTTTCCTCCTGGCTATGCGACGCTGACGTTC
AGGCTTGACCAGAAGTATTCTTGTTCGAGGATCTAA
>Passalora_sequoiae_9LC2

MRPTIAFLALLALAAAKHGHNKAGKFVYKCDHVEWCQWWRIQDWMQQECEAIGGGVTDYKVEYKNTEGLTTGGEAVCLCAHGDTEQHDYSIVDEVHW
FPRGTVTLRFGLGTTFGVCHKNG*
ATGAGTTCGCACTTGCCCATACCTCCTGAGAATCTATAACACTAAGACTTTGTCAGGACCCACCATCGCCTTCTTAGCCCTGCTCGCTCTCGCCGCA
GCTAAACATGGTCATAACAAGGCAGGCAAGTTCGTATACAAATGCGATCATGTCGAATGGTGCCAATGGTGGCGCATACAAGACTGGATGCAGCAGG
AATGCGAAGCTATTGGCGGAGGAGTGACGGACTACAAGGTTGAGTACAAGAACACAGAAGGACTGACGACTGGAGGGGAGGCCGTCTGCCTATGTGC
ACACGGAGATACAGAGCAGCATGATTACAGCATCGTCGACGAAGTGCATTGGTTCCCGCGAGGTACAGTTACCCTAAGGTTCGGCCTTGGTACGACG
TTCGGCGTTTGCCATAAGAACGGATGA
>Teratosphaeria_pseudoeucalypti_CMW51515
MRLSILPIILGLFFLSATAKHGKNKPCNIDFQCDLQLQLAKKCDETKMKDWACKGIGAASVAAAGTTFDLHTNPVGYHAVCTCAHGNTKDHDYVISG
DTGQWTKGTATLRCSAPFVKGRWTCHNTA*
ATGAGTACGCACACCTGTGCTCAGCTGCTGTTCATTCATCTAACATCTCATTGACAGGGCTCTCAATCCTTCCGATCATCCTCGGACTCTTCTTCCT
GTCGGCGACGGCCAAGCATGGCAAGAACAAGCCATGCAACATAGACTTCCAGTGCGATCTTCAACTACAACTGGCGAAAAAATGCGACGAGACGAAG
ATGAAAGACTGGGTTGCTCAGGTTAGCATGCCTGACAGTCGCAAAGCTTCCTTCATGAATCACGAAGGACCAACTGATCAGTCCGCTTTCTAGGCTT
GCAAGGGCATCGGCGCCGCTTCCGTGGCAGCCGCCGGTACTACGTTCGATCTGCACACCAATCCAGTAGGCTATCACGCGGTTTGTACGTGCGCACA
CGGCAACACGAAGGACCACGACTACGTCATTAGCGGTGACACAGGGCAATGGACAAAGGGTACCGCGACGTTGAGGTGCAGTGCCCCGTTTGTTAAA
GGGCGTTGGACCTGTCACAATACCGCATGA
>Teratosphaeria_pseudoeucalypti_CMW49161
MRLSILPIILGLFFLSATAKHGKNKPCNIDFQCDLQLQLAKKCDETKMKDWACKGIGAASVAAAGTTFDLHTNPVGYHAVCTCAHGNTKDHDYVISG
DTGQWTKGTATLRCSAPFVKGRWTCHNTA*
ATGAGTACGCACACCTGTGCTCAGCTGCTGTTCATTCATCTAACATCTCATTGACAGGGCTCTCAATCCTTCCGATCATCCTCGGACTCTTCTTCCT
GTCGGCGACGGCCAAGCATGGCAAGAACAAGCCATGCAACATAGACTTCCAGTGCGATCTTCAACTACAACTGGCGAAAAAATGCGACGAGACGAAG
ATGAAAGACTGGGTTGCTCAGGTTAGCATGCCTGACAGTCGCAAAGCTTCCTTCATGAATCACGAAGGACCAACTGATCAGTCCGCTTTCTAGGCTT
GCAAGGGCATCGGCGCCGCTTCCGTGGCAGCCGCTGGTACTACGTTCGATCTGCACACCAATCCAGTAGGCTATCACGCGGTTTGTACGTGCGCACA
CGGCAACACGAAGGACCACGACTACGTCATTAGCGGTGACACAGGGCAATGGACAAAGGGTACCGCGACGTTGAGGTGCAGTGCCCCGTTTGTTAAA
GGGCGTTGGACCTGTCACAATACCGCATGA
>Teratosphaeria_pseudoeucalypti_CMW49159
MRLSILPIILGLFFLSATAKHGKNKPCNIDFQCDLQLQLAKKCDETKMKDWACKGIGAASVAAAGTTFDLHTNPVGYHAVCTCAHGNTKDHDYVISG
DTGQWTKGTATLRCSAPFVKGRWTCHNTA*
ATGAGTACGCACACCTGTGCTCAGCTGCTGTTCATTCATCTAACATCTCATTGACAGGGCTCTCAATCCTTCCGATCATCCTCGGACTCTTCTTCCT
GTCGGCGACGGCCAAGCATGGCAAGAACAAGCCATGCAACATAGACTTCCAGTGCGATCTTCAACTACAACTGGCGAAAAAATGCGACGAGACGAAG
ATGAAAGACTGGGTTGCTCAGGTTAGCATGCCTGACAGTCGCAAAGCTTCCTTCATGAATCACGAAGGACCAACTGATCAGTCCGCTTTCTAGGCTT
GCAAGGGCATCGGCGCCGCTTCCGTGGCAGCCGCTGGTACTACGTTCGATCTGCACACCAATCCAGTAGGCTATCACGCGGTTTGTACGTGCGCACA
CGGCAACACGAAGGACCACGACTACGTCATTAGCGGTGACACAGGGCAATGGACAAAGGGTACCGCGACGTTGAGGTGCAGTGCCCCGTTTGTTAAA
GGGCGTTGGACCTGTCACAATACCGCATGA
>Teratosphaeria_zuluensis_CMW17320
MKLSTIPVILGLLSIATEAKHGKNKPCKIEFNCDAAVQQSAAGCNSDQMEAWACTGIGAHDLAENGFMSDPNHRYIGYSAICICAHGNTKNHDYVID
QGWEKGTATLRCGHPDTSGWYTCASEG*
ATGAGTACCCACAGCCACCACTCAAGATGCTTGTCGTCCTTCTGACATCAAGCCATAGAGCTCTCGACTATTCCAGTCATCCTCGGCCTCTTGTCAA
TCGCGACAGAGGCCAAGCACGGCAAGAACAAACCATGCAAGATTGAATTCAACTGTGATGCCGCGGTGCAACAATCAGCAGCAGGCTGCAACTCCGA
TCAAATGGAAGCCTGGGTTAAACAAGTCAGTCAATTCTCACTGCGATATTCGGGCCTCACTCCTTGACTGACTTTGCACTGCATTACCTCAGGCTTG
CACCGGCATAGGTGCTCACGATCTTGCGGAGAACGGTTTCATGAGTGATCCTAATCATAGGTATATCGGCTACTCAGCGATCTGCATATGCGCGCAT
GGGAACACCAAGAATCATGACTATGTCATCGACCAAGGGTGGGAGAAAGGCACGGCAACATTACGATGCGGTCATCCTGACACCAGCGGGTGGTACA
CTTGTGCCTCGGAGGGATGA
>Teratosphaeria_eucalypti_CMW54005
MRLSILPIILGLFFLSATAKHGKNKPCNIDFQCDPQLQLAKKCDETKMTDWACKGIGAASVASAGTTFDLHTNPVACHAVCTCAHGNTKDHDYVISG
DTGQWTKGTAKLRCSAPFVKGNWGCHDTA*
ATGAGTACGCACACCTGTGCTCAGCCGCTGTTCATTTATCTAACATCTCATTGACAGGGCTCTCAATCCTTCCGATCATCCTCGGACTCTTCTTCCT
GTCGGCGACGGCCAAGCATGGCAAGAACAAGCCATGCAACATAGACTTCCAGTGCGATCCTCAACTACAACTGGCGAAAAAATGCGACGAGACGAAG

ATGACAGACTGGGTTGCTCAGGTAAGCATGCCTGACAGTCGCAAAGCTTCCTTCATGAATCACGAAGGACCAACTGATCAGTCCGCTTTCTAGGCTT
GCAAGGGCATCGGCGCCGCTTCCGTGGCGTCCGCCGGTACTACGTTCGATCTGCACACCAATCCAGTAGCCTGTCACGCGGTGTGTACGTGCGCACA
CGGCAACACGAAGGACCACGACTACGTCATTAGCGGTGACACAGGGCAATGGACAAAGGGTACCGCGAAGTTGAGGTGCAGTGCCCCGTTTGTTAAA
GGGAATTGGGGCTGTCACGATACCGCATGA
>Teratosphaeria_eucalypti_CMW55930
MRLSILPIILGLFFLSATAKHGKNKPCNIDFQCDPQLQLAKKCDETKMTDWACKGIGAASVASAGTTFDLHTNPVACHAVCTCAHGNTKDHDYVISG
DTGQWTKGTAKLRCSAPFVKGNWGCHDTA*
ATGAGTACGCACACCTGTGCTCAGCCGCTGTTCATTTATCTAACATCTCATTGACAGGGCTCTCAATCCTTCCGATCATCCTCGGACTCTTCTTCCT
GTCGGCGACGGCCAAGCATGGCAAGAACAAGCCATGCAACATAGACTTCCAGTGCGATCCTCAACTACAACTGGCGAAAAAATGCGACGAGACGAAG
ATGACAGACTGGGTTGCTCAGGTAAGCATGCCTGACAGTCGCAAAGCTTCCTTCATGAATCACGAAGGACCAACTGATCAGTCCGCTTTCTAGGCTT
GCAAGGGCATCGGCGCCGCTTCCGTGGCGTCCGCCGGTACTACGTTCGATCTGCACACCAATCCAGTAGCCTGTCACGCGGTGTGTACGTGCGCACA
CGGCAACACGAAGGACCACGACTACGTCATTAGCGGTGACACAGGGCAATGGACAAAGGGTACCGCGAAGTTGAGGTGCAGTGCCCCGTTTGTTAAA
GGGAATTGGGGCTGTCACGATACCGCATGA
>Passalora_sequoiae_9LC2
MKPTSISFFALLAVAMARHKGNKPGEIGVFCFKGIKCYAWDLNNWSWQKCKSVGGHALLDIHFEHEFDPKSVSGMTSICQCAHGDTEEQRYIPITDQ
VTDYTFGWAYLTFGLPEKHGCQRTGAQTP*
ATGAGCAAGTCTCTGCTACGCGGCTTGTTGTCATATACTAAGCGCTGTCTAGAACCTACCTCTATTTCCTTCTTTGCGCTGCTTGCCGTCGCTATGG
CTAGGCATAAAGGAAATAAGCCGGGTGAAATCGGCGTGTTCTGCTTCAAGGGCATCAAATGTTATGCTTGGGACCTCAATAACTGGTCGTGGCAAAA
ATGCAAAAGCGTCGGCGGACACGCATTGCTCGATATTCACTTCGAACACGAGTTTGACCCGAAATCGGTTTCGGGAATGACAAGCATCTGCCAGTGC
GCACATGGTGATACCGAAGAGCAACGTTACATCCCTATTACCGATCAGGTCACGGACTACACATTTGGCTGGGCTTATCTGACGTTCGGGCTTCCTG
AAAAGCATGGATGTCAAAGGACGGGCGCACAGACCCCTTAG
>Passalora_sequoiae_9LC2
MRLAILSALFTILTLTTAKRGRNKPAKIDFTCYPPNGSKILLRCDEAHIETAIKKHCSDLGGDQLYNDLVNIRTDAGQVTGADAICLCAHGNAVEAK
WYDNETDPPGEVWVHFGLGTTFGVCRKNG*
ATGAGACTCGCCATCCTCTCGGCCCTCTTCACGATCCTGACCCTCACCACCGCCAAACGCGGCCGCAACAAACCCGCCAAAATCGACTTCACATGCT
ATCCCCCCAACGGATCCAAAATCCTCCTCCGCTGCGACGAAGCCCACATCGAAACCGCCATCAAAAAACACTGCAGCGACCTCGGTGGCGACCAGCT
GTATAATGATTTGGTCAATATCAGGACTGATGCGGGACAAGTGACGGGGGCGGATGCCATCTGTTTATGTGCGCATGGGAATGCGGTTGAGGCGAAG
TGGTATGATAATGAGACGGATCCGCCGGGGGAGGTGTGGGTTCATTTTGGGCTGGGGACCACGTTTGGAGTTTGTCGTAAGAATGGGTAG
>Teratosphaeria_gauchensis_CMW17545
MKLSTFPVLLGLLSIATEAKHGKNKPCKIEFTCDNTVQKSAAGCNTEQMQAWACTGIGAHDLAENGTISDGERNIGYSALCICAHGNTKNHDYVIDQ
GWEKGTATLRCGPPDTSGWYTCASEG*
ATGAGTACGCACAGCCATCACTCAGATGCTCGGCGTCGTTCTAACATCAAGCCCACAGAGCTCTCGACCTTTCCAGTCCTGCTTGGCCTCTTGTCTA
TCGCGACAGAGGCCAAGCACGGCAAGAACAAACCATGCAAGATTGAATTCACATGCGATAACACAGTGCAAAAATCAGCAGCAGGCTGCAACACCGA
GCAAATGCAAGCCTGGGTTACACAAGTCAGTCCAATTCTCACACGGCGATCCTTGGACCTCACTCTTTGACTGACTTCGCACTGCAATACCTCAGGC
TTGCACCGGCATAGGTGCTCACGACCTTGCGGAGAACGGGACCATTAGTGATGGTGAGAGGAACATTGGCTACTCAGCGCTCTGCATATGCGCGCAC
GGGAACACCAAGAATCATGACTATGTCATCGACCAGGGGTGGGAGAAAGGTACGGCCACATTGCGATGCGGTCCGCCTGACACCAGCGGATGGTACA
CATGTGCCTCGGAGGGTTGA
>Teratosphaeria_viscida_CMW51324
MRLSILPIILGLFFLSATAKHGKNKPCNIAFVCDPQIQLAKKCDEQKMTDWVTEACKGIGADSVALAGLTYDLKLNPVGYHAVCKCAHGNTKDDDYV
ISDDSGQWTKGTAMLRCSAPDVTGRFGCHQKA*
ATGAGTACGCACACCTGTGCTCAGCCGCTGTTCATTTATCTAACATCTCATTGACAGGGCTCTCAATCCTTCCGATCATCCTCGGACTCTTCTTCCT
GTCGGCGACGGCCAAGCATGGCAAGAACAAGCCATGCAACATAGCCTTCGTGTGCGATCCTCAAATACAACTGGCGAAAAAATGCGACGAGCAGAAG
ATGACAGACTGGGTTACTGAGGTTAGCATGCCTGACAGTCGCAAAGCTTCCTTCATAAATCACGAAGGACCAACTGATCAATCCGCTTTCGAGGCTT
GCAAGGGCATCGGCGCCGATTCCGTGGCATTGGCCGGTCTCACGTACGATCTGAAGCTCAATCCAGTAGGCTATCACGCGGTGTGTAAGTGCGCACA
CGGCAACACGAAGGACGACGACTACGTCATTAGCGATGACTCGGGGCAATGGACAAAGGGTACCGCGATGTTGAGGTGCAGTGCCCCGGATGTTACC
GGGCGTTTTGGCTGTCATCAAAAGGCATGA
>Teratosphaeria_destructans_CMW45661MRLSILPIILGLFFLSATAKHGKNKPCNIEFVCDPQVQLAKKCDEQKMTDWVNQACKGIGA
DSVAFAGLTYDMKLNPIGYHAVCKCAHGNTKDDDYVISDDTGQWTKGTAMLRCSAPDVIGQFGCHQRA*

ATGAGTACGCACACCTGTGCTCAGCCGCTGTCCATCTATCTAACATCTCATTGACAGGGCTCTCAATCCTTCCGATCATCCTCGGACTCTTCTTCCT
GTCGGCGACGGCCAAGCATGGCAAGAACAAGCCATGCAACATAGAATTCGTGTGCGATCCTCAAGTACAACTGGCGAAAAAATGCGACGAGCAGAAG
ATGACAGACTGGGTTAATCAGGTTAGCATGCCCGACAGTCGCAAAGCTTCCTTCATGAATCACGAAGGACCAACTGATCAATCCGCTTTCTAGGCTT
GCAAGGGCATCGGCGCCGATTCCGTTGCATTCGCCGGTCTCACGTACGATATGAAGCTCAATCCAATAGGCTATCACGCGGTGTGTAAGTGCGCACA
CGGCAACACGAAGGACGACGACTACGTCATTAGCGATGACACGGGGCAATGGACAAAGGGTACCGCGATGTTGAGGTGCAGTGCCCCGGATGTTATC
GGGCAATTTGGCTGTCATCAAAGGGCATGA

>Teratosphaeria_destructans_CMW45982

MRLSILPIILGLFFLSATAKHGKNKPCNIEFVCDPQVQLAKKCDEQKMTDWVNQACKGIGADSVAFAGLTYDMKLNPIGYHAVCKCAHGNTKDDDYV
ISDDTGQWTKGTAMLRCSAPDVIGQFGCHQRA*

ATGAGTACGCACACCTGTGCTCAGCCGCTGTCCATCTATCTAACATCTCATTGACAGGGCTCTCAATCCTTCCGATCATCCTCGGACTCTTCTTCCT
GTCGGCGACGGCCAAGCATGGCAAGAACAAGCCATGCAACATAGAATTCGTGTGCGATCCTCAAGTACAACTGGCGAAAAAATGCGACGAGCAGAAG
ATGACAGACTGGGTTAATCAGGTTAGCATGCCCGACAGTCGCAAAGCTTCCTTCATGAATCACGAAGGACCAACTGATCAATCCGCTTTCTAGGCTT
GCAAGGGCATCGGCGCCGATTCCGTTGCATTCGCCGGTCTCACGTACGATATGAAGCTCAATCCAATAGGCTATCACGCGGTGTGTAAGTGCGCACA
CGGCAACACGAAGGACGACGACTACGTCATTAGCGATGACACGGGGCAATGGACAAAGGGTACCGCGATGTTGAGGTGCAGTGCCCCGGATGTTATC
GGGCAATTTGGCTGTCATCAAAGGGCATGA

>Teratosphaeria_destructans_CMW44962

MRLSILPIILGLFFLSATAKHGKNKPCNIEFVCDPQVQLAKKCDEQKMTDWVNQACKGIGADSVAFAGLTYDMKLNPIGYHAVCKCAHGNTKDDDYV
ISDDTGQWTKGTAMLRCSAPDVIGQFGCHQRA*

ATGAGTACGCACACCTGTGCTCAGCCGCTGTCCATCTATCTAACATCTCATTGACAGGGCTCTCAATCCTTCCGATCATCCTCGGACTCTTCTTCCT
GTCGGCGACGGCCAAGCATGGCAAGAACAAGCCATGCAACATAGAATTCGTGTGCGATCCTCAAGTACAACTGGCGAAAAAATGCGACGAGCAGAAG
ATGACAGACTGGGTTAATCAGGTTAGCATGCCCGACAGTCGCAAAGCTTCCTTCATGAATCACGAAGGACCAACTGATCAATCCGCTTTCTAGGCTT
GCAAGGGCATCGGCGCCGATTCCGTTGCATTCGCCGGTCTCACGTACGATATGAAGCTCAATCCAATAGGCTATCACGCGGTGTGTAAGTGCGCACA
CGGCAACACGAAGGACGACGACTACGTCATTAGCGATGACACGGGGCAATGGACAAAGGGTACCGCGATGTTGAGGTGCAGTGCCCCGGATGTTATC
GGGCAATTTGGCTGTCATCAAAGGGCATGA

>Passalora_sequoiae_9LC2

MKITNIPFLALIASATALFHKHGDNKAGKLIYICDKNDDIHDCNPGDMLYWLEGKCWQIGGHQTINEKFKRDSNNKINGLELICLCAHGDTVQDSWR
IDGRVAPIGLVVLHFGLDTTFGVCHKNG*

ATGAGTGCGCATCCATTCCATGAGAGGATTCCACGACTGCATACTGACCTTCGAGCAGAGATAACCAACATCCCATTCCTAGCCCTCATCGCCTCCG
CCACAGCTCTCTTTCATAAGCACGGAGACAACAAAGCCGGCAAATTGATATACATCTGCGATAAGAACGACGACATACACGATTGCAACCCGGGCGA
TATGTTGTATTGGCTAGAGGGCAAGTGCTGGCAAATTGGCGGACACCAGACCATCAACGAGAAGTTCAAGCGTGACAGCAACAACAAGATCAACGGA
CTAGAACTTATCTGCTTGTGCGCGCACGGAGACACGGTACAGGATAGTTGGCGAATCGACGGGAGAGTTGCCCCAATTGGTTTGGTGGTGCTTCACT
TTGGGCTTGACACGACATTTGGGGTTTGTCATAAAAATGGATGA

>Passalora_sequoiae_9LC2

MKITIRPALLALLAATTVKGSGNKPGKIVATCHPPANSWNILNCDYDHLHTAIMKQCNKIGGDRLYHDHVQVTVDQNKNVRKGEAICLCAHGKTAQG
SWFDQDTIPPVSVDLSFGLGSTYNACQRNG*

ATGAAAATCACCATACGTCCAGCGCTCCTCGCACTCCTCGCCGCGACTACAGTTAAAGGAAGCGGCAACAAGCCCGGCAAAATCGTCGCTACTTGCC
ATCCCCCCGCCAATTCCTGGAACATACTCAACTGCGACTATGATCATTTACACACAGCCATCATGAAACAATGCAACAAAATCGGCGGGGACAGGTT
GTATCACGATCACGTCCAAGTGACCGTAGACCAGAACAAAAATGTCCGCAAAGGCGAAGCTATCTGCCTCTGCGCACACGGGAAAACCGCGCAGGGT
TCGTGGTTCGATCAGGATACTATACCGCCAGTCTCGGTGGATCTTTCGTTCGGACTTGGGTCGACGTACAATGCTTGTCAGAGGAATGGGTAG

>Teratosphaeria_nubilosa_CBS_116005

MRPSILPIILGLFYLSATAKHGKNKPCEIDFKCDPQIQAEKNCDMQKMMDWVHQACNGTGAASTELDSYWYDLKGNPTGFHAICTCAHGNTKNHDYV
ISGATGQWTPGTATLRCGPPAITSRYDCHRF*

ATGAGTACGCACACCTGTAATAAGCCGCTATTCATTGTTTTAACAACTCGTTGATAGGGCCCTCAATCCTTCCGATCATCCTCGGACTCTTCTACCT
GTCGGCGACGGCCAAGCATGGCAAGAACAAGCCATGCGAAATCGACTTCAAATGCGATCCTCAAATACAAGCGGAAAAAAACTGCGACATGCAAAAG
ATGATGGATTGGGTTCATCAGGTTAGCATGCCTAACAGTCGCAAAGCCTCCTTCACGGATCACGAACGACCTACTGAACCTGCCCACTTTTCAGGCT
TGCAATGGTACCGGCGCCGCTTCCACGGAATTAGATAGTTATTGGTACGATTTGAAGGGCAATCCAACAGGCTTTCACGCGATCTGTACGTGCGCAC
ATGGCAATACGAAAAATCACGACTACGTCATTAGCGGTGCCACAGGCCAATGGACGCCAGGTACCGCGACGTTGAGGTGCGGGCCTCCGGCCATCAC
ATCTCGTTACGATTGTCACAGATTCTAG

>Teratosphaeria_epicoccoides_CMW31933

MKTLILALLLALSPDASCKHGKHKPCSIEFSCDPVNRFSATGCNDTKMYEWDWQLCKGIGGYKWWGPGIVTNMTGGPVGWNMICECAHGNTKNRDFV
IESEGGQWEKGTATLRCGRPYIDAPYDCGRI*

ATGAGTATGCAAATCCGTTCCTAGACCACCCACCGACCGTTTGTCTAACCTTCCGTGTGCAGAGACCTTAATTCTCGCGCTCCTCCTCGCGCTGTCG
CCCGACGCATCATGCAAGCACGGCAAACACAAACCATGCTCAATTGAATTCTCGTGTGATCCTGTGAACCGATTTTCCGCGACGGGCTGCAATGATA
CCAAGATGTACGAATGGGATTGGCAGGTGAGTGGACCCAAAACGCACACATTCCATAGGCATGTCGACTCACTTTGGCCTCTATCTCCAGCTCTGTA
AAGGCATTGGTGGTTACAAGTGGTGGGGTCCTGGAATTGTGACCAACATGACTGGCGGCCCAGTTGGATGGAACATGATCTGTGAGTGCGCACACGG
GAACACCAAGAACCGCGACTTTGTCATTGAATCCGAGGGAGGGCAATGGGAGAAGGGCACTGCGACGTTGAGATGTGGTCGACCCTACATTGATGCG
CCTTATGATTGCGGGAGGATTTAA
